# Iterative computational design and crystallographic screening identifies potent inhibitors targeting the Nsp3 Macrodomain of SARS-CoV-2

**DOI:** 10.1101/2022.06.27.497816

**Authors:** Stefan Gahbauer, Galen J. Correy, Marion Schuller, Matteo P. Ferla, Yagmur Umay Doruk, Moira Rachman, Taiasean Wu, Morgan Diolaiti, Siyi Wang, R. Jeffrey Neitz, Daren Fearon, Dmytro Radchenko, Yurii Moroz, John J. Irwin, Adam R. Renslo, Jenny C. Taylor, Jason E. Gestwicki, Frank von Delft, Alan Ashworth, Ivan Ahel, Brian K. Shoichet, James S. Fraser

## Abstract

The nonstructural protein 3 (NSP3) of the severe acute respiratory syndrome coronavirus 2 (SARS-CoV-2) contains a conserved macrodomain enzyme (Mac1) that is critical for pathogenesis and lethality. While small molecule inhibitors of Mac1 have great therapeutic potential, at the outset of the COVID-19 pandemic there were no well-validated inhibitors for this protein nor, indeed, the macrodomain enzyme family, making this target a pharmacological orphan. Here, we report the structure-based discovery and development of several different chemical scaffolds exhibiting low- to sub-micromolar affinity for Mac1 through iterations of computer-aided design, structural characterization by ultra-high resolution protein crystallography, and binding evaluation. Potent scaffolds were designed with *in silico* fragment linkage and by ultra-large library docking of over 450 million molecules. Both techniques leverage the computational exploration of tangible chemical space and are applicable to other pharmacological orphans. Overall, 160 ligands in 119 different scaffolds were discovered, and 152 Mac1-ligand complex crystal structures were determined, typically to 1 Å resolution or better. Our analyses discovered selective and cell-permeable molecules, unexpected ligand-mediated protein dynamics within the active site, and key inhibitor motifs that will template future drug development against Mac1.

**Significance Statement:** SARS-CoV-2 encodes a viral macrodomain protein (Mac1) that hydrolyzes ribo-adenylate marks on viral proteins, disrupting the innate immune response to the virus. Catalytic mutations in the enzyme make the related SARS-1 virus less pathogenic and non-lethal in animals, suggesting that Mac1 will be a good antiviral target. However, no potent inhibitors of this protein class have been described, and pharmacologically the enzyme remains an orphan. Here, we computationally designed potent inhibitors of Mac1, determining 150 inhibitor-enzyme structures to ultra-high resolution by crystallography. In silico fragment linking and molecular docking of > 450 million virtual compounds led to inhibitors with submicromolar activity. These molecules may template future drug discovery efforts against this crucial but understudied viral target.

## Introduction

The macrodomain of SARS-CoV-2 NSP3 (Mac1) presents an intriguing target for drug discovery (1–5). Upon viral infection, host cells initiate an innate interferon-mediated immune response leading to the expression of poly-(ADP-ribose)-polymerases (PARPs), which catalyze the antiviral post-translational addition of ADP-ribose (ADPr) to a large range of target proteins (6). Mac1 enzymatically reverses this mono-ADP-ribosylation, counteracting immune signaling (7). Promisingly, inactivation of Mac1 by single-point mutations in the ADPr-binding site significantly reduced lethality and pathogenicity in mice after SARS-CoV infection (8). Small molecule inhibitors of SARS-CoV-2 Mac1 should therefore offer novel therapeutics to mitigate COVID-19 (9, 10).

A challenge for the development of such inhibitors has been the lack of small molecule modulators of macrodomain activity, other than ADPr; indeed, only recently have quantitative assays been developed (10, 11). This is true not only for Mac1 from SARS-CoV-2, but for the overall family of enzymes, which lack good chemical matter by which their activity can be probed, despite their importance in several areas of health and diseases. Accordingly, to map the recognition determinants of Mac1, we adopted a biophysical approach, screening for fragment ligands using protein crystallography, molecular docking, isothermal titration calorimetry (ITC), differential scanning fluorimetry (DSF) and a novel binding assay based on homogeneous time-resolved fluorescence (HTRF) (12). Mac1 proved to be unusually amenable to structure-determination, enabling us to determine the structures of over 230 fragment complexes, typically to ultra-high resolution (often better than 1.1 Å), affording us a detailed map of enzyme hot-spots with chemical matter of sufficient potency with which to optimize a quantitative assay (12, 13).

Nevertheless, our best fragments remained of modest potency, with none more potent than 180 μM. Here we describe the discovery and optimization of potent macrodomain ligands using two strategies (**Fig. 1**). In the first, we sought to link and merge pairs of fragments to create larger molecules that exploited multiple hot-spots, so reaching higher affinities. This used a new fragment-linking method (12, 14), adapted to explore a virtual library of 22 billion readily-synthesizable molecules (15). In a second approach, we exploited the hot-spots revealed by the initial fragments to guide computational docking of ultra-large chemical libraries of lead-like molecules, potentially more potent than the fragments docked in our original study (12). Both approaches ultimately led to compounds with IC_50_ values as low as 0.4 μM for the merged fragments and as low as 1.7 μM for the docking hits (**Fig. 1**). These represent the most potent inhibitors reported for any member of the broad family of macrodomains. Furthermore, the X-ray crystal structures that were determined for initial fragment-linking and docking hits as well as for molecules optimized for affinity or physical properties provide a comprehensive resource for drug development campaigns against this promising antiviral target.

**Figure 1.**
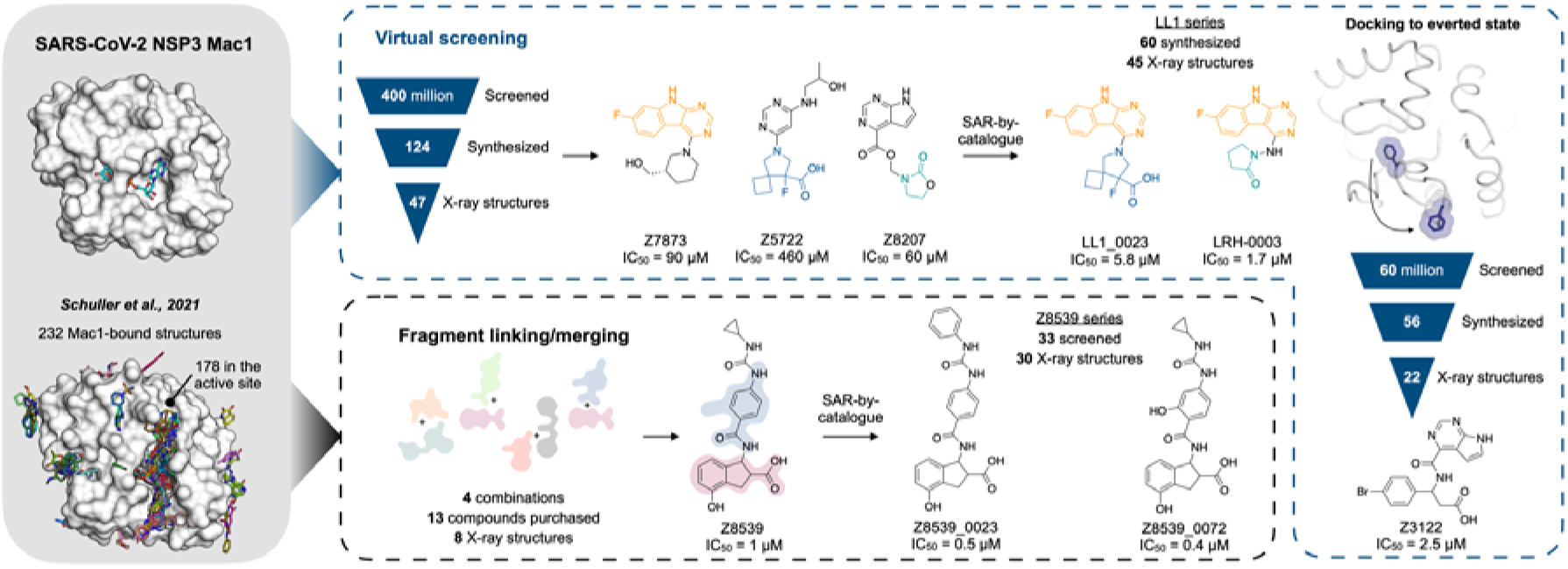
Overview of the structure-based strategies used to discover ligands that bind to the NSP3 macrodomain of SARS-CoV-2 (Mac1).

## Results

### Hit discovery through fragment-merging

The large collection of Mac1-fragment crystal structures revealed interaction patterns between initial ligands and the Mac1 active site (12). The largest subset of fragments bound in the adenine-recognition subsite, hydrogen-bonding to Asp22 and Ile23, and stacking with Phe156. Another group of mainly acidic fragments occupied a sub-pocket formed by the backbone NH groups of Phe156 and Asp157, which we labeled the “oxyanion subsite”. Although ADPr itself does not directly interact with this oxyanion site, the most potent compound that emerged from the fragment screen (ZINC263392672, PDB 5RSG, IC_50_ = 180 μM) placed a pyrrolo-pyrimidine group in the adenine subsite and carboxylate in the oxyanion subsite, suggesting that molecules able to bridge between both subsites hold potential for potent ligand design. An interactive dataset of the initial hits can be found at https://fragalysis.diamond.ac.uk/viewer/react/preview/target/Mac1.

Consequently, we sought to improve the affinity of the individual fragments by fusing pairs together into a larger, more potent molecule. Such fragment-linking has traditionally been considered technically difficult (16), as the linkage must minimally disturb the positioning of the two original fragments, and such a molecule must be synthetically accessible. Here, we tried to do so using a new automated fragment-linking approach, *Fragmenstein* (14), that searches purchasable chemical space to find molecules that could meet the design. From their crystallographic binding poses, fragments were merged based on superposed atoms or linked via hydrocarbon ethers. These virtually merged scaffolds were automatically modeled into the protein binding pocket by ensuring faithful placement of corresponding molecular segments onto the position of the original fragments (**Fig. 2**A,B). These virtually merged molecules became templates to search the make-on-demand chemical library of the Enamine REAL database, using the 2D molecular similarity search engine SmallWorld (http://sw.docking.org) and the substructure browser Arthor (http://arthor.docking.org) (17). We pursued four combinations of fragment hits to explore linked or merged scaffolds. Specifically, ZINC337835 (PDB 5RSW) was linked with ZINC922 (PDB 5RUE) (**Fig. 2** and **Fig. S1**) or ZINC98208711 (PDB 5RU5) (**Fig. S2**), ZINC26180281 (PDB 5RSF) was merged with ZINC89254160_N3 (PDB 5RSJ) (**Fig. S2**), and Z44592329 (PDB 5S2F) was merged with ZINC13514509 (PDB 5RTN) (**Fig. S2**). A total of 16 purchasable analogs (four for each linked or merged scaffold) were prioritized, of which 13 were successfully synthesized by Enamine. In subsequent crystal soaking experiments using the pan-dataset density analysis (PanDDA) algorithm to identify hits (18), 8/13 (~60%) bound to Mac1 (**Fig. 2**, **Fig. S1** and **Fig. S2**), seven had dose-responsive thermal up-shift of at least 0.5°C in DSF (**Dataset S1**), and two molecules had measurable binding to Mac1 in a HTRF-based ADPr-conjugated peptide displacement assay (**Fig. 2**).

**Figure 2.**
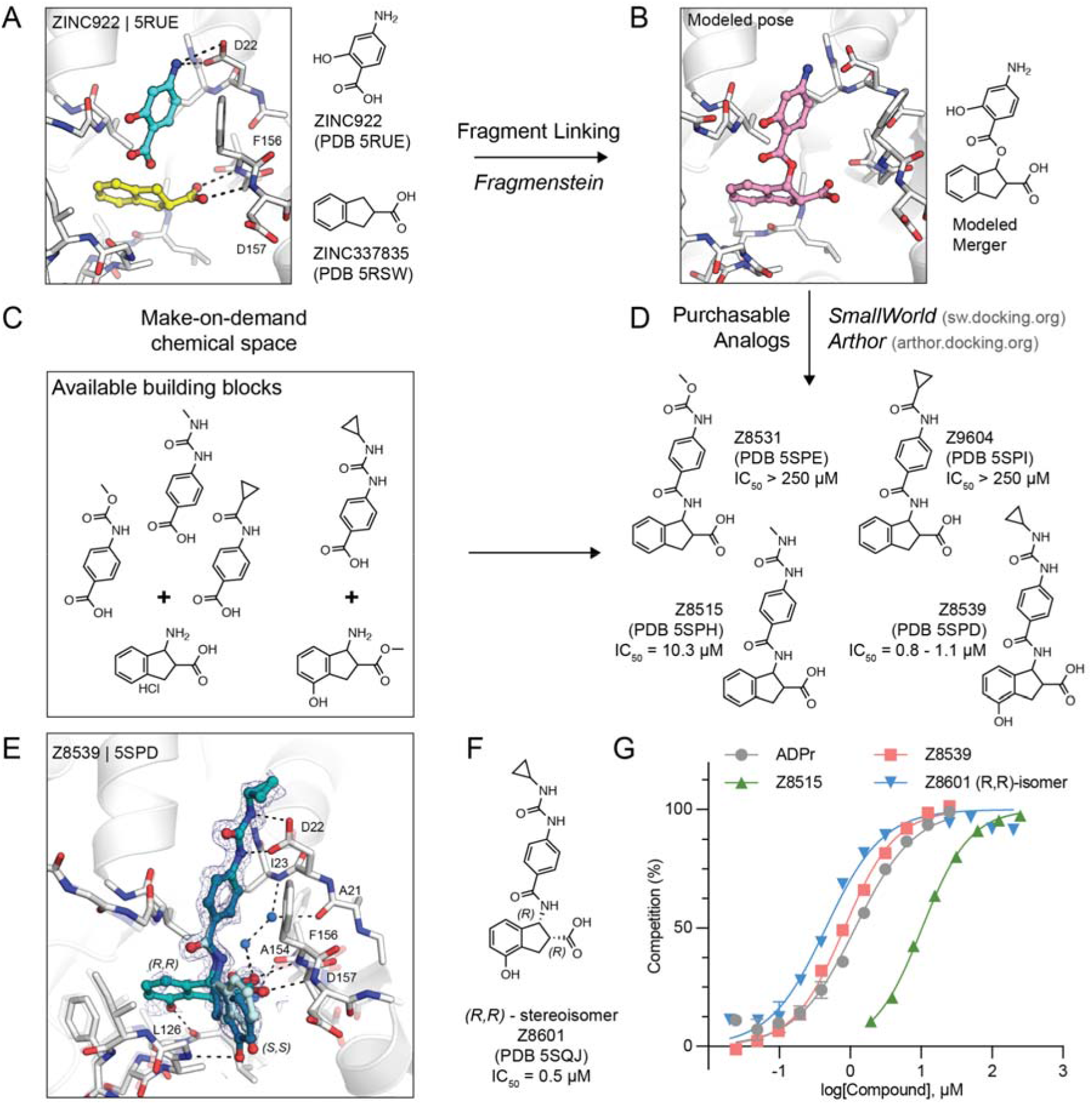
*In silico* fragment-linking targeting the adenosine site of Mac1. **A**) Binding pose of two fragments identified in the previously reported fragment screen (12). Fragment-protein hydrogen bonds are shown with dashed black lines. **B**) Theoretical linked scaffold of ZINC922 and ZINC337835 generated using *Fragmenstein (14*). **C**)Availability of corresponding chemical building blocks and reactions in the Enamine REAL database. **D**) Readily accessible analogs of the theoretical scaffold shown in **(B**). **E**) X-ray crystal structure of Mac1 bound to Z8539. Three conformations of Z8539 [(*R*,*R*) and two (*S,S*)] could be resolved in the PanDDA event map (blue mesh contoured at 2 σ). Water molecules that form bridging hydrogen bonds between Z8539 and the protein are shown as blue spheres. The apo state of Mac1 is shown with transparent white sticks. **F**) 2D structure of the most potent (*R,R*)-stereoisomer of Z8539 (Z8601). **G**) ADPr-peptide competition (%) of Z8539, Z8515 and Z8601 on Mac1 determined by an HTRF-based displacement assay. ADPr was used as reference. Data are presented as the mean ± SEM of at least two technical replicates.

### Identification of promising fragment merger

The linked scaffold combining the fragment hit ZINC922 (PDB 5RUE), occupying the adenine-recognizing subsite, with ZINC337835 (PDB 5RSW), placing a carboxylic acid at the oxyanion subsite, provided a promising template for a molecular scaffold bridging between both subsites (**Fig. 2**B). While the exact hypothetical merger was not readily available from the make-on-demand chemical space, we found four close analogs that were: Z4718398531 (Z8531), Z4574659604 (Z9604), Z4718398515 (Z8515), and Z4718398539 (Z8539) (**Dataset S1**). The main difference between these four accessible scaffolds and the initial merger model was the substitution of the fragment-linking ester by an amide, and the removal of the phenolic function of ZINC922 (**Fig. 2**D), both of which likely improve the *in vivo* stability of the molecules. The four analogs also differed in the substituents extending from the aniline amine, and Z8539 adds a hydroxyl-group to the indane of the initial fragment hit ZINC337835.

Remarkably, all four analogs were confirmed to bind Mac1 in crystallographic soaking experiments, with high fidelity between the predicted binding pose and the crystallographic result (**Fig. 2**E and **Fig. S1**). In the HTRF-based binding assay (12), Z8531 and Z9604 had IC_50_ values above 250 μM, while Z8515 and Z8539 had IC_50_ values of 7.9 μM and 0.8–1.1 μM, respectively. The more potent analogs both share a phenylurea group occupying the adenine subsite to stack with Phe156 and form bidentate hydrogen bonds between the urea and Asp22. Z8539 is among the most potent Mac1 compounds described with an affinity comparable to ADPr in the HTRF assay (0.9-1.3 μM) (**Fig. 2**G). The *K_D_* of the ADPr-conjugated peptide used in the HTRF assay was determined to be 2.7 μM by ITC (**Dataset S1**), therefore, the measured IC_50_ values of the molecules are similar to the binding affinities estimated using the Cheng-Prusoff equation (19).

All four molecules possess two chiral centers in the acid-bearing indane group, and initially the compounds were synthesized as diastereomeric mixtures, with evidence for at least two of the four diastereomers observed in the PanDDA event map for Z8539 (**Fig. 2**A). Chiral separation and testing of Z8539 confirmed that the (*R,R*) stereoisomer (Z8601), most faithful to the initial fragment hits, had the highest affinity for Mac1 with an IC_50_ of 0.5 μM, i.e. two-fold more potent than the diastereomeric mixture (**Fig. 2**F). In this configuration, the indane group partially inserts into the phosphate binding domain and the terminal phenol hydrogen-bonds to the backbone oxygen of Leu126. In the binding pose of the (*S,S*) stereoisomer (IC_50_ = 2.9 μM), the phenol is mainly solvent exposed and the hydroxyl hydrogen-bonds with the backbone nitrogen of Gly130 (**Fig. 3**A). By contrast, the two *trans* diastereomers showed reduced affinities with IC_50_ values between 43 and 55 μM. The X-ray crystal structure shows that the carboxylic acid of the (*R,S*)isomer only forms a single hydrogen bond to the oxyanion subsite (**Fig. S3**), while a structure of the (*S,R*) isomer was not obtained. The (*R,R*) stereoisomer (Z8601) was tested for off-target activity against two human macrodomains, MacroD2 and TARG1, using an adapted HTRF-based peptide displacement assay. The human proteins MacroD2 and TARG1 were chosen to test selectivity because they are the most similar human proteins to SARS-CoV-2 Mac1 (5). Z8601 showed no displacement of the ADPr-conjugated substrate at 50 μM against either target and approximately 50% displacement at high concentrations of 1 mM (**Fig. S4**). The selectivity of this scaffold for the viral over the tested human macrodomains is likely related to sequence differences within the ADPr-binding pockets between all three proteins: while Ala52 in the viral Mac1 offers ample space to accommodate the compound’s phenyl-urea functional group, MacroD2 and TARG1 carry considerably larger residues at the corresponding position, namely Leu50 and Cys104, respectively (**Fig. S4**).

**Figure 3.**
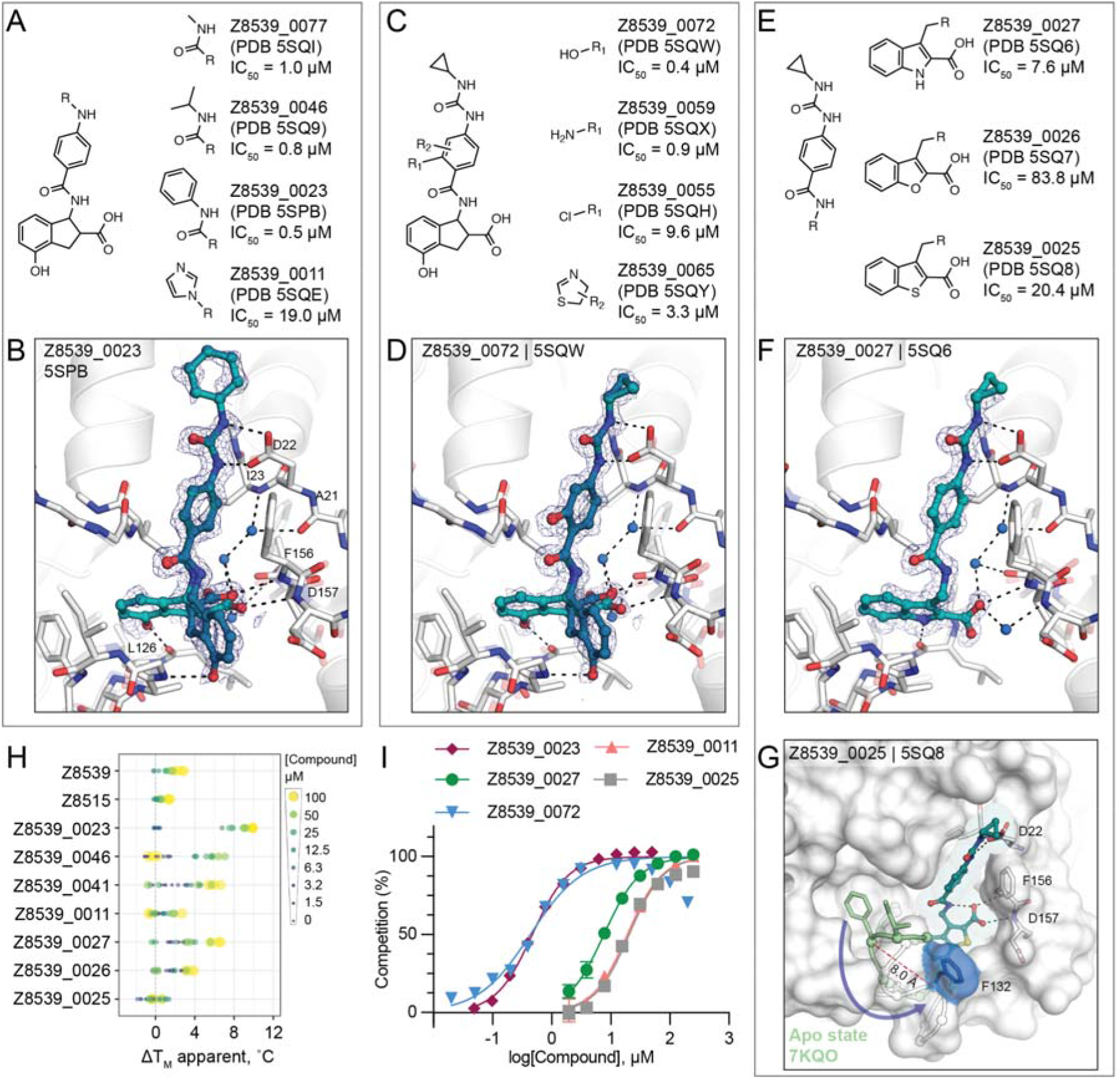
Structure-based optimization of Z8539. **A**) Modifications of the cyclopropyl-phenylurea group. **B**) X-ray crystal structure of Mac1 bound to Z8539_0023. The PanDDA event map is shown around the ligand (blue mesh contoured at 2 σ). **C**) Modifications of the central benzene. **D**) X-ray crystal structure of Mac1 bound to Z8539_0072. **E**)Modifications of the indane group. **F**) X-ray crystal structure of Mac1 bound to Z8539_0027. **G**) X-ray crystal structure of Mac1 bound to Z8539_0025. The Gly130-Phe132 loop is aligned to the apo-state conformation in green (PDB 7KQO). The Z8539_0025-Mac1 structure is shown with a transparent white surface. **H**) DSF-derived temperature upshifts. Data are presented for three technical replicates. **I**) HTRF-based peptide displacement dose-response curves. Data are presented as the mean ± SEM of at least two technical replicates.

The 1.05 Å resolution crystal structure of Mac1 in complex with the (*R,R*) isomer of Z8539 (Z8601) reveals an extended water-mediated hydrogen bond network between the ligand’s central amide, its carboxylic acid and Ile23, Ala21 as well as Ala154 (**Fig. 2**, **Fig. S3**). Interestingly, methylation of Z8539’s central amide group (Z8539_0056, **Fig. S3**) rendered the compound inactive, likely because of the interruption of this network. It is uncertain whether the initially generated ester-linked merger (**Fig. 2**B) can form this water network, and our preference for readily synthesized molecules may have conferred an unexpected advantage over the initial theoretical merger.

### Structure-based optimization of the merged scaffold

To further explore the Z8539 scaffold, we generated a structure-activity relationship (SAR) series (**Fig. 3**A-E). Here too, 2D-based similarity searches of the Enamine REAL database were used to find readily accessible and SAR-useful analogs, while analogs unavailable in the REAL database were also designed. Approximately 21,000 analogs (roughly 4,000 mono-anions) were identified via *SmallWorld* and subsequently docked against the Mac1-Z8539 crystal structure. Visual inspection of top-ranked (mostly) anionic compounds led to the selection of 19 readily accessible make-on-demand analogs, while nine compounds were manually designed; of these 28, 26 were successfully synthesized at Enamine. Of these 26 analogs, 23 were confirmed to bind Mac1 by crystallography and 20 showed activity in the HTRF assay (**Dataset S1**, **Dataset S2, Dataset S3**).

Most analogs bore modification of the cyclopropyl-phenylurea group of Z8539 (**Fig. 3**A). Removal of the cyclopropyl (Z8539_0041, PDB 5SPA) or replacement by either methyl (Z8539_0077, PDB 5SQI) or isobutyl (Z8539_0046, PDB 5SQ9) did not substantially change binding affinity, however, phenyl replacement (Z8539_0023, PDB 5SPB) improved the IC_50_ to 0.5 μM and showed a significantly increased thermal up-shift of 9°C in DSF (for the stereoisomeric mixture) (**Fig. 3**B,H). The resulting diphenyl-urea superimposes well with known fragment hits, e.g. Z44592329 (PDB 5S2F) or Z321318226 (PDB 5S2G) (12) (**Fig. S2**). Compound Z8539_0011 (IC_50_ = 19 μM) contains an imidazole moiety that forms an additional hydrogen bond to Lys55 (**Dataset S3**A.5). Addition of hydrogen bond donors such as amine (Z8539_0059, PDB 5SQX) or hydroxyl (Z8539_0072, PDB 5SQW) at the amide-ortho-position of the central benzene yielded relatively potent analogs with affinities of 0.9 μM and 0.4 μM, respectively (**Fig. 3**C,D). The corresponding crystal structures do not reveal additional interactions between the newly introduced substituents and the protein, however, the binding poses of the ligands indicate the formation of an internal hydrogen bond with the molecules’ central amides (**Fig. 3**D). Furthermore, the hydroxyl of Z8539_0072 formed a hydrogen bond with the backbone nitrogen of Lys11 of a symmetry mate, which closely matches the lattice interaction seen in the initial fragment hit ZINC922 (**Fig. S5**). Z8539_0072 did not show any off-target activity against either human TARG1 and MacroD2 at a concentration of 50 μM or 1 mM (**Fig. S4**), indicating selectivity for the viral Mac1 protein.

Finally, we tested analogs modulating the acid-carrying indane group (**Fig. 3**E). Of particular interest were achiral analogs where the indane was replaced by benzothiophene (Z8539_0025, PDB 5SQ8), benzofuran (Z8539_0026, PDB 5SQ7) or indole (Z8539_0027, PDB 5SQ6). The indole analog had low micromolar affinity (IC_50_ = 7.6 μM) for Mac1 and the crystal structure revealed a hydrogen bond between the indole amine and Leu126 (**Fig. 3**F). The lower affinity of this indole versus the parent compound may reflect the sub-optimal placement of the carboxylate in the oxyanion subsite. Surprisingly, although the benzothiophene (IC_50_ = 20 μM) and furan (IC_50_ = 84 μM) analogs only differ in one atom compared to the indole analog, the crystal structures in complex with Mac1 indicate that they adopt different poses, with a substantial rearrangement of the protein (**Fig. 3**G). The compounds’ cyclopropyl-phenylurea groups are shifted by 2.7 Å compared to the parent Z8539, while the benzothiophene or -furan groups are tilted by roughly 65° relative to the indole group in Z8539_0027, leaving the phosphate binding region vacant but enabling intramolecular hydrogen bonding between the carboxylic acid and the central amide. The loop formed by residues Ala129 to Pro136 adopts an everted conformation in which Phe132 is displaced by 8 Å and becomes almost fully solvent exposed, indicating high conformational flexibility in the phosphate binding region. Intriguingly, the displaced phenylalanine is reported to be crucial for catalytic function of macrodomains, e.g. mutation of Phe272 in human MacroD1 reduced enzymatic activity by approximately two-fold (20). This truly atomic structure-activity-relationship offers an unprecedented insight into the complex nature of protein-ligand interactions.

Although this compound series led to potent molecules, the Z8539 scaffold had low cell permeability (11 nm/s) in MDCK cells (**Fig. 4**A), which likely limits its potential antiviral activity. As carboxyl bioiosteres were not readily available for make-on-demand synthesis, we attempted to increase membrane permeability by replacing the cyclopropyl-phenylurea with a benzodiazol group, which only marginally reduced the IC_50_ value versus the parent urea (**Fig. 4**B). Z8539_0002 contains a methanol group that was designed to maintain the bidentate interaction with Asp22, however, the crystal structure instead indicated a hydrogen bond formed with the symmetry mate in the crystal lattice (**Fig. 4**C). Removing the alcohol group did not affect the binding affinity (Z8539_2001, **Fig. 4**D) and both compound analogs were selective for viral Mac1 over both tested human macrodomains (**Fig. S4**). However, despite lacking the urea, these compounds had similar P_app_ values compared to Z8539 (**Fig. 4**A), indicating that the carboxylate is most likely responsible for the observed low cell membrane permeability. Competitive substitutions of carboxylates for Mac1 inhibitors are presented at the end of the manuscript.

**Figure 4.**
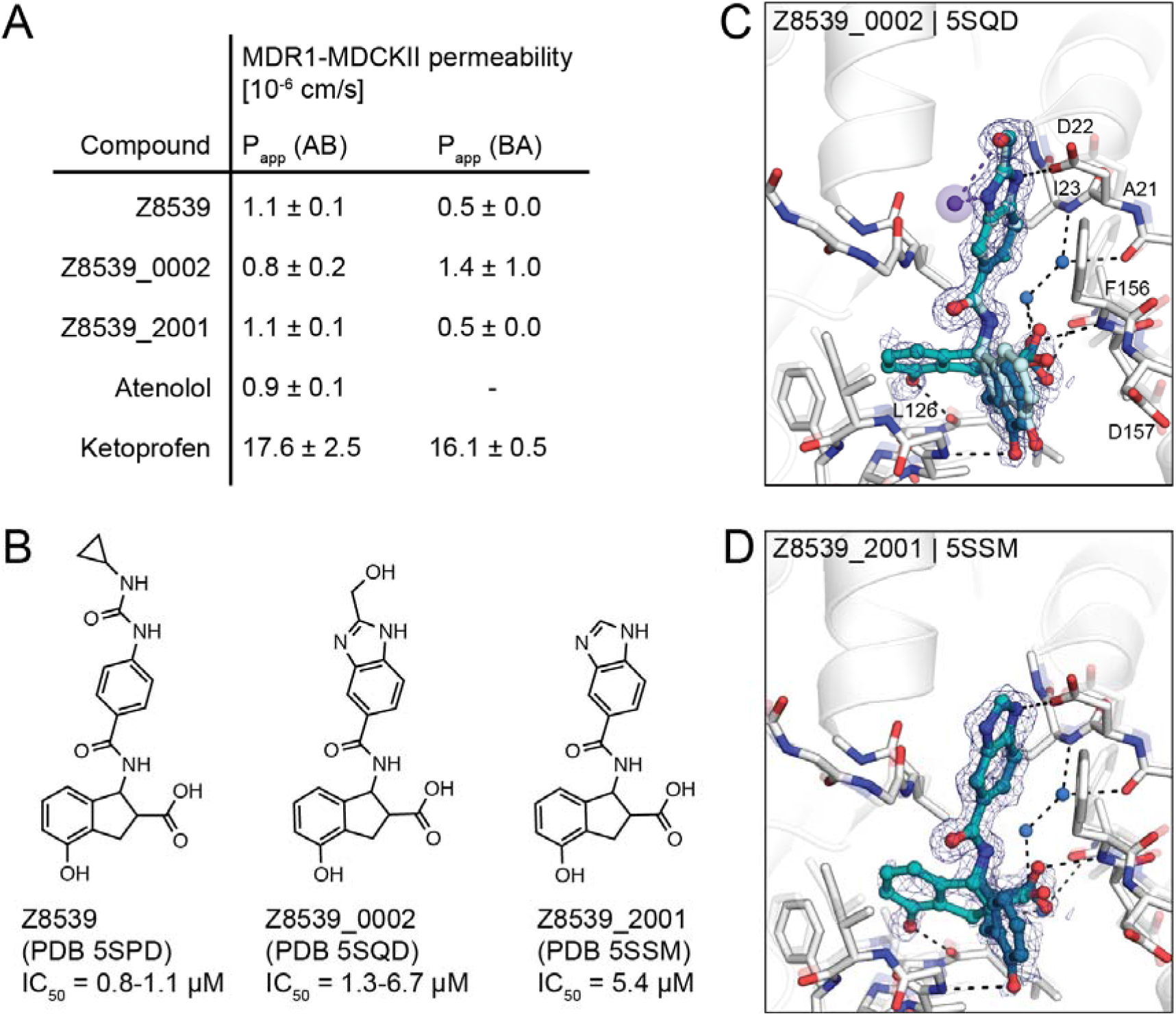
Z8539 analog with enhanced cell membrane permeability. **A**) Apparent permeability (P_app_) assayed with MDR1-MDCKII cells. Permeability was measured in apical (A)-to-basolateral (B) direction and vice versa. Atenolol and Ketoprofen were included as control compounds. **B**) 2D structures of Z8539, Z8539_0002 and Z8539_2001. **C**) X-ray crystal structure of Mac1 bound to Z8539_0002. Hydrogen bonding interactions between ligand and the Lys11 backbone nitrogen of a symmetry mate are shown with purple dashes/spheres. PanDDA event maps are shown around the ligand (blue mesh contoured at 2 σ). **D**) Crystal structure of Mac1 bound to Z8539_2001.

In summary, the new in silico fragment-linking approach employed here led to a promising and potent inhibitor scaffold based on only two fragments out of the roughly 200 fragment hits in the active site; many others remain to be considered. This method to explore the recent huge expansion of purchasable chemical space (21) may now allow the discovery of compounds that merge and minimally displace the key interactions of the parent fragments, which has previously limited fragment merging approaches. The combination of fragment-linking and large chemical library exploration might offer a pragmatic and relatively rapid strategy to generate active chemical matter for a vast group of protein targets with little to no known chemical matter.

### Novel inhibitors by molecular docking

Seeking even newer chemotypes, we docked ultra-large libraries of lead-like “tangible” (make-on-demand) molecules against Mac1 (21), leveraging the hotspots revealed by the initial fragment binding experiment (12). Molecules were screened against two different protein models, either using an ADPr-bound structure (PDB 6W02 (22)) or subsequently using a structure bound to a first-round lead-like docking hit (see below). The first screen of approximately 350 million molecules of the ZINC15 database (23), belonging mainly to libraries from Enamine and WuXi AppTec, with molecular weight ranging from 250 to 350 amu and calculated (c)logP below 3.5, was performed against the same docking template that we previously used in the computational fragment screen (ADPr-bound Mac1, PDB 6W02) (12). Molecules were targeted to the adenosine binding pocket of Mac1; molecules that docked to form polar interactions with the adenine-recognizing residues Asp22, Ile23 and Phe156, or with residues within the phosphate binding region such as Val49 or Ile132, were prioritized for experimental testing. Overall, 78 highly ranked molecules were selected for experimental testing, of which 22 (28%) were confirmed to bind Mac1 in crystallographic soaking screens, 11 (14%) showed binding in the HTRF assay at concentrations below 1 mM, and 30 (38%) revealed statistically significant thermal up-shifts of ≥0.5°C in DSF (**Dataset S1**).

In a second docking campaign, scoring parameters were optimized based on the results from the computational and crystallographic fragment screens as well as the first lead-like docking campaign (24). Here, the crystal structure of Mac1 in complex with Z6511 (PDB 5SOI, **Fig. 5**L) was used and the docking parameters were calibrated to ensure higher ranking of 172 previously confirmed fragment hits against a background of 2,384 molecules (mostly fragments) that did not bind to Mac1 in the crystal soaking experiments. Compared to the first docking model, this new screen better ranked acidic compounds interacting with the oxyanion subsite (Methods). Approximately 300 million compounds were docked, including ca. 250 million neutral and anionic compounds with molecular weights between 250 and 350 amu and clogP below 3.5 from the ZINC15 library (23), and 50 million compounds from in-house virtual anion libraries (with molecular weights between 250-400 amu) containing additional, mostly negatively charged molecules from the Enamine REAL database (15). From among the top-ranking molecules, 46 were obtained from Enamine, 25 (54%) of which were confirmed to bind Mac1 by X-ray crystallography, five (11%) showed activity in the HTRF binding assay at concentrations below 250 μM and eight (18%) were classified as hits in the DSF experiment (Methods).

**Figure 5.**
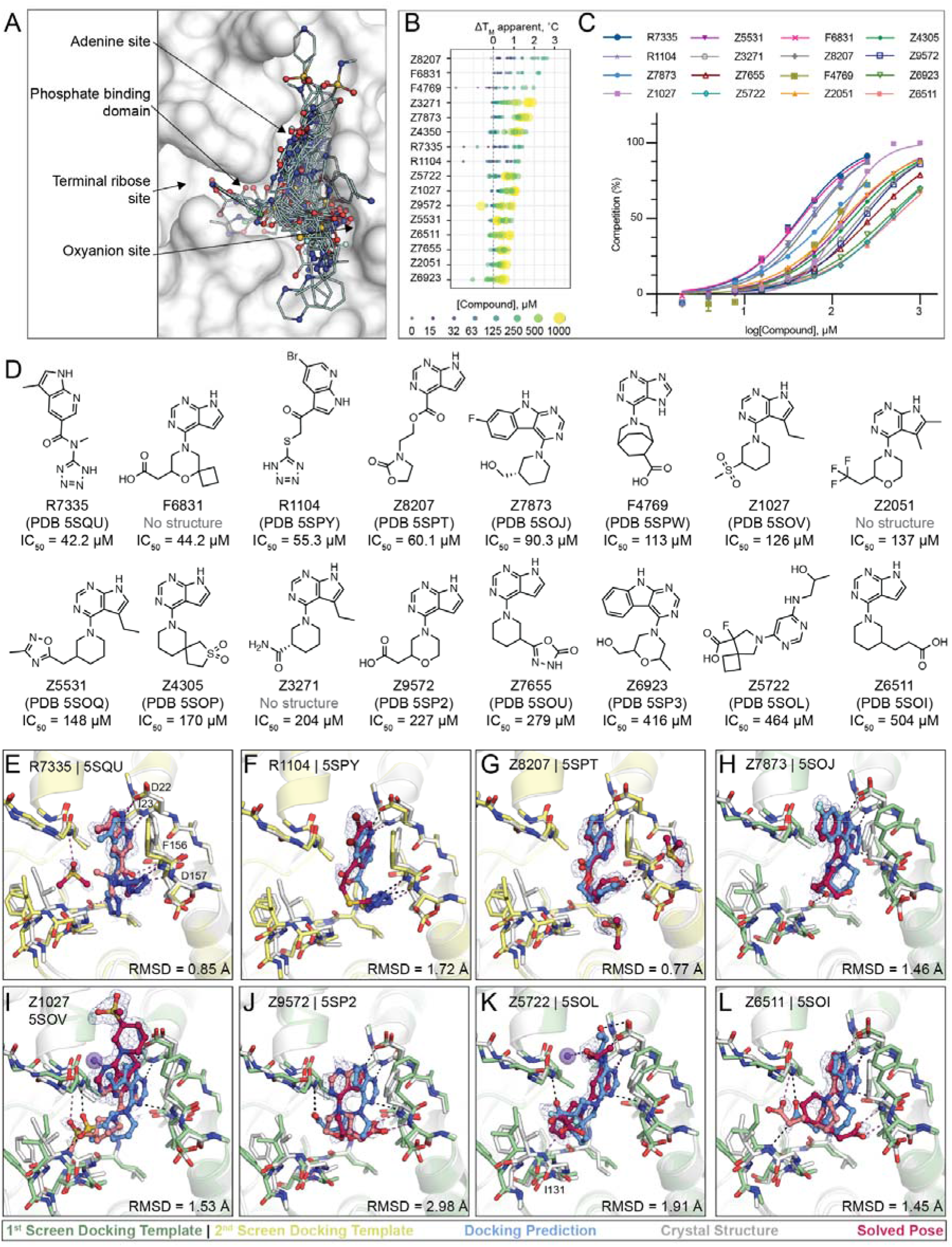
Large scale docking targeting the adenosine site of Mac1. **A**) Binding poses of 47 docking hits confirmed by X-ray crystallography. The ADPr-bound structure of Mac1 (PDB 6W02) is shown with a white surface. Thermal upshifts measured ibyn DSF. Data are presented for three technical replicates. **C**) HTRF-based peptide displacement dose-response curves. Data are presented as the mean ± SEM of at least 2 repeat measurements. **D**) 2D structures of docking hits with activity in the HTRF assay. **E**-**L**) Crystal structures of Mac1 bound to R7335, R1104, Z8207, Z7873, Z1027, Z9572, Z5722, Z6511, respectively. The protein structure used in the first docking screen is shown in green, the structure from the second screen is colored yellow. The predicted binding poses are shown in blue. Protein crystal structures are shown in gray and the solved binding poses are shown in red, with alternative ligand conformations colored salmon. Hydrogen bonding interactions between ligands and the Lys11 backbone nitrogen of a symmetry mate are shown with purple dashes/spheres. Hungarian Root Mean Square deviations (RMSD) between the docked and solved ligand poses were calculated with DOCK6. PanDDA event maps are shown for each ligand (contoured at 2 σ).

In summary, 124 molecules were selected from virtually screening more than 400 million distinct molecules in lead-like chemical space, finding 50 Mac1 ligands (40% hit rate) (**Fig. 5**). Of these, 47 were confirmed by crystallographic screening, and 13 showed measurable binding in the HTRF-based peptide displacement assay with IC_50_ values ranging from 42 to 504 μM. Only three molecules that showed ADPr-peptide competition in the HTRF assay were not confirmed by X-ray crystallography (F6831, Z2051, Z3271). The seemingly much higher hit-rate in the crystallographic soaking versus the HTRF-based peptide displacement experiments likely reflects the higher compound concentrations used in crystal soaking (10-20 mM) compared to the highest tested concentration in the HTRF-based assay e.g. 1 mM in the first docking campaign and 250 μM for the second campaign. Thirty-eight compounds showed significant thermal upshifts of more than 0.5°C in DSF (**Dataset S1**), thereby compounds with activity in the HTRF assay often had upshifts of >1°C. Ten compounds were confirmed by all three techniques.

### Docking hits explore the targeted adenosine binding pocket

Consistent with the docking predictions, almost all of the hits bound to the adenosine binding pocket in the Mac1 active site. A common structural motif among docking hits was a pyrimidine-containing headgroup that interacted with the adenine-recognizing residues of Mac1 (Asp22, Ile23, Ala154). Additional polar or even anionic moieties of docking hits typically bound in either the phosphate subsite or interacted with the oxyanion subsite (**Fig. 5**A). Two compounds, namely F9192 (PDB 5SPO) and Z4273 (PDB 5SPU) did not bind within the active site but occupied a shallow pocket near the terminal ribose binding site (**Fig. S6**, **Dataset S4**B.29, B.39). Although we previously identified several fragments binding in this site, they lack high quality interactions and are therefore unlikely to serve as starting points for ligands targeting this site. Good agreement between computationally predicted and crystallographically determined binding poses with Hungarian (symmetry corrected) root mean square deviations (RMSD) below 2 Å (25) was achieved for molecules with measurable binding affinity (e.g. R7335, R1104, Z8207, Z7873, **Fig. 5**D-G, **Dataset S1**), whereas larger deviations between docked and experimentally solved binding modes were observed for compounds with binding affinities outside of the tested range. For molecules predicted to place large, often cyclic moieties into the phosphate binding region, the corresponding crystal structures suggested binding modes extending from the adenine subsite to areas outside of the ADPr-binding active site, e.g. Z9710 (PDB 5SOK), Z8186 (PB 5SP1) or Z3280 (PDB 5SON) (see **Dataset S4**).

Although many different headgroups for the adenine subsite were explored among docking hits (see **Dataset S4**), molecules that were active in the peptide-displacement assay typically shared a pyrrolopyrimidine scaffold forming hydrogen bonds with Asp22, Ile23 and stacking with Phe156, e.g R7335 (PDB 5SQU), Z8207 (PDB 5SPT), Z6511 (PDB 5SOI) (see **Fig. 5**C, **Fig. 5**D,F,K). Two compounds, Z7837 (PDB 5SOJ) (**Fig. 5**G) and Z6923 (PDB 5SP3), extend the bicyclic purine headgroups into tricyclic pyrimidoindole scaffolds revealing moderate IC_50_ values of up to 90 μM, indicating favorable shape complementarity of larger segments in the adenine subsite compared to the nucleobase of ADPr. Of note, similar to what we observed in the fragment screen, four adenine-containing compounds (Z1211, Z4827, Z0893, Z0078) were not correctly synthesized and showed alkyl derivatives from the N3 rather than the intended N9 nitrogen in their corresponding crystal structures (see **Dataset S4**, **Dataset S1**) (12).

Among the most potent molecules were anions placing acidic functional groups such as a carboxylate (F6831, F4769, Z9572, **Fig. 5**C, **Fig. 5**J) or a tetrazole (R7335, R1104, **Fig. 5**E,F) in the oxyanion subsite. Interestingly, Z8207 (**Fig. 5**G) places oxazolidin-2-one, a polar but neutral functional group, in the oxyanion site, and has an IC_50_ of 60 μM. Ketone groups at the oxyanion site offer neutral alternatives to acid functional groups characteristic of many of the Mac1 inhibitors found to date (below). Two docking hits with measurable IC_50_ values inserted carboxylates into the phosphate binding region: Z5722 (IC_50_ = 464 μM, **Fig. 5**K) uses a rigid acid-carrying spiro-octane group to hydrogen bond with Ile131, while Z6511 (IC_50_ = 504 μM, **Fig. 5**L) projects a flexible butyrate side chain toward the oxyanion site.

### Ligand-mediated stabilization of alternative protein conformations

Surprisingly, in the crystal structures of three docking hits, namely Z4305 (PDB 5SOP, IC_50_ = 170 μM), F4769 (PDB 5SPW, IC_50_ = 113 μM) and Z5531 (PDB 5SOQ, IC_50_ = 148 μM) the compounds appear to stabilize alternative, open states of the phosphate binding region, wherein the loop formed by residues Leu127 to Pro136 adopts an everted conformation relative to the apo structure (**Fig. 6**A-C). Compared to the previously described structures of Mac1 bound to Z8539_0025 or Z8539_0026 (**Fig. 3G**), the docking hits induced even larger rearrangements within the active site. The magnitude of the loop rearrangement is surprising given the tightly packed Mac1 crystal lattice, formed prior to ligand soaking. All three compounds occupy the adenosine subpocket, forming hydrogen bonds between their pyrrolo-pyrimidine containing groups and Asp22 as well as Ile23. Z4305 and F4769 interact with the oxyanion subsite via sulfone or carboxylic acid, respectively (**Fig. 6**A,B). Both compounds stabilize the same loop rearrangement in which the Cα of Phe132 is displaced by 11 Å versus the canonical closed state, which does not seem able to accommodate the rigid and large non-aromatic cyclic moieties of the molecules, which would clash with Gly130. Z5331 stabilized a similar everted loop conformation (**Fig. 6**C). Whereas Z5331 does not interact with the oxyanion subsite, it inserts methyl-oxadiazole into the phosphate binding region, forming direct and water-mediated hydrogen bonds with Ser128 and Val49, respectively (**Fig. 6**C). As opposed to Z4305 and F4769, the central piperidine of Z5531 does not clash with Gly130, however, its methyl-oxadiazole would clash with

**Figure 6.**
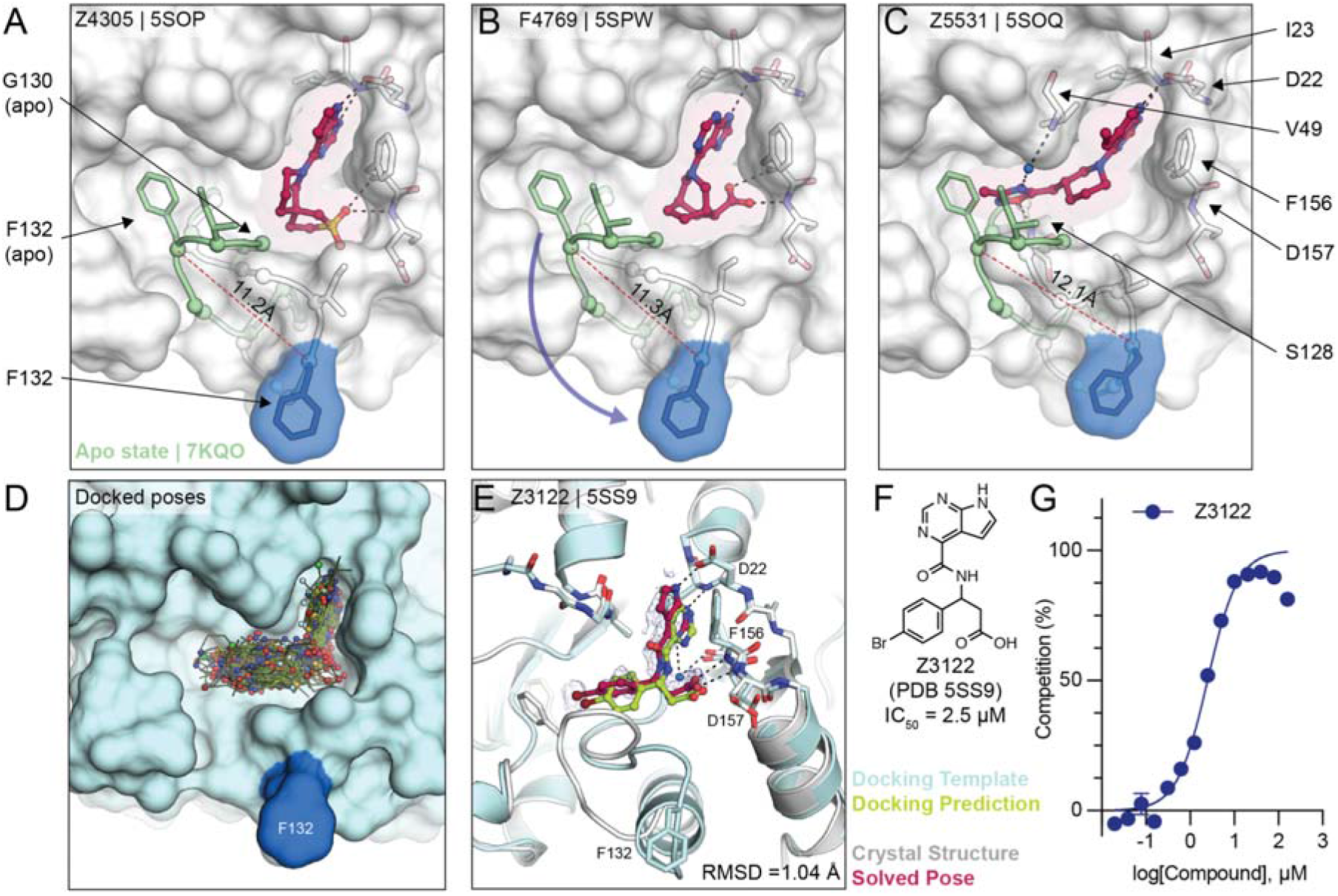
Stabilization of everted phosphate binding region by docking hits. **A,B,C**) The ligand-bound Mac1 crystal structures are shown in gray with Phe132 highlighted in blue. The Gly130-Phe132 loop of the Mac1 apo structure is depicted in green. Experimentally determined ligand-binding poses are shown in red. **D**) Predicted binding poses of molecules docked against the Z4305-bound Mac1 structure (PDB 5SOP). **E**) Crystal structure of Z3122 (red) bound to Mac1 (gray) compared to the predicted complex (Mac1 in blue, Z3122 in green). The PanDDA event map is shown around the ligand (blue mesh contoured at 2 σ). The Hungarian RMSD between solved and docked binding poses was calculated with DOCK6. **F**)Chemical structure of Z3122. **G**) HTRF-derived ADPr-peptide competition curve of Z3122. Data are presented as the mean ± SEM of three technical repeats.

Phe132 in the apo form. A similar conformational change in the Phe132-containing loop was observed for the merged fragment Z8580 (**Fig. S7**). The observed ligand-induced flexibility within the active site of Mac1 may hint to a catalytic mechanism requiring conformational flexibility to efficiently bind, cleave and release ADPr from different target proteins (13, 26).

### Docking to everted protein conformation

To investigate the potential ligandability of the everted Mac1 conformation, we virtually screened roughly 60 million anionic compounds of the ZINC22 virtual library (https://cartblanche22.docking.org) against the open state structure discovered in complex with Z4305. Ligands of this open state are predicted to bind with similar headgroups in the adenine site as closed state ligands, including polar interactions with Asp22 and Ile23, and stacking with Phe156. In addition, compared to the closed state, Ser128 was more solvent-exposed and was therefore targeted by molecules selected from this docking screen. Interactions with these three anchor points (Asp22, Ile23 and Ser128) were used to select molecules for experimental testing leading to a final set of 56 molecules that were synthesized by Enamine. On testing, 22 of these (39%) bound to Mac1 in crystal soaking experiments, of which five showed activity in the HTRF-based peptide displacement assay. While docking generated more favorable scores for the molecules against the open state than the closed state (**Dataset S1**), in the crystal structures all 22 hits bound to the closed state (see **Dataset S4**). Still, among the five in-solution hits, Z3122 (PDB 5SS9, see **Fig. 6**F) had an IC_50_ of 2.5 μM against Mac1 and had no measurable activity against the human macrodomains TARG1 or MacroD2 at 160 μM (**Fig. S8**), offering yet another promising, selective scaffold for future optimization.

### Structure-based optimization of docking hits

To improve the affinity of initial docking hits, we explored combinations of molecular substructures bound at different subsites, templated by their crystal structures. The fluoro-pyrimidoindole of Z7873 (PDB 5SOJ), occupying the adenine-subsite, was introduced into docking hits with mainly bicyclic purine scaffolds (e.g. Z9572, Z6511, Z5531) or combined with the spiro-octane-carboxylic acid of Z5722 (**Fig. 7A**). Nine analogs designed with this strategy were accessible in the Enamine REAL database and were synthesized for testing against Mac1. Of these nine, seven were confirmed to bind crystallographically, five were active in the DSF assay (see **Dataset S1**), and four bound in the HTRF assay (**Fig. S8**). Low micromolar affinities were measured for LL1_0023 (PDB 5SQO, IC_50_ = 6-10 μM) and LL1_0014 (PDB 5SQ3, IC_50_ = 16-29 μM), both containing the pyrimidoindole headgroup to occupy the adenine subsite and placing carboxylic acid in the phosphate binding region (**Fig. 7**B,D). Both compounds showed 50% displacement of the ADPr-conjugated peptide when tested against TARG1 and MacroD2 at 1 mM, whereas only LL1_0023 was active against TARG1 at 50 μM (**Fig. S6**). Compared to the Z8539 scaffold, LL1_0023 was 3-fold more permeable in MDR1-MDCKII cells.

**Figure 7.**
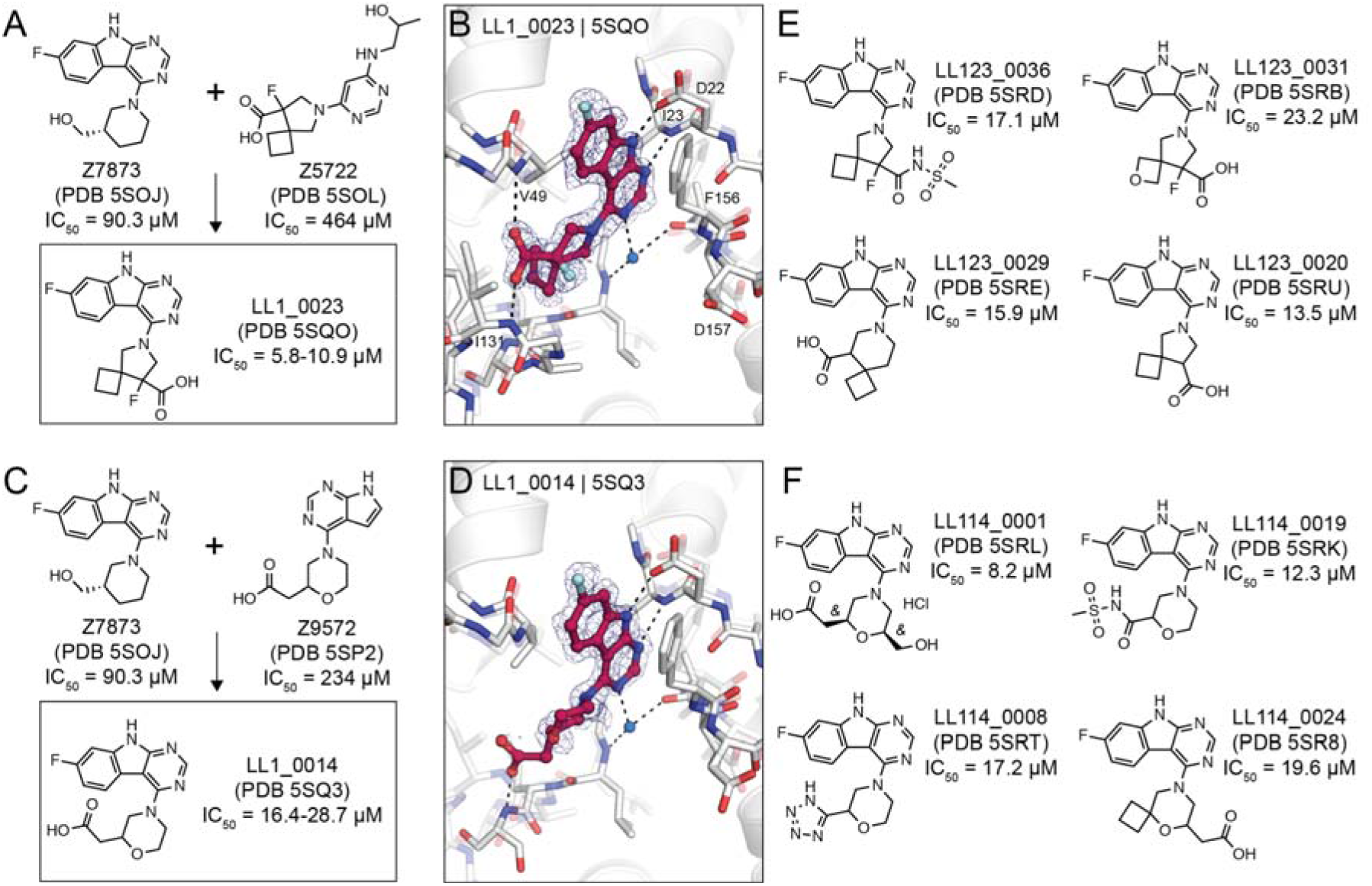
Structure-based optimization of docking hits. **A**) Design of LL1_0023. **B**) X-ray crystal structure of LL1_0023. The PanDDA event map is shown around the ligand (contoured at 2 σ). Hydrogen bonds are shown with dashed black lines. **C, D**) Design and X-ray crystal structure of LL1_0014, respectively. **E, F**) Selected analogs of LL1_0023 and LL1_0014, respectively.

Thirteen analogs of the LL1_0023 scaffold were selected and synthesized from the Enamine chemical space to investigate structure-activity-relationship for this scaffold. Eleven of these bound in crystal soaking experiments (**Dataset S5**), while nine analogs had IC_50_ values below 200 μM in the ADPr-peptide displacement assay (**Fig. S6**). No improvement of affinity was achieved by replacing the carboxylic acid of LL1_0023 by sulfonamide (LL123_0036, PDB 5SRD, IC_50_ = 17 μM, **Fig. 7**E), or replacing the cyclobutane with oxetane (LL123_0031, PDB 5SRB, IC_50_ = 23 μM). In addition, modifications of the compound’s core spiro-octane e.g. replacement by spiro-nonane (LL123_0029, PDB 5SRE, IC_50_ = 16 μM) or removal of a fluoro group (LL123_0020, PDB 5SRU, IC_50_ = 14 μM, **Fig. 7**E) did not change affinity notably. Correspondingly, removal or neutralization of the acidic functional group by methylation increased IC_50_ values to over 200 μM (**Dataset S5**).

For the LL1_0014 scaffold, 18 analogs were designed and synthesized by Enamine, 16 of which bound to Mac1 in the soaking or HTRF-based binding experiments. Here, addition of an ethanolic group to the central morpholino group, reflecting the initial docking hit Z7873 (**Fig. 7**C), improved the IC_50_ value to 8 μM (LL114_0001, PDB 5SRL, **Fig. 7**F). Similarly, the addition of cyclobutane to the morpholino group, which mimicked the docking hit F6831 (**Fig. 5**C), showed slight improvement of affinity (LL114_0024, PDB 5SR8, IC_50_ = 20 μM). Furthermore, exchanging the carboxylic acid by bioisosteres such as sulfonamide (LL114_0019, PDB 5SRK, IC_50_ = 12 μM) or tetrazole (LL114_0008, PDB 5SRT, IC_50_ = 17 μM) seemed to moderately improve the ligands’ binding affinities (**Fig. 7**F). In subsequent screens against TARG1 and MacroD2, only the tetrazole-containing analog (LL114_0008) showed measurable peptide displacement against TARG1 and MacroD2 at 50 μM (**Fig. S8**). Additional analogs are shown in the Supporting Information (see **Dataset S5**).

In summary, large-library docking and subsequent structure-based optimization revealed several potent inhibitors of Mac1, structurally unrelated to those obtained from fragment-linking. This expanded the number of low μM scaffolds, each topologically unrelated to the others, to at least five families of molecules inhibiting a key viral enzyme for which none had been previously known.

### Towards potent neutral Mac1 inhibitors

Although our initial SAR for Mac1 ligands showed the benefit of carboxylate binding to the oxyanion subsite, ADPr instead interacts with this subsite via a water-mediated hydrogen bond to a neutral ribose hydroxyl. The development of non-anionic inhibitors might hold several advantages for antiviral drug discovery, especially considering drugs will need to cross cell membranes to engage viral targets residing within infected host cells. To identify neutral alternatives to carboxylate and other anions at this site, we designed a small set of analogs by linking the previously identified pyrrolo-pyriminde or pyrimidoindole to small moieties bearing hydrogen bond donor or acceptor functionality (e.g., sulfones, hydroxyls, pyridines, or ketones) (see **Fig. 8**A, **Dataset S6**). A total of 124 molecules (290 enantiomers) were generated in 3D conformer libraries for computational docking (see Methods). We selected 21 compounds based on the predicted docking poses of which 20 were synthesized by Enamine. Fourteen of these 20 molecules (70%) were confirmed to bind to Mac1 by X-ray crystallography and four (20%) showed binding in the HTRF-based assay.

**Figure 8.**
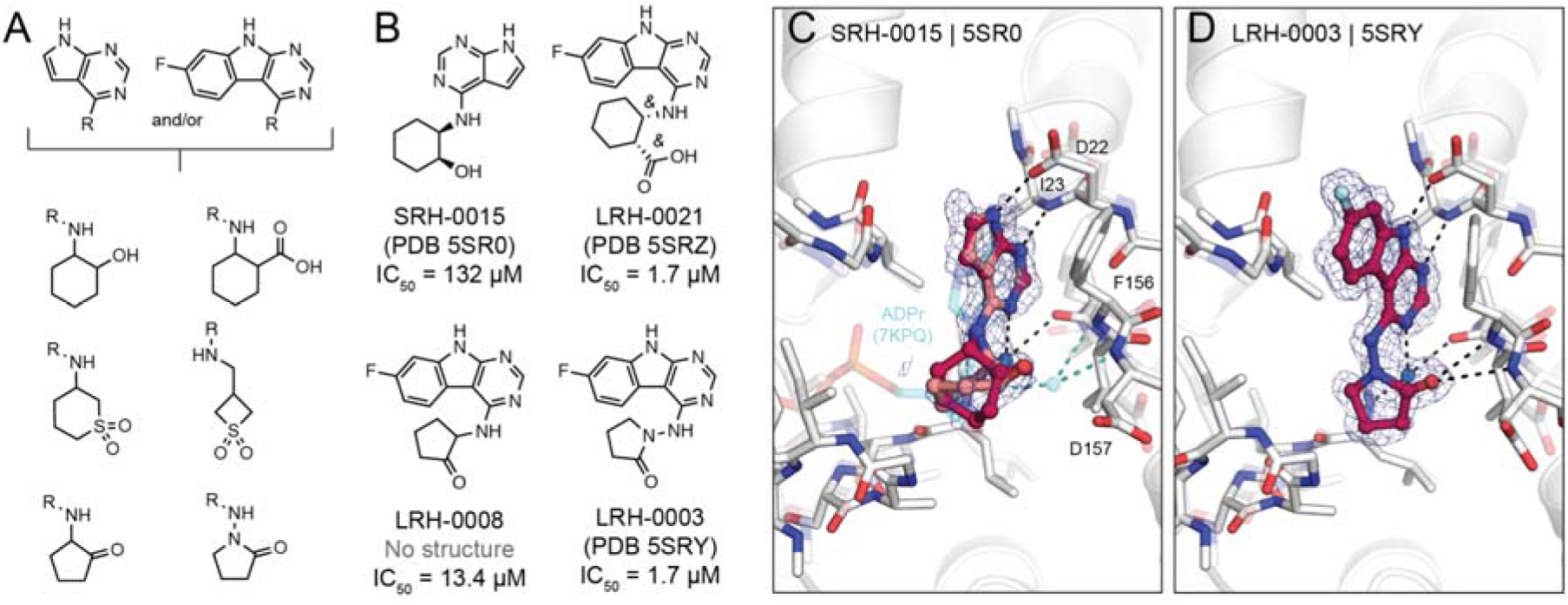
Probing neutral functional groups in the Mac1 oxyanion subsite. **A**) Design strategy of analog set. **B**) Chemical structures of most potent hits. **C**) Crystal structure of Mac1 bound to SRH-0015. ADPr and the water-mediated hydrogen bond to the oxyanion subsite are shown for reference (PDB 7KQP, transparent cyan sticks/spheres). Both of the *trans* stereoisomers were modeled: the (*S,R*) is colored dark red and the (*R,S*) isomer is colored salmon. PanDDA event maps are shown around the ligand (blue mesh contoured at 2 σ). **D**) Crystal structure of the Mac1-LRH-0003 complex.

Promisingly, SRH-0015 (PDB 5SR0, IC_50_ = 132 μM, MW = 232 amu), notably active for its small size, placed a hydroxyl group towards the oxyanion subsite, mimicking the placement of a ribose-hydroxyl group of ADPr (see **Fig. 8**B,C). While the crystal structure of Mac1 in complex with ADPr revealed a water-mediated hydrogen bond between the corresponding ADPr-hydroxyl and the oxyanion site, the structure of the Mac1-SRH-0015 complex does not suggest direct or water-mediated hydrogen bonding (**Fig. 8**C). The most promising analogs from this series were LRH-0008 (IC50 = 13.4 μM) and LRH-0003 (PDB 5SRY, IC50 = 1.7 μM) (see **Fig. 8**B,D). These compounds contain fluoro-pyrimidoindole headgroups joined to 2-aminocyclopentan-1-one or 1-aminopyrrolidin-2-one rings, respectively. The crystal structure of LRH-0003 bound to Mac1 revealed favorable placement of its hydrazide carbonyl function at the oxyanion site, enabling simultaneous hydrogen bonding to both NH groups of Phe156 as well as Asp157 (**Fig. 8**D). The enhanced potency of LRH-0003 versus LRH-0008 is consistent with a stronger hydrogen bonding interaction in the former, given the greater basicity of the hydrazide carbonyl present in LRH-0003 as compared to the ketone in LRH-0008. Notably, the similar anionic analog LRH-0021 (PDB 5SRZ, **Dataset S6**) was equipotent to LRH-0003 indicating that neutral compounds can indeed offer competitive alternatives to anionic Mac1 ligands. Encouragingly, both LRH-0003 and LRH-0021 obtained high permeability values in MDR1-MDCKII cell-based assays of 138 and 120 nm/s in apical to basal and 243 and 91 nm/s in basal to apical direction, respectively. Thereby, the carboxylate of LRH-0021 may form an internal hydrogen bond to the compound’s central amine group leading to improved membrane permeability. Although the anionic compound LRH-0021 showed binding to TARG1 at 160 μM, neither neutral compounds LRH-0003 and LRH-0008 had measurable binding to TARG1 and MacroD2 (**Fig. S8**), suggesting higher selectivity for the neutral isosteres.

## Discussion

Like many antiviral targets to emerge from SARS-CoV-2, Mac1 is both highly attractive and challenging. While animal studies in SARS have highlighted its crucial role in viral pathogenesis, there were no reliable chemical tools, or really inhibitors of any kind, for the enzyme. Fortunately, Mac1 crystallized readily and diffracted to ultra-high resolution (often better than 1 Å), supporting fragment-based exploration of its recognition determinants, both empirically and computationally (12). Capitalizing on this, over 230 fragment structures were determined. The binding poses of the ligands tiled the active site of the enzyme, but despite often favorable ligand efficiencies, none of the fragments had affinities more potent than 180 μM. Here, we built on the molecular determinants revealed by the fragment structures to discover potent, selective, and cell-permeable molecules, making progress towards chemical probes and leads for drug development.

Four key points emerge from this effort. First, an automated fragment merging and linking strategy, allied with searches of ultra-large libraries, identified molecules that combined key groups of pairs of fragments *and* were readily available from make-on-demand synthesis. This led to the rapid discovery of molecules with low μM affinity that were subsequently optimized to affinities as low as 430 nM (compound Z8539_0072), an overall improvement of >400-fold compared to the best starting fragment. Second, templated again by the ligand-recognition patterns revealed by the fragments, molecular docking screens found compounds with affinities down to 2.5 μM, with several in the mid-μM range that were also optimizable to the low μM. The best of these had ligand efficiencies that were measurably better than even the merged fragments. Third, while most of these molecules were anionic with high polar surface areas that reduced cell permeability, structure-based optimization found analogs with fewer hydrogen-donating groups like ureas, alcohols and phenols, and enabled the replacement of anionic warheads with neutral ones. This suggests that it may be possible to improve cell membrane permeability for several of the scaffold classes here. Finally, these efforts occurred against an understudied target from an enzyme family without validated chemical probes, hinting at the potential of structure-based approaches to advance chemical matter against other understudied proteins.

We used X-ray crystallography both as a primary screening tool to identify macrodomain-binding compounds from computational design, and to provide structural information to guide compound optimization. The success of this approach was partly due to the high-quality nature of the Mac1 crystals in the P4_3_ space group; they grew readily, withstood high concentrations of DMSO and diffracted consistently to <1 Å. The high resolution diffraction, coupled with analysis of electron density with PanDDA (18), allowed us to identify fragments with occupancies below 20% in the initial fragment screen (12). Low occupancy fragments included ZINC337835 and ZINC922, which were linked together in the present work to generate Z8539, a potent binder of Mac1 (**Fig. 2**), testifying to the potential of this approach. Although the initial fragments were soaked at high concentrations (10 mM), only hints of fragment binding were visible in F_O_-F_C_ difference maps, and the fragment binding signal was largely obscured by ground-state solvent (**Fig. S9**). However, both ZINC337835 and ZINC922 could only be modeled unambiguously into PanDDA event maps (**Fig. S9**). This contradicts recent arguments that no useful conclusions can be derived from ligands modeled at the low occupancies detected by PanDDA (27). Our work, and that of others (28, 29), shows how low-occupancy ligands can inspire the design of more potent analogs. In addition to identifying the fragments that led to Z8539, PanDDA helped to identify the most potent stereoisomer of Z8539. We initially obtained this compound as a mixture of diastereomers, and although the density indicated that the major isomer was (*S,S*), inspection of the PanDDA event map at low contour level hinted that the (*R,R*) isomer might be present (**Fig. S9**). This prompted us to test the four diastereomers separately, which revealed that the (*R,R*) isomer was the most potent in solution, with good agreement between the fragments modeled using PanDDA and the theoretical model (**Fig. S1**).

One notable complication to using X-ray crystallography to screen ligands is the influence of crystal lattice interactions on ligand binding (30). Our initial fragment screen revealed that the P4_3_ crystal form had a substantially higher hit rate compared to the C2 crystal form (24% versus 6%) (12). We partly attributed the difference in hit rates to fortuitous crystal packing in the P4_3_ form: the backbone nitrogen of Lys11 on a symmetry mate is ideally positioned to interact with compounds binding in the adenine subsite. Indeed, 66 of the 123 fragments identified in or near the adenine subsite formed hydrogen bonds with Lys11 (12). Similarly, in the present work, several of the compounds that were identified by virtual screening, and subsequent optimization, adopted alternative conformations that were stabilized by hydrogen bonds with Lys11 (e.g. Z1027, Z9020, LL123_0006 and LL123_0016,). Although one might be tempted to discard these conformations as artifacts, our current work indicates that they can be useful. One of the two fragments that were linked to create Z8539 contained a hydroxyl that formed a hydrogen bond with Lys11 (**Fig. S5**). The compound lacking the hydroxyl (4-aminobenzoic acid, ZINC920) did not bind to Mac1 in the fragment screen (12). Crystal lattice interaction may explain the large difference been predicted and observed binding mode for several of the hits from virtual screening (e.g. F9046, F0346, R3575, Z6744, Z6684, Z5740, Z6689, Z6567).

We were surprised to find several ligands that induced large scale re-arrangement of the active site loop consisting of residues 127-136 (**Fig. 6**, **Fig. S7**). Conformational changes involving Ala129, Phe132 and Asn99 have been characterized in this loop in the ADPr-bound state (12) and in the ligand-free enzyme at low pH (13), but these are relatively minor compared to the 7-12 Å shifts in Phe132 seen here. Everted loop conformations have also been observed for other macrodomains, including human MacroD1 (PDB 2X47) (31) and PARP14 (PDB 5O2D) (32) (**Fig. S7**). Despite the apparent flexibility of this region, our initial virtual screening campaign did not identify any compounds that stabilized the flipped conformation of Ala129 that is present in the ADPr-bound state, despite using this state as a template for docking (PDB 6W02) (**Fig. 5**). However, during compound optimization, several structures were determined with Ala129 in the flipped state. These included LL114_0041, which places a carboxylic acid in the phosphate binding subsite, and LL123_0020, which stabilizes a water molecule in a similar position (**Fig. S6**). A similar rearrangement in water networks was seen for the docking hit Z0828, although the shift in Ala129 was smaller (**Fig. S6**). These ligands offer new opportunities for structure-guided design efforts targeting the phosphate binding subsite of Mac1.

Certain caveats merit discussion. The anti-viral or immunomodulating effect of the developed compounds has not been shown. This partly reflects limitations of the molecules themselves–e.g., their current low cell permeability–but it also reflects the lack of suitable cell-based assays to monitor the effect of Mac1 inhibition on interferon signaling. The development of such assays is an urgent need in the field; currently, our only way to measure the efficacy of Mac1 inhibitors, outside of the enzyme itself, is *in vivo*. On a technical level, while hit rates of computational docking were high in the X-ray soaking assay, only a few truly potent compounds were identified in the HTRF-based binding assay. Furthermore, while many docking predicted poses corresponded well to the crystallographically determined poses, compared to the previous fragment docking screen, larger deviations between docked and crystallographic poses were sometimes observed, especially among molecules that were predicted to insert deep into the phosphate-binding pocket. Also, ligand-induced stabilization of alternate conformations of the mobile active site loop was not predicted. While docking against a Mac1 structure with the everted Phe132 loop conformation (PDB 5SOP) led to a potent 2.5 μM inhibitor (Z3122), the Mac1-Z3122 crystal structure showed binding in the closed state (**Fig. 6**E). In addition to shortcomings of computational docking, our fragment-linking strategy relied on the access to chemicals mimicking theoretically linked scaffolds. In our case, the purchasable analogs offered promising templates, however, some differed noticeably from the initial model e.g. they replaced a central hydrogen bond acceptor (ester group) with a donor (amide group). Although this exchange seemed actually beneficial in our Z8539-series, similar changes might lead to loss of activity in other cases.

These caveats do not diminish the central observations of this study. From an initial mapping of the Mac1 binding site with >230 fragment crystal structures (12), fragment-linking and -merging led to compounds that bound >400-fold better than the best fragment. The same mapping identified hot spots that supported ultra-large library docking that identified mid- and low μM binders falling into still newer families. Overall, the determination of 150 new Mac1-ligand crystal structures supported the discovery and optimization of 19 low- and sub-μM compounds falling into eight different scaffolds and chemotypes, while another 28 compounds in eleven scaffolds were discovered in the 10 to 50 μM range. While these compounds retain permeability liabilities, structure-based optimization suggests routes to improving their physical properties, including by reducing hydrogen-bond donors and swapping anionic for neutral warheads, without substantial loss of affinity for the enzyme. From a technical standpoint, the rich of structure-activity-relationships combined with X-ray crystal structures for most compounds described here creates a dataset for benchmarking and improving computational techniques for drug discovery, such as free energy perturbation (33, 34). From a therapeutic perspective, the compounds and structures described in this study will support progress towards first-in-class antiviral therapeutics targeting the NSP3 macrodomain of SARS-CoV-2.

## Materials and Methods

### Fragment merging/linking

Fragment mergers and linkers were generated using *Fragmenstein (14)*. Specifically, spatially superposed atoms or rings are combined, while attempting to maintain bonding, and separate fragments are linked, depending on distance, via a bond, oxygen bridge or hydrocarbon ether bridge. The resulting compounds are corrected for any defects, such as impossible valence, and minimized under strong constraints using PyRosetta. The merging and the search for purchasable similar compounds was performed similarly to the example Colab notebook for *Fragmenstein (14)*. The structure PDB 6WOJ (7) was chosen as a template structure and was energy minimized with 15 cycles of FastRelax in PyRosetta restrained against the electron density map and with ADPr parameterised. The initial fragments were processed and merged pairwise. The mergers that were predicted with a combined RMSD less than 1 Å were sorted by Rosetta-predicted binding Gibbs free energy and the top mergers were manually inspected. The SmallWorld server was queried for purchasable compounds similar to the top merged compounds (17), which were then placed restrained to the initial fragments.

### Computational docking

Docking calculations were performed with DOCK3.7 (24, 35) using precomputed scoring grids for rapid evaluation of docked molecules. Scoring grids for van der Waals interactions were generated with CHEMGRID and electrostatic potentials within the targeted binding pocket were calculated by numerical solution of the Poisson-Boltzmann equation with QNIFFT (36). Therefore, AMBER united-atom charges (37) were assigned to the protein and selected structural water molecules. Ligand desolvation scoring grids were computed using Solvmap (38).

In the first docking screen, the crystal structure of SARS-CoV-2 NSP3 Mac1 bound to ADPr (PDB 6W02 (22)) was used as a template for docking. All water molecules except for HOH324, HOH344, HOH383 and HOH406 as well as chain B were removed. Next, the Mac1-ADPr complex with selected water molecules was prepared for docking following the protein prepwizard protocol of Maestro (Schrödinger v. 2019-3) (39). Accordingly, Epik was used to add protons and protonation states were optimized with PROPKA at pH 7 (40). The complex was energetically minimized using the OPLS3e force field. Thereby, the maximum heavy-atom root-mean-square deviation from the initial crystal structure was 0.3 Å. The atomic coordinates of the adenosine substructure within the co-crystallized ADPr molecule were used to generate 45 matching spheres for placement of ligand atoms by the docking program (24). For the calculation of the binding pocket electrostatic potential, the dielectric boundary between the low dielectric protein environment and high dielectric solvent was moved outwards from the protein surface by 1.9 Å using spheres generated by Sphgen. In addition, partial atomic charges of backbone amide hydrogen atoms of residues Ile23 and Phe156 were increased by 0.2 elementary charge units (e) while partial charges of the corresponding backbone carbonyl oxygen atoms were reduced by the same amount, hence, retaining the residues’ net charges. Furthermore, the dielectric boundary was extended by 0.4 Å from the protein surface for the generation of ligand desolvation scoring grids (24). At the time we launched the first lead-like docking screen against Mac1, ADPr was the only known ligand of the enzyme. Consequently, we calibrated the docking parameters according to their ability to place and score adenosine, adenine and ribose within the adenosine-binding site against a background of 250 property-matched decoys generated with the DUDE-Z approach (41). In addition, an Extrema set was screened to ensure prioritization of mono-anions and neutral molecules (24).

A total of 330,324,265 molecules with molecular weights ranging from 250 to 350 amu and calculated (c)logP below 3.5 from the ZINC15 lead-like library were screened (23). In total, 316,505,043 compounds were successfully scored, each exploring on average 3,111 orientations and 405 conformations leading to the evaluation of roughly 175 trillion complexes in 65,794 core hours or roughly 66 hours on a 1000-core cluster. The predicted poses of the top-scored 500,000 molecules were filtered for internal molecular strain (total strain <6.5 TEU; maximum single torsion strain <1.8 TEU (42)) and their ability to form hydrogen bonds to residues Asp22, Ile23, Gly48, Val49, Gly130 or Phe156. Molecules with unsatisfied hydrogen bond donors or more than three unsatisfied acceptors were deprioritized (43). Finally, 90 molecules were purchased from Enamine, of which 78 (87%) were successfully synthesized.

For the second docking campaign, the crystal structure of Mac1 in complex with the first-round docking hit ZINC000078036511 (Z6511, PDB 5SOI) was used as the structural template. Chain B and all water residues were removed and the Z6511-Mac1 complex (using conformation B of the ligand) was prepared according to the protein prepwizard protocol using Maestro (see above) (39). During scoring grid preparation, the low dielectric protein environment was extended by 1.8 Å outwards from the protein surface. In addition, the partial atomic charge of the backbone amide hydrogen atom of Ile23 was increased by 0.4 e whereas the partial charges of the backbone amide hydrogen atoms of Phe156 and Asp157 were increased by 0.2 e without modulating the net charge of the residues. Forty five matching spheres for ligand placement by docking were generated based on atomic coordinates obtained from various first-round lead-like docking hits as well as previously described fragments: ZINC000078036511 (PDB 5SOI), ZINC000292637864 (PDB 5SOT), ZINC901381520 (PDB 5S6W), ZINC57162 (PDB 5RV3), ZINC26180281 (PDB 5RSF) and ZINC336438345 (PDB 5RSE) (12). The described docking parameters were evaluated by control calculations ensuring the enrichment of 142 previously identified fragment ligands and 24 first round lead-like docking hits against a background of 2,384 experimentally determined non-binders (2,333 fragments, 51 lead-like molecules).

Using the ZINC15 database, 246,246,485 neutral and monoanionic molecules from the lead-like set were docked against this Mac1 model, resulting in the scoring of 156 trillion complexes where each scored molecule was on average sampled in 3,431 orientations and 428 conformations within 63 hours on a 1000-core computer cluster. In addition, an in-house anion library containing (mostly) negatively charged molecules with molecular weight between 250 and 400 amu from the 22B Enamine REAL database was screened. In total, 39 million anions were identified by performing SMART pattern searches in RDKit (www.rdkit.org) of carboxylic acid and 33 bioisosteres. In the docking screen, 37,556,136 molecules were scored, each sampled in 4,134 orientations and 343 conformations on average resulting in the evaluation of 19.5 trillion complexes in approximately 20 hours on a 1000-core computer cluster. A final set of ca. 16 million mostly anionic molecules from the February-2020 release of Enamine REAL was docked against Mac1. Within 10,703 core hours, 15,957,174 molecules were scored by evaluating a total of 12 trillion complexes where each molecule sampled on average 5,142 orientations and 495 conformations.

The top 1 million scored compounds from each screen were investigated for intramolecular strain (total strain <7.5 TEU, maximum single torsion strain <2.5 TEU (42)) and hydrogen bonding with Asp22, Ile23, Gly48, Val49, Phe156 and Asp157. Molecules with unsatisfied hydrogen bond donors or more than three unsatisfied acceptors were not considered for experimental evaluation. The second docking campaign led to 54 molecules that we selected for synthesis at Enamine of which 46 (85%) were obtained. The small analog set designed to probe neutral alternatives of negatively moieties binding in the oxyanion subsite were docked using the parameters from the second large-scale docking campaign. Molecules were protonated using ChemAxon Jchem 2019.15 (https://chemaxon.com/) at pH 7.4, rendered into 3D with Corina (v.3.6.0026, Molecular Networks GmbH, https://mn-am.com/products/corina/) and conformational libraries were generated with Omega (v.2.5.1.4, OpenEye Scientific Software; https://www.eyesopen.com/omega).

A third docking screen was performed against the Z4305-stabilized, everted conformation of Mac1 (PDB 5SOP). Before docking to the open structure, MDMix (44) (that utilizes AMBER18 (45)) was performed to assess binding hotspots in this less explored state. For this, the protein was solvated in pre-equilibrated mixtures of 20% ethanol and water, as well as 20% methanol and water. Three replicates of 50 ns simulations (six simulations total) were performed. Settings for minimization, equilibration and the production phase were set to default (44). After the simulation, all trajectories in the three independent simulations for each solvent mixture were aligned, after which the observed density was converted to binding free energies using the inverse Boltzmann relationship. Low energy regions were visualized and inferred to be probable binding hotspots.

The crystal structure of Mac1 in complex with Z4305 was prepared for docking following the same steps as above, i.e. protonation, minimization and grid preparation. The dielectric boundary between the low dielectric protein environment and high dielectric solvent was moved outwards from the protein surface by 1.9 Å. Forty-five matching spheres were generated based on 26 atomic coordinates of Z4305 and Z5531 as well as 19 randomly placed spheres covering the oxyanion subsite and the surface near Ser128. Partial atomic charges of backbone amide hydrogen atoms of residues Ile23 were increased by 0.4 elementary charge units, while backbone amide hydrogen atoms of residues Phe156, Asp157 and Ser128 were increased by 0.2 elementary charge units. Partial charges of the corresponding backbone carbonyl oxygen atoms were reduced by the same amount. The described docking parameters were evaluated by control calculations the same way as described above for the second docking screen.

Using a new virtual library, ZINC22 (https://cartblanche22.docking.org), a collection of 60,732,663 monoanionic lead-like compounds (heavy atom count 17 to 25) were screened. Within 40 hours on a 1000-core computer cluster, roughly 57 million compounds were scored, each sampled in approx. 5,336 orientations and 359 conformations resulting in more than 54 trillion complexes. The molecules that reached a total score threshold of −35 kcal/mol (comprising 4.3 M molecules) were filtered for internal molecular strain (<6.5 TEU; maximum single torsion strain <1.8 TEU) after which 1.7 M molecules remained. Next, molecules with more than one unsatisfied hydrogen bond donor or more than three unsatisfied acceptors were removed. Four independent sets were clustered by similarity for visual inspection, namely compounds able to interact with i) Asp22, Asp157 and Ser128, ii) Asp22, Phe156 and Ser128, iii) Ile23, Phe156 and Ser128, and iv) Asp22, Ile23 and Ser128, which led to 2, 249, 1761 and 2,249 compounds, respectively, ultimately leading to 70 being purchased from Enamine, of which 56 (80%) could be synthesized.

### Crystallization and ligand soaking

Crystals of SARS-CoV-2 NSP3 Mac1 were grown using an expression construct that crystallized in the P4_3_ space group, as described previously (12) (**Dataset S2**). This construct crystallizes with two molecules in the asymmetric unit: the active site of protomer A is accessible to ligands while the active site of protomer B is obstructed by a crystal lattice interaction (12). The P4_3_ crystal system was chosen because the crystals grow readily, diffract to atomic resolution and tolerate soaks in 10% DMSO for at least 6 hours (12). Briefly, crystals were grown by microseeding in 96-well sitting drop plates (SWISSCI, 3W96T-UVP), using 30 μl of 28% PEG 3000 and 100 mM CHES pH 9.5 in the reservoir and crystallization drops containing 100 nl seeds, 100 nl reservoir and 200 nl protein (40 mg/ml in 150 mM NaCl, 20 mM Tris pH 8.5, 5% glycerol and 2 mM DTT). Crystals grew to maximum size in ~24 hours at 19°C. Compounds were prepared in DMSO to 100 mM, or to the maximum concentration allowed by solubility (see **Dataset S2** for compound concentrations). Compounds in DMSO were added to crystallization drops using acoustic dispensing with an Echo 650 liquid handler (Labcyte) (46). Soaks were performed with either 40 or 80 nl of compound per crystallization drop, giving a nominal concentration of 10 or 20 mM (see **Dataset S2**). After incubating for 2-4.5 hours at room temperature, crystals were vitrified in liquid nitrogen using a Nanuq cryocooling device (Mitegen). No additional cryoprotectant was added prior to vitrification. Although there was no observed decrease in diffraction quality with increased soak time (**Fig. S9**), certain compounds (namely Z8601 and LRH-0003) induced substantial disintegration of crystals after two hours, possibly linked to disruption of the crystal lattice by binding of compounds to the protomer B active site (12). Despite the crystal disintegration, reflections were recorded to <1 Å for crystals soaked with both compounds (**Dataset S2**).

### X-ray diffraction data collection and data reduction

Diffraction datasets were collected at beamline 8.3.1 at the Advanced Light Source, beamlines 12-1 and 12-2 at the Stanford Synchrotron Radiation Lightsource, or beamline 17-ID-2 at the National Synchrotron Light Source II. The data collection parameters used at each beamline are listed in **Dataset S2**. X-ray diffraction images were indexed, integrated and scaled with XDS (47), using a reference P4_3_ dataset to ensure consistent indexing. The high resolution limit for each dataset was chosen based on a CC_1/2_ value of ~0.3 in the highest resolution shell (48). The diffraction resolution of crystals frequently exceeded the maximum resolution achievable with the experimental set-up; for these datasets, the high resolution limit was set to achieve ~95% completeness in the highest resolution shell. Data were merged with Aimless (49), and free R flags were copied from a reference P4_3_ dataset. Structure factors intensities for all datasets have been uploaded to Zenodo in MTZ format (DOI: 10.5281/zenodo.6856943). For some compounds, datasets were collected from multiple crystals. Data collection and reduction statistics for all datasets summarized are in **Dataset S2**.

### Ligand identification, modeling and refinement

All datasets were initially refined with the Dimple pipeline (50) run through CCP4 (51) using a starting model refined from a crystal soaked only in DMSO (dataset UCSF-P0110 in **Dataset S2**). Ligands were identified using PanDDA version 0.2.14 (18), with a ground-state map calculated using 34 datasets collected from crystals soaked only in DMSO. PanDDA was run an additional two times with ground-state maps calculated using 35 or 62 datasets from the ligand-soaked crystals where no ligands were detected. This procedure led to the identification of an additional 19 binding events, four of which were not identified in the first PanDDA run. Datasets used for ground-state map calculation for each of the PanDDA runs are annotated in **Dataset S2**. For ligands with multiple crystals/datasets, only the highest occupancy event was modeled. Ligands were modeled into PanDDA event maps using COOT version 0.8.9.2 (52) with ligand restraints generated using *phenix.elbow* (53) or ACEDRG (54) from a SMILES strings, or from coordinates generated using LigPrep version 2022-1 (55). Based on the background density correction (BDC) values, ligand occupancies ranged from ~10-90% (**Dataset S2**). Many of the ligands had multiple conformations and/or isomers present. The isomers modeled, and the estimated ratios based on PanDDA event maps, are listed in **Dataset S2**. Datasets were collected from soaks performed with two batches of Z8539_0002; one of the datasets was modeled with the (*R,R*) and (*S,S*) isomers (PDB 5SQD), while the other was only modeled with the (*R,S*) isomer (PDB 5SSN). Two compounds, LRH-0022 (PDB 5SRH) and LRH-0031 (PDB 5SRI), were only modeled with their pyrimido-indole core.

For all ligands, we modeled changes in protein structure and water in the ligand binding sites into PanDDA event maps. Alternative conformations were included for residues where the heavy-atom RMSD value of the ligand-bound model to the ground-state model was greater than 0.15 Å. This cut-off was chosen with reference to the RMSD values for the 34 ground-state structures, where 99.7% of residues had RMSD values <0.15 Å (**Fig. S9**). In these multi-conformer models, the ground-state model was assigned the alternative occupancy identifier (altloc) A and the ligand-bound state was assigned altloc B (and C/D when overlapping conformations/isomers were present). Water molecules modeled into PanDDA event maps were assigned altloc B, and ground-state water molecules were included within 2.5 Å of ligand-bound state ligands or water (assigned altloc A).

Refinement of the ligand-bound multi-conformer models was performed with *phenix.refine* using five refinement macrocycles (56). Coordinates and atomic displacement parameters (ADPs) were refined for all protein heavy atoms, and hydrogens were refined using a riding model. Based on previous observations (18), the occupancy of the ligand-bound and ground-states were set to 2*(1-BDC) and 1-2*(1-BDC) respectively, and occupancy refinement was switched off. Water molecules were automatically added to peaks in the mF_O_-DF_C_ difference density map >3.5 σ using *phenix.refine*. To prevent the multi-conformer water molecules being removed by the automatic solvent picking, the ligand- and ground-state waters were renamed from HOH to WWW. After one round of refinement, maps and coordinates were inspected, and additional water molecules were placed manually using COOT into peaks in the mF_O_-DF_C_ difference map. Based on positive/negative peaks in the mF_O_-DF_C_ difference maps after refinement, the occupancies for some ligands were adjusted (initial and adjusted occupancies are listed in **Dataset S2**). Next, a second round of refinement was performed with ADPs refined anisotropically for non-hydrogen atoms, with automatic water picking, and the refinement of water coordinates, switched off. Data refinement statistics are summarized in **Dataset S2**. Coordinates, structure factor intensities and PanDDA event maps for all datasets have been deposited in the Protein Data Bank under the group deposition IDs G_1002236, G_1002238 and G_1002239. Additionally, the PanDDA input and output files have been uploaded to Zenodo (DOI: 10.5281/zenodo.6856943).

### Homogeneous Time Resolved Fluorescence assay

Binding of the compounds to macrodomain proteins was assessed by the displacement of an ADPr conjugated biotin peptide from His_6_-tagged protein using a HTRF-technology based screening assay which was performed as previously described (12). The expression sequences used for SARS-CoV-2 Mac1, and the human macrodomains TARG1 and MacroD2, are listed in **Dataset S2**. All proteins were expressed and purified as described previously for SARS-CoV-2 Mac1 (12). Compounds were dispensed into ProxiPlate-384 Plus (PerkinElmer) assay plates using an Echo 525 liquid handler (Labcyte). Binding assays were conducted in a final volume of 16 μl with 12.5 nM NSP3 Mac1 protein, 400 nM peptide ARTK(Bio)QTARK(Aoa-RADP)S (Cambridge Peptides), 1:20000 Anti-His6-Eu3+ cryptate (HTRF donor, PerkinElmer) and 1:125 Streptavidin-XL665 (HTRF acceptor, PerkinElmer) in assay buffer (25 mM 4-(2-hydroxyethyl)-1-piperazineethanesulfonic acid (HEPES) pH 7.0, 20 mM NaCl, 0.05% bovine serum albumin and 0.05% Tween-20). TARG1 and MacroD2 binding were measured at 100 nM and 12.5 nM, respectively. Assay reagents were dispensed manually into plates using a multichannel pipette while macrodomain protein and peptide were first dispensed and incubated for 30 min at room temperature. This was followed by addition of the HTRF reagents and incubation at room temperature for 1 h. Fluorescence was measured using a PHERAstar microplate reader (BMG) using the HTRF module with dual emission protocol (A = excitation of 320 nm, emission of 665 nm, and B = excitation of 320 nm, emission of 620 nm) or a Synergy H1 (Biotek) using the HTRF filter set (A =excitation 330/80 nm, emission of 620/10 nm, and B = excitation of 330/80 nm and emission of 665/8 nm). Raw data were processed to give an HTRF ratio (channel A/B × 10,000), which was used to generate IC_50_ curves. The IC_50_ values were determined by nonlinear regression using GraphPad Prism v.8 (GraphPad Software, CA, USA).

### Isothermal Titration Calorimetry and estimation of *K_i_* values

To determine *K_i_* values from the obtained HTRF IC_50_s, binding experiments were carried out on a VP-ITC microcalorimeter (MicroCal) to determine the dissociation constant, *K*_D_, of Mac1 for the ADPr-peptide used in the HTRF assay. The protein was dialysed overnight at 4°C in ITC buffer (25 mM HEPES pH 7.0 and 20 mM NaCl) using D-tube Dialysis Midi MWCO 3.5 kDa (Novagen) dialysis tubes before the experiment. Titration experiments were then performed at 22°C, a reference power of 12 μCal s^-1^ and a stirring speed of 307 rpm with an initial injection of 2 μl followed by 27 identical injections of 10 μl (duration of 4 s per injection and spacing of 240 s between injections). Data were analyzed using the MicroCal PEAQ-ITC analysis software (Malvern). *K_i_* values were calculated using the Cheng-Prusoff equation (19).

### Differential Scanning Fluorimetry

DSF and associated compound handling was performed as described (12), with 5 μM dye “Fluorescent Yellow” (Jacquard iDye Cat #JID1405) used in place of SYPRO Orange. Compounds were tested in triplicate, at seven concentrations in two-fold serial dilutions, at a top concentration of either 1000 or 100 μM. Data were analyzed using DSFworld (57) by fitting raw RFU values from 25 to 85°C to the second DSFworld model (single transition with initial decay). For each compound, the Spearman coefficient was calculated between compound concentration and ΔTma. A “DSF positive” compound was defined as any compound which met all three criteria: a positive mean thermal shift ≥0.5°C at any tested concentration, positive Spearman estimate, and Spearman p value ≤0.05. All data used to determine temperature shifts by DSF are included in **Dataset S7**.

### MDR1-MDCK II cell permeability

Permeability of compounds was assessed using canine MDR1 knockout, human MDR1 knockin MDCKII cells (MDR1-MDCKII) (Sigma-Aldrich, MTOX1303) in confluent monolayers expressing P-glycoprotein (P-gp) at Enamine biological services Bienta LTD (Kyiv, Ukraine). Cell suspension (400 μl) was added to each well of high throughput screening multiwell insert system plates. Test compounds were prepared as 20 mM DMSO stocks. The test compound (300 μl) was dissolved in transport buffer (9.5 g/l Hanks’ balanced salt solution and 0.35 g/l NaHCO_3_ with 0.81 mM MgSO_4_, 1.26 mM CaCl_2_, 25 mM HEPES, pH adjusted to 7.4) and added into filter wells whereas 1000 μl of transport buffer was added to transport analysis plate wells in order to determine apical (A) to basolateral (B) transport. Basolateral to apical transport was measured by adding 1000 μl of the test compound solution into the transport analysis plate wells whereas 300 μl of buffer was used to fill the filter plate wells. Final concentrations of test compounds were 10 μM. Plates were incubated for 90 min at 37°C under continuous shaking (100 rpm), 75 μl aliquots were taken from the donor and receiver compartments for LC-MS/MS analysis. Samples were mixed with acetonitrile followed by protein sedimentation by centrifugation at 1000 rpm for 10min. HPLC coupled with tandem mass spectroscopy was performed using the Shimadzu Prominence HPLC system coupled with the API 5000 (PE Sciex) spectrometer. Both the positive and negative ion modes of the TurboIonSpray ion source were used. The apparent permeability (Papp) was computed using the equation 1, where *V_A_* is the volume of transport buffer in acceptor wel, *Area* is the surface area of the insert, *Time* is the assay time, *[drug]_acc_* is the peak area of test compound in acceptor well, and *[drug]_initial,d_* is the initial amount of the test compound in a donor well.

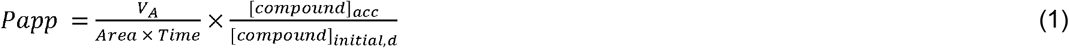

## Acknowledgments

The X-ray crystal structures reported in this work were determined using diffraction data collected at the ALS, the SSRL and the NSLS-II. The ALS, a U.S. DOE Office of Science User Facility under contract no. DE-AC02-05CH11231, is supported in part by the ALS-ENABLE program funded by the NIH, National Institute of General Medical Sciences, grant P30 GM124169-01. Use of the SSRL, SLAC National Accelerator Laboratory, is supported by the U.S. Department of Energy, Office of Science, Office of Basic Energy Sciences under contract no. DE-AC02-76SF00515. The SSRL Structural Molecular Biology Program is supported by the DOE Office of Biological and Environmental Research and by the NIH, National Institute of General Medical Sciences (P30GM133894). This research used beamline 17-ID-2 of the NSLS-II, the DOE Office of Science User Facility operated for the DOE Office of Science by Brookhaven National Laboratory under contract no. DE-SC0012704. The Center for BioMolecular Structure (CBMS) is primarily supported by the NIH, NIGMS through a Center Core P30 Grant (P30GM133893), and by the DOE Office of Biological and Environmental Research (KP1605010). Structural biology applications used for the analysis and modeling of crystallographic data were compiled and configured by SBGrid (58). We acknowledge the use of the Wynton high-performance compute cluster at UCSF.

This work was supported by the NIH and NIAID Antiviral Drug Discovery (AViDD) grant (U19AI171110) (to J.S.F., B.K.S., A.R., A.A.), by NSF Rapid 2031205, and a TMC Award from the UCSF Program for Breakthrough Biomedical Research, funded in part by the Sandler Foundation (to J.S.F. and J.E.G.), by NIH R35GM122481 and DARPA HR0011-19-2-0020 (to B.K.S.); GM141299 (to J.E.G.); and by GM133836 and GM071896 (to J.J.I.); by the Wellcome Trust (210634 and 223107), Oxford University Challenge Seed Fund (USCF 456), Biotechnology and Biological Sciences Research Council (BB/R007195/1), Ovarian Cancer Research Alliance (813369) and Cancer Research United Kingdom (C35050/A22284) (to I.A.) as well as NIHR Oxford Biomedical Research Centre (to J.C.T.).

## References

1. W. Yan, Y. Zheng, X. Zeng, B. He, W. Cheng, Structural biology of SARS-CoV-2: open the door for novel therapies. Signal Transduct Target Ther 7, 26 (2022).

2. A. K. L. Leung, D. E. Griffin, J. Bosch, A. R. Fehr, The Conserved Macrodomain Is a Potential Therapeutic Target for Coronaviruses and Alphaviruses. Pathogens 11, 94 (2022).

3. H. Yang, Z. Rao, Structural biology of SARS-CoV-2 and implications for therapeutic development. Nat. Rev. Microbiol. 19, 685–700 (2021).

4. W. Fu, et al., The search for inhibitors of macrodomains for targeting the readers and erasers of mono-ADP-ribosylation. Drug Discov. Today 26, 2547–2558 (2021).

5. J. G. M. Rack, et al., Viral macrodomains: a structural and evolutionary assessment of the pharmacological potential. Open Biol. 10, 200237 (2020).

6. L. C. Russo, et al., The SARS-CoV-2 Nsp3 macrodomain reverses PARP9/DTX3L-dependent ADP-ribosylation induced by interferon signaling. J. Biol. Chem. 297, 101041 (2021).

7. Y. M. O. Alhammad, et al., The SARS-CoV-2 Conserved Macrodomain Is a Mono-ADP-Ribosylhydrolase. J. Virol. 95(2021).

8. A. R. Fehr, et al., The Conserved Coronavirus Macrodomain Promotes Virulence and Suppresses the Innate Immune Response during Severe Acute Respiratory Syndrome Coronavirus Infection. MBio 7(2016).

9. L. M. Sherrill, et al., Design, synthesis and evaluation of inhibitors of the SARS-CoV-2 nsp3 macrodomain. Bioorg. Med. Chem. 67, 116788 (2022).

10. A. Roy, et al., Discovery of compounds that inhibit SARS-CoV-2 Mac1-ADP-ribose binding by high-throughput screening. Antiviral Res. 203, 105344 (2022).

11. S. T. Sowa, et al., A molecular toolbox for ADP-ribosyl binding proteins. Cell Rep Methods 1, 100121 (2021).

12. M. Schuller, et al., Fragment binding to the Nsp3 macrodomain of SARS-CoV-2 identified through crystallographic screening and computational docking. Sci Adv 7(2021).

13. G. J. Correy, et al., The mechanisms of catalysis and ligand binding for the SARS-CoV-2 NSP3 macrodomain from neutron and x-ray diffraction at room temperature. Sci Adv 8, eabo5083 (2022).

14. M. Ferla, Scaffold hopping between bound compounds by stitching them together like a reanimated corpse. Fragmenstein.

15. O. O. Grygorenko, et al., Erratum: Generating Multibillion Chemical Space of Readily Accessible Screening Compounds. iScience 23, 101873 (2020).

16. S. B. Shuker, P. J. Hajduk, R. P. Meadows, S. W. Fesik, Discovering high-affinity ligands for proteins: SAR by NMR. Science 274, 1531–1534 (1996).

17. J. J. Irwin, et al., ZINC20—A Free Ultralarge-Scale Chemical Database for Ligand Discovery. J. Chem. Inf. Model. 60, 6065–6073 (2020).

18. N. M. Pearce, et al., A multi-crystal method for extracting obscured crystallographic states from conventionally uninterpretable electron density. Nat. Commun. 8, 15123 (2017).

19. C. Yung-Chi, W. H. Prusoff, Relationship between the inhibition constant (KI) and the concentration of inhibitor which causes 50 per cent inhibition (I50) of an enzymatic reaction. Biochem. Pharmacol. 22, 3099–3108 (1973).

20. X. Yang, et al., Molecular basis for the MacroD1-mediated hydrolysis of ADP-ribosylation. DNA Repair 94, 102899 (2020).

21. J. Lyu, et al., Ultra-large library docking for discovering new chemotypes. Nature 566, 224–229 (2019).

22. K. Michalska, et al., Crystal structures of SARS-CoV-2 ADP-ribose phosphatase: from the apo form to ligand complexes. IUCrJ 7, 814–824 (2020).

23. T. Sterling, J. J. Irwin, ZINC 15 – Ligand Discovery for Everyone. Journal of Chemical Information and Modeling 55, 2324–2337 (2015).

24. B. J. Bender, et al., A practical guide to large-scale docking. Nat. Protoc. 16, 4799–4832 (2021).

25. W. J. Allen, R. C. Rizzo, Implementation of the Hungarian algorithm to account for ligand symmetry and similarity in structure-based design. J. Chem. Inf. Model. 54, 518–529 (2014).

26. Q. Zhao, et al., Enhanced Sampling Approach to the Induced-Fit Docking Problem in Protein-Ligand Binding: The Case of Mono-ADP-Ribosylation Hydrolase Inhibitors. J. Chem. Theory Comput. 17, 7899–7911 (2021).

27. M. Jaskolski, A. Wlodawer, Z. Dauter, W. Minor, B. Rupp, Group depositions to the Protein Data Bank need adequate presentation and different archiving protocol. Protein Sci. 31,784–786 (2022).

28. W. Jahnke, et al., Fragment-to-Lead Medicinal Chemistry Publications in 2019. J. Med. Chem. 63, 15494–15507 (2020).

29. I. J. P. de Esch, D. A. Erlanson, W. Jahnke, C. N. Johnson, L. Walsh, Fragment-to-lead medicinal chemistry publications in 2020. J. Med. Chem. 65, 84–99 (2022).

30. C. R. Søndergaard, A. E. Garrett, T. Carstensen, G. Pollastri, J. E. Nielsen, Structural artifacts in protein-ligand X-ray structures: implications for the development of docking scoring functions. J. Med. Chem. 52, 5673–5684 (2009).

31. D. Chen, et al., Identification of macrodomain proteins as novel O-acetyl-ADP-ribose deacetylases. J. Biol. Chem. 286, 13261–13271 (2011).

32. M. Schuller, et al., Discovery of a Selective Allosteric Inhibitor Targeting Macrodomain 2 of Polyadenosine-Diphosphate-Ribose Polymerase 14. ACS Chem. Biol. 12, 2866–2874 (2017).

33. L. Wang, et al., Accurate and reliable prediction of relative ligand binding potency in prospective drug discovery by way of a modern free-energy calculation protocol and force field. J. Am. Chem. Soc. 137, 2695–2703 (2015).

34. V. Gapsys, et al., Accurate absolute free energies for ligand–protein binding based on nonequilibrium approaches. Communications Chemistry 4, 1–13 (2021).

35. R. G. Coleman, M. Carchia, T. Sterling, J. J. Irwin, B. K. Shoichet, Ligand Pose and Orientational Sampling in Molecular Docking. PLoS ONE 8, e75992 (2013).

36. K. Gallagher, K. Sharp, Electrostatic Contributions to Heat Capacity Changes of DNA-Ligand Binding. Biophysical Journal 75, 769–776 (1998).

37. S. J. Weiner, et al., A new force field for molecular mechanical simulation of nucleic acids and proteins. J. Am. Chem. Soc. 106, 765–784 (1984).

38. M. M. Mysinger, B. K. Shoichet, Rapid context-dependent ligand desolvation in molecular docking. J. Chem. Inf. Model. 50, 1561–1573 (2010).

39. G. Madhavi Sastry, M. Adzhigirey, T. Day, R. Annabhimoju, W. Sherman, Protein and ligand preparation: parameters, protocols, and influence on virtual screening enrichments. J. Comput. Aided Mol. Des. 27, 221–234 (2013).

40. M. H. M. Olsson, C. R. Søndergaard, M. Rostkowski, J. H. Jensen, PROPKA3: Consistent Treatment of Internal and Surface Residues in Empirical pKa Predictions. J. Chem. Theory Comput. 7, 525–537 (2011).

41. R. M. Stein, et al., Property-Unmatched Decoys in Docking Benchmarks. J. Chem. Inf. Model. 61, 699–714 (2021).

42. S. Gu, M. S. Smith, Y. Yang, J. J. Irwin, B. K. Shoichet, Ligand Strain Energy in Large Library Docking. J. Chem. Inf. Model. 61, 4331–4341 (2021).

43. A. V. Fassio, et al., Prioritizing virtual screening with interpretable interaction fingerprints. bioRxiv, 2022.05.25.493419 (2022).

44. D. Alvarez-Garcia, X. Barril, Molecular simulations with solvent competition quantify water displaceability and provide accurate interaction maps of protein binding sites. J. Med. Chem. 57, 8530–8539 (2014).

45. J. Wang, R. M. Wolf, J. W. Caldwell, P. A. Kollman, D. A. Case, Development and testing of a general amber force field. J. Comput. Chem. 25, 1157–1174 (2004).

46. P. M. Collins, et al., Gentle, fast and effective crystal soaking by acoustic dispensing. Acta Crystallogr D Struct Biol 73, 246–255 (2017).

47. W. Kabsch, XDS. Acta Crystallogr. D Biol. Crystallogr. 66, 125–132 (2010).

48. P. A. Karplus, K. Diederichs, Linking crystallographic model and data quality. Science 336, 1030–1033 (2012).

49. P. R. Evans, G. N. Murshudov, How good are my data and what is the resolution? Acta Crystallogr. D Biol. Crystallogr. 69, 1204–1214 (2013).

50. R. Keegan, M. Wojdyr, G. Winter, A. Ashton, DIMPLE: a difference map pipeline for the rapid screening of crystals on the beamline. Acta Crystallogr. A Found. Adv. 71, s18–s18 (2015).

51. M. D. Winn, et al., Overview of the CCP4 suite and current developments. Acta Crystallogr. D Biol. Crystallogr. 67, 235–242 (2011).

52. P. Emsley, B. Lohkamp, W. G. Scott, K. Cowtan, Features and development of Coot. Acta Crystallogr. D Biol. Crystallogr. 66, 486–501 (2010).

53. N. W. Moriarty, R. W. Grosse-Kunstleve, P. D. Adams, electronic Ligand Builder and Optimization Workbench (eLBOW): a tool for ligand coordinate and restraint generation. Acta Crystallogr. D Biol. Crystallogr. 65, 1074–1080 (2009).

54. F. Long, et al., AceDRG: a stereochemical description generator for ligands. Acta Crystallogr D Struct Biol 73, 112–122 (2017).

55. L. L. C. Schrödinger, LigPrep.

56. P. V. Afonine, et al., Towards automated crystallographic structure refinement with phenix.refine. Acta Crystallogr. D Biol. Crystallogr. 68, 352–367 (2012).

57. T. Wu, et al., Three Essential Resources to Improve Differential Scanning Fluorimetry (DSF) Experiments. bioRxiv, 2020.03.22.002543 (2020).

58. A. Morin, et al., Collaboration gets the most out of software. Elife 2, e01456 (2013).

