## Supplementary Information for "Iterative computational design and crystallographic screening identifies potent inhibitors targeting the Nsp3 Macrodomain of SARS-CoV-2"

**This PDF file includes:**

Figures S1 to S9

Legends for Datasets S1 to S6

**Other supporting materials for this manuscript include the following:**

Datasets S1 to S6

**Supporting Information Text**

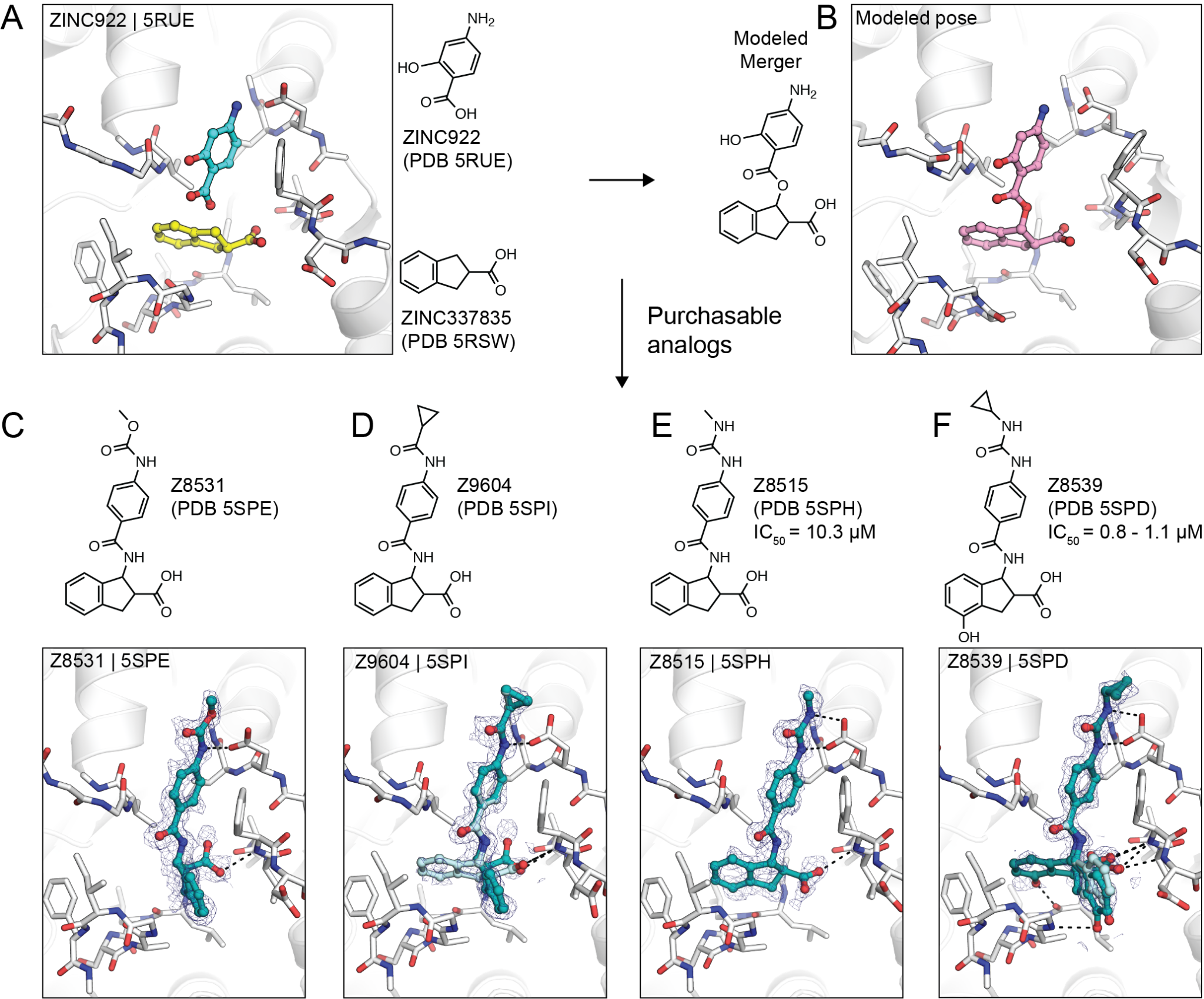

**Fig. S1.** **Overview of fragment linking approach that led to Z8539.** **A)** Alignment of X-ray crystal structures of the fragments ZINC922 and ZINC337835. **B)** Linked fragments designed by *Fragmenstein* modeled into the Mac1 active site. **C-F)** 2D and X-ray crystal structures of purchased and tested analogs of the *Fragmenstein*-derived merger. PanDDA event maps are shown for ligands (blue mesh contoured at 2 σ).

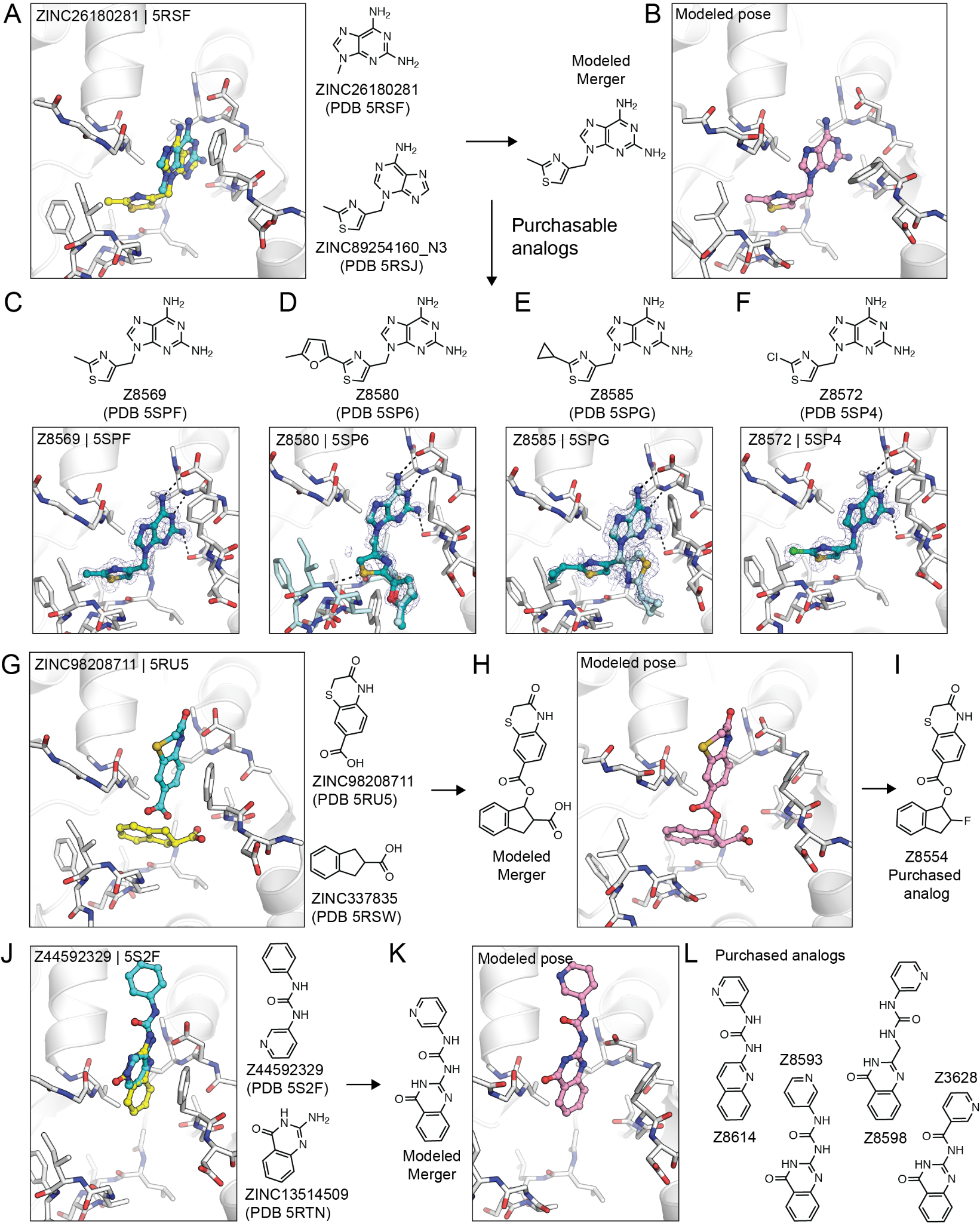

**Fig. S2. Fragment linking and merging with *Fragmenstein*.** **A)** Overlay of the ZINC26180281 and ZINC89254160_N3 fragments identified in the initial screen (1). **B)** Modeled ZINC26180281-ZINC89254160_N3 merger. **C-F)** 2D and X-ray crystal structures of purchased and tested analogs of the ZINC26180281-ZINC89254160_N3 merger. PanDDA event maps are shown around the ligands (blue mesh contoured at 2 σ). **G)** Overlay of the ZINC98208711 and ZINC337835 fragments. **H)** Modeled ZINC98208711-ZINC337835 merger. **I)** Purchased and tested analog of the ZINC98208711-ZINC337835 merger. **J)** Overlay of the Z44592329 and ZINC13514509 fragments. K) Modeled Z44592329-ZINC13514509 merger. L) Purchased and tested analogs of the Z44592329-ZINC13514509 merger.

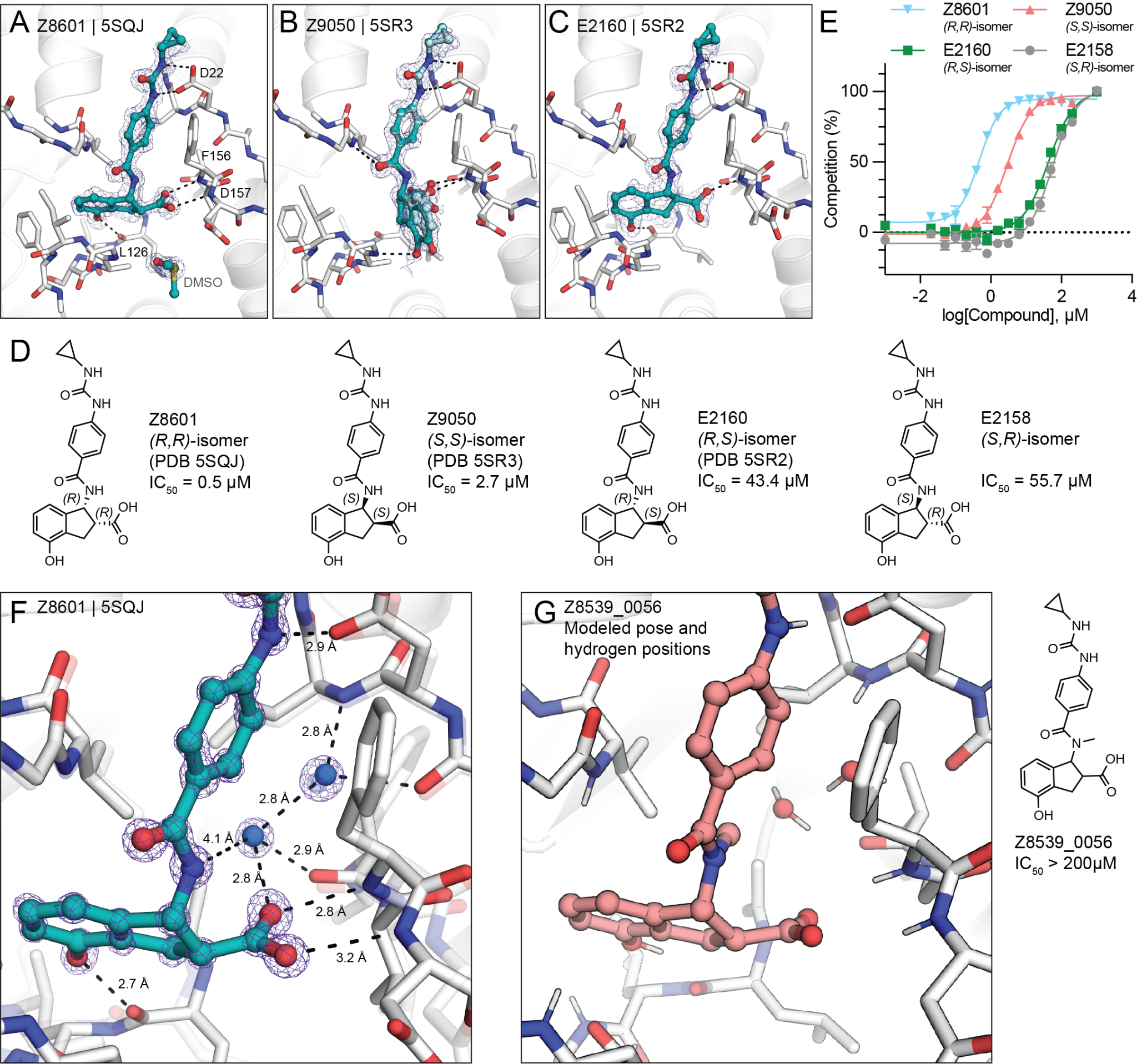

**Figure S3. Chiral separation and testing of the Z8539 stereoisomers. A-C)** Crystal structures of three of the four possible Z8539 stereoisomers. PanDDA event maps are shown around the ligands (blue mesh contoured at 2 σ). **D)** 2D structures and HTRF-derived IC_50_ values of Z8539 stereoisomers. **E)** HTRF-based binding curves of Z8539 stereoisomers. Water-mediated interaction between Mac1 and Z8601. **F)** X-ray crystal structure of Mac1 bound to the (*R*,*R*) isomer of Z8539. The 2mF_O_-DF_C_ electron density is shown around the ligand and nearby solvent after refinement (purple mesh contoured at 3 σ). The ligand-bound state (teal/white sticks) was refined at 78% occupancy and the apo state (transparent sticks and spheres) was refined at 22% occupancy. **G)** Same view as **(F)**, but showing the modeled binding pose of the inactive, methylated analog Z8539_0056 with nearby solvent.

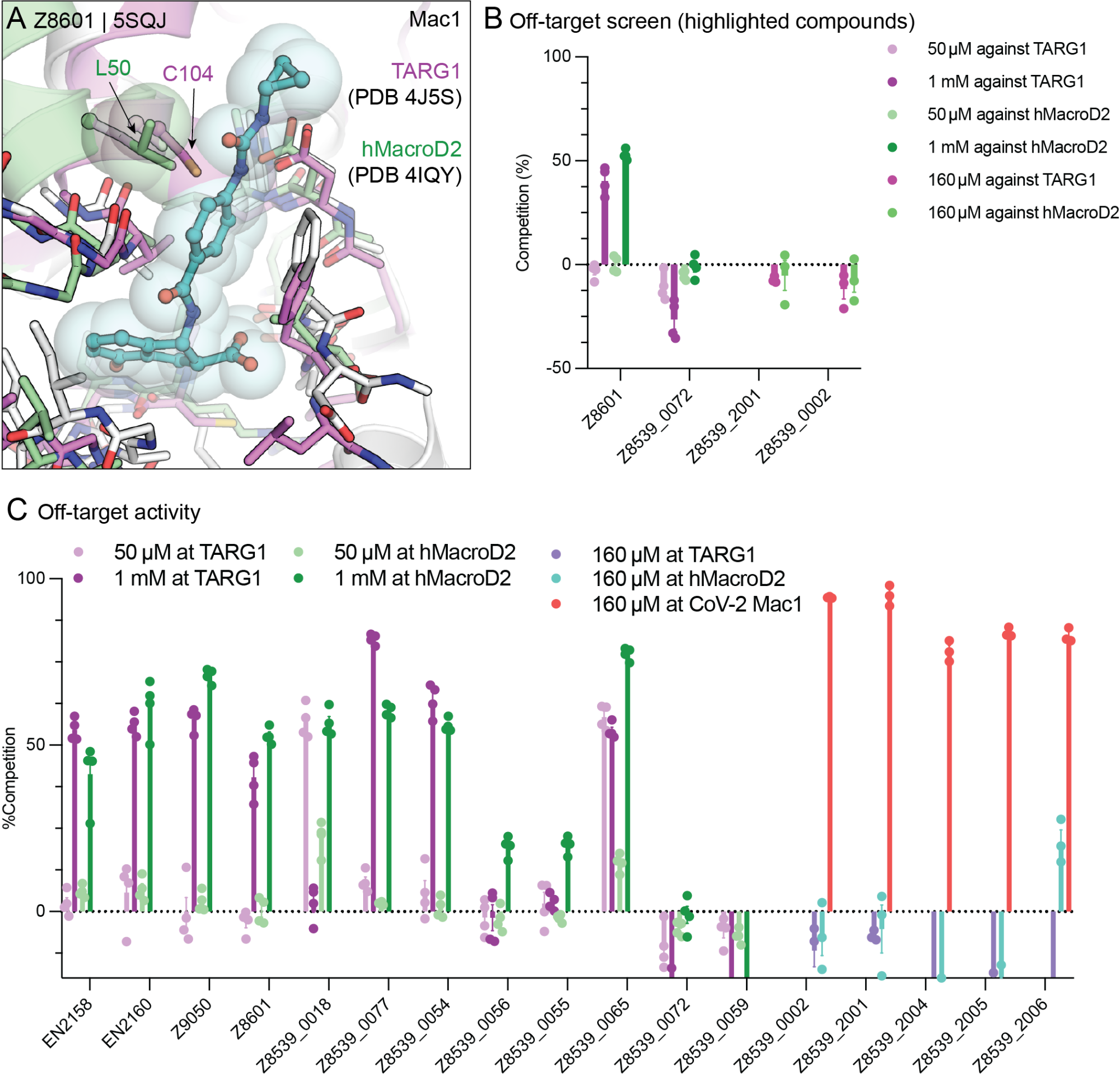

**Figure S4.** **Assessment of ligand selectivity against the human macrodomains MacroD2 and TARG1. A)** Crystal structures of Mac1, TARG1 (2) and MacroD2 (3) were aligned by superimposing the co-crystallized ADPr (7KQP was used for Mac1, not shown). **B,C)** Off-target activities of Z8539 analogs at human macrodomains TARG1 and MacroD2.

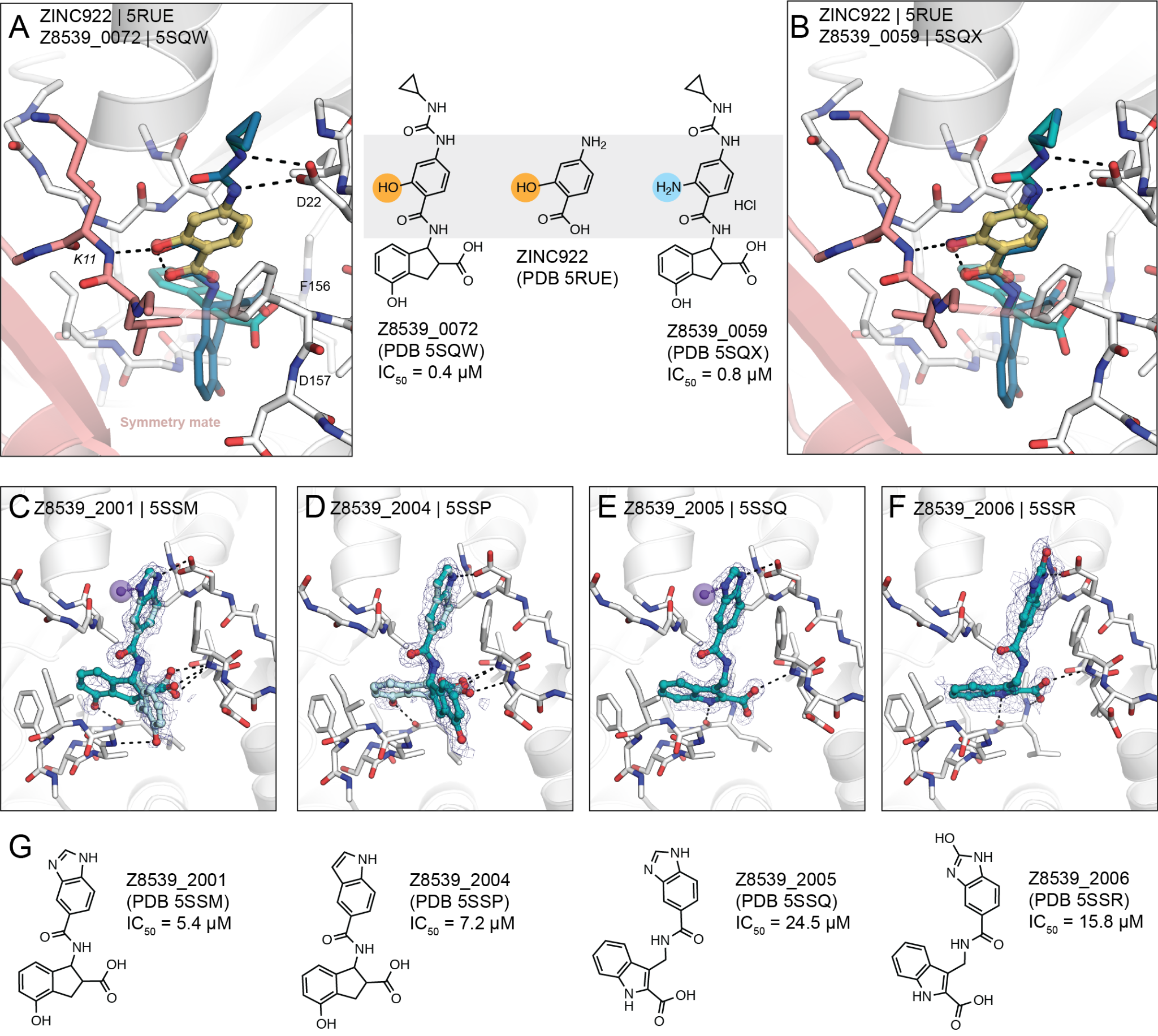

**Figure S5. Crystal packing interactions of Z8539 derivatives. A-B)** Interactions of Z8539_0072 or Z8539_0059 within the crystal lattice, respectively. As observed for the initial fragment hit ZINC922, the added hydroxyl (or amine) group forms a hydrogen to the Lys11 of the symmetry mate. **C-F)** Crystal structure of Z8539_2001, Z8539_2004, Z8539_2005 and Z8539_2006, respectively. PanDDA event maps are shown for the ligands (contoured at 2 σ). **G)** Corresponding 2D structures and HTRF-derived IC_50_ values.

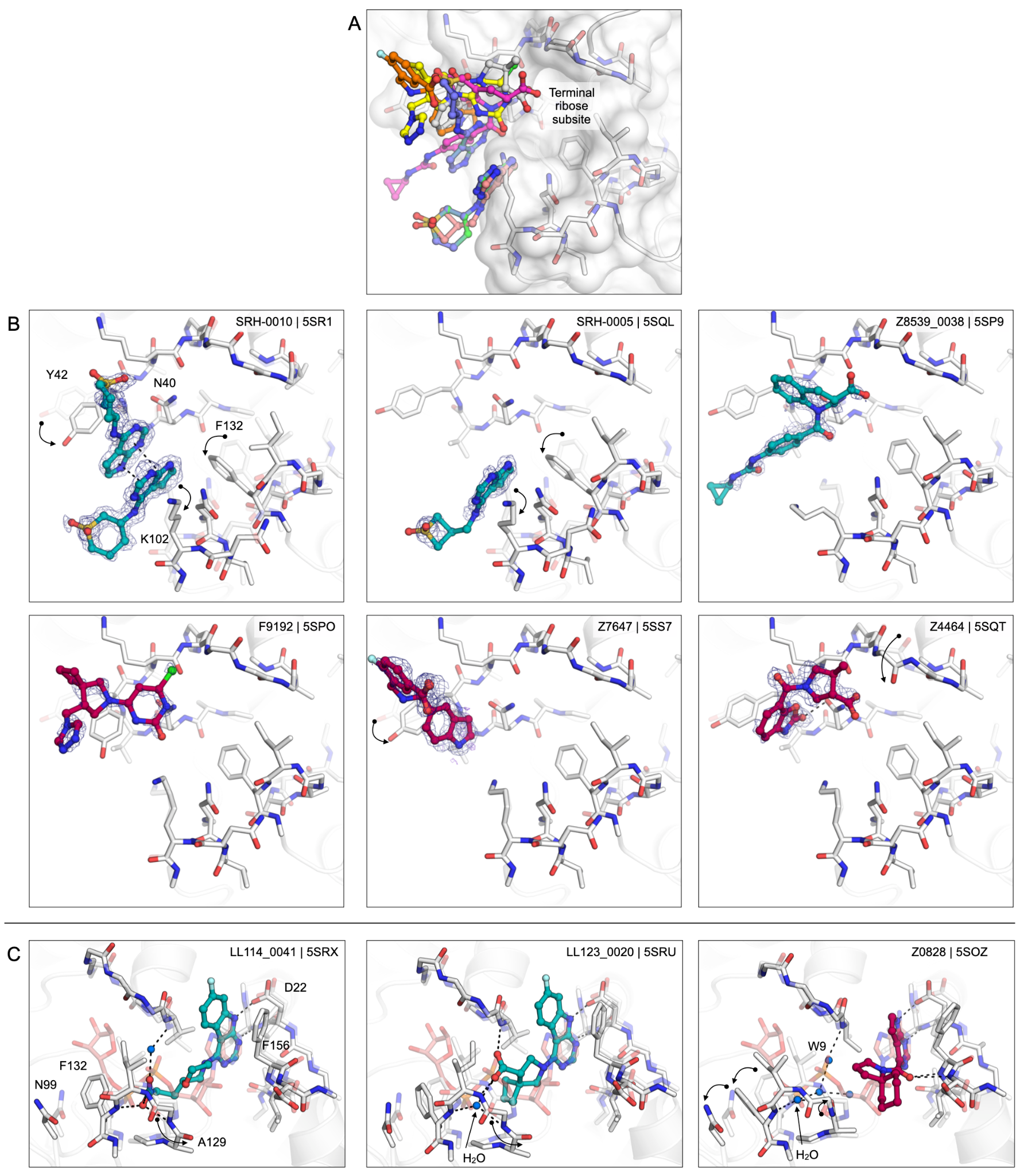

**Figure S6. Ligands binding in the catalytic site of Mac1. A)** Overlay of the six ligands identified binding in the catalytic site of Mac1. **B)** Structures of the six ligands with conformational changes relative to the apo structure (transparent white sticks) annotated with black arrows. PanDDA event maps are shown around the ligands (blue mesh contoured at 2 σ). **C)** LL114_0041 stabilizes an alternative, flipped conformation of Ala129 relative to the apo state (transparent white sticks). The flipped conformation matches the ADPr-bound structure (ADPr is shown with transparent red sticks). The flipped conformation is also stabilized in the structure with LL123_0020 bound, but a water molecule occupies the key phosphate binding site. A similar rearrangement of the water network is seen for Z0828.

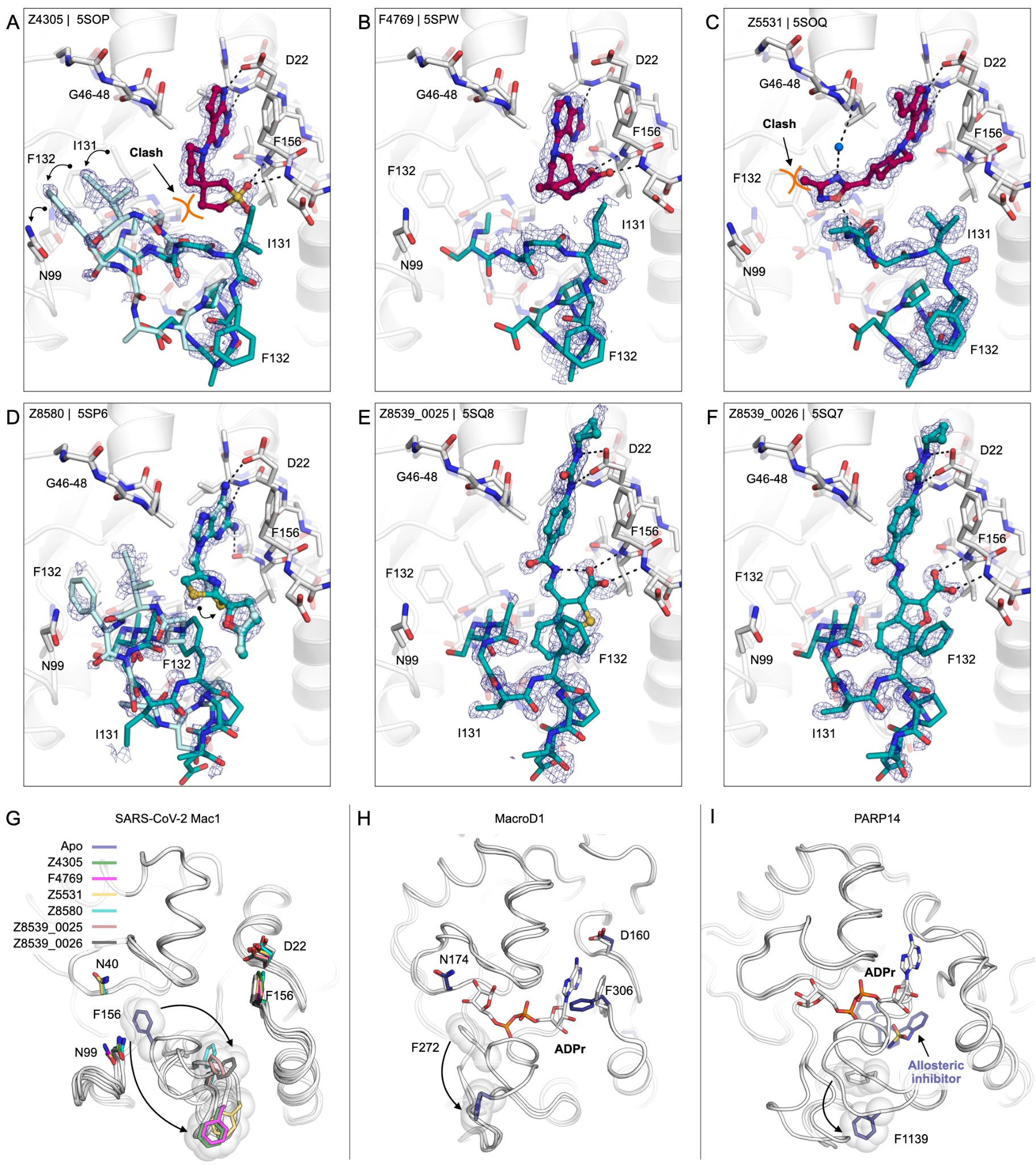

**Figure S7. Everted structures of Mac1. A**-**F)** X-ray crystal structures of the six ligands that stabilize the everted state of Mac1. PanDDA event maps are shown around the ligands and residues 129-135 (blue mesh contoured at 2 σ). Two conformations of the 129-135 loop were modeled in the structures of Z4305 and Z8580 (shown with teal/cyan sticks). **G)** Overlay of the six everted state structures with the apo state. Sticks are colored according to the ligand bound, and the ligands are omitted for clarity. **H)** Structure of MacroD1 showing Phe272 in an everted state (PDB 2X47 (4), blue sticks). The everted structure is overlaid with an ADPr-bound structure (PDB 6LH4 (5), white sticks). **I)** Structure of PARP14 macrodomain 2 bound to an allosteric inhibitor (PDB 5O2D (6), blue sticks). The structure is overlaid with an ADPr-bound structure (PDB 3Q71 (7), white sticks).

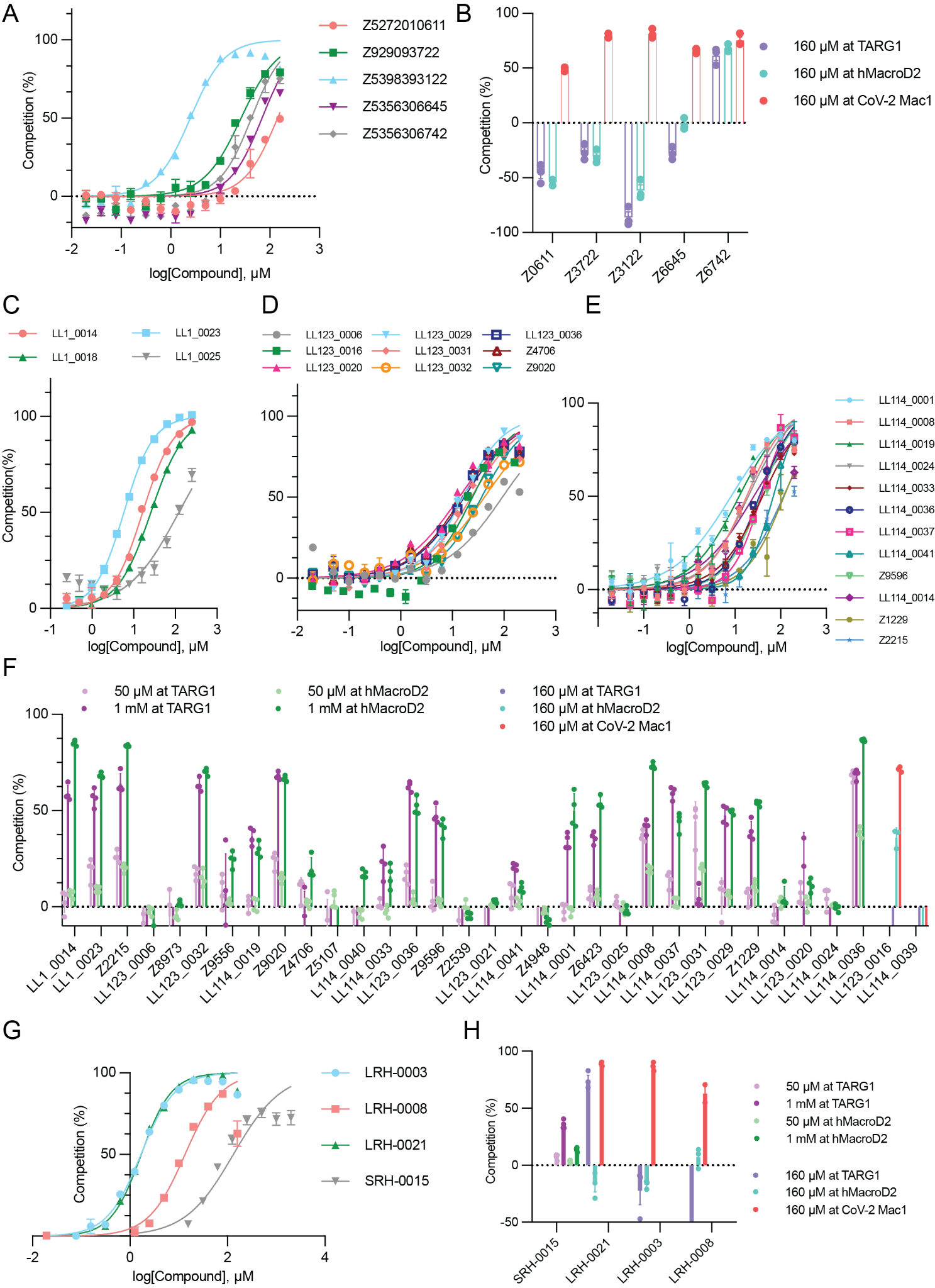

**Figure S8. Assessing activity and selectivity of Mac1 ligands. A)** HTRF-based peptide displacement assay of hits found against the everted structure of Mac1. Data are presented as the mean ± SEM of three technical repeats. **B)** Off-target activities of open-state hits. **C-E)** HTRF-based binding curves of compounds from the LL1 series. **F)** Off-target activities of LL1 compounds. **G)** HTRF-based binding curves of neutral probe set compounds. **H)** Off-target activities of neutral probe set hits.

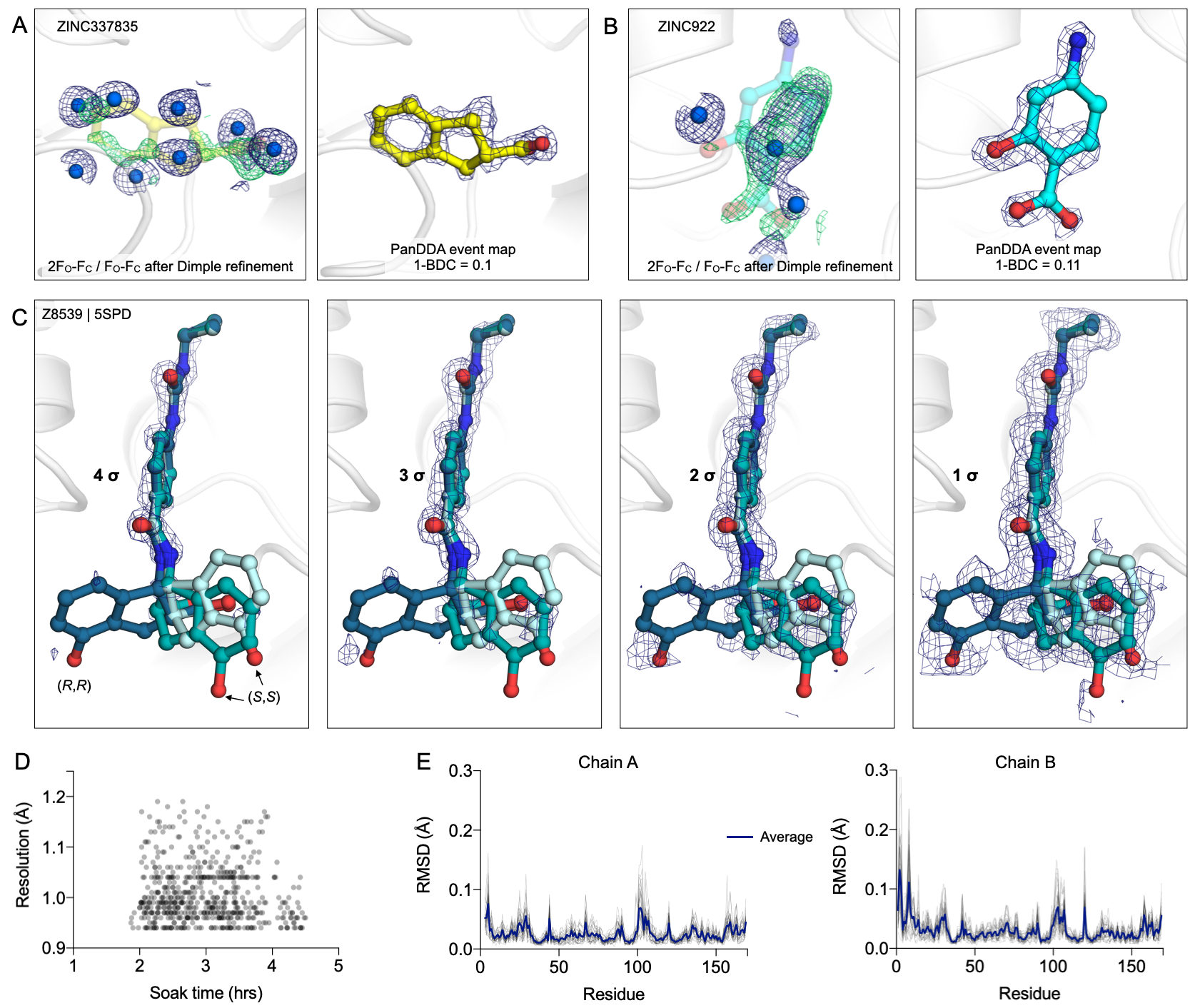

**Figure S9. PanDDA reveals clear density for low occupancy states. A)** Electron density maps after Dimple refinement (left) and PanDDA (right) from a crystal soaked with ZINC337835 (8). There is weak positive difference density consistent with the fragment in the F_O_-F_C_ map (green mesh contoured at 2 σ around the fragment), however this is largely obscured by modeled solvent in the 2F_O_-F_C_ map (blue mesh contoured at 1 σ around the fragment). In contrast, the fragment is unambiguously present in the PanDDA event map (blue mesh contoured at 2 σ around the fragment). **B)** Same as **(A)** but showing Dimple and PanDDA maps for ZINC922. **C)** PanDDA event maps helped to resolve conformational/chemical heterogeneity for the stereoisomeric mixture of Z8539 soaked into Mac1 crystals. Each panel shows the three modeled conformations of Z8539 [(*R*,*R*) and two conformations of (*S*,*S*)] with the PanDDA event density contoured from 4-1 σ. The 1-BDC value for the ligand binding event was 0.2, which is consistent with an occupancy of 40% (9). Given the relative intensity of the PanDDA map, this suggests that the (*R*,*R*) isomer is likely present at <10% occupancy. **D)** Plot showing resolution of all 591 datasets collected as a function of soak time. **E)** Plot showing per-residue heavy-atom RMSD of each of the 34 models obtained from crystals soaked in DMSO after alignment to the refinement starting model.

**Dataset S1 (separate file).** Spreadsheet showing all compounds tested in this work.

**Dataset S2 (separate file).** Spreadsheet documenting the X-ray crystallography results reported in this work.

**Dataset S3 (separate file).** Images showing crystal structures of Mac1 in complex with compounds from the Z8539 series.

**Dataset S4 (separate file).** Images showing crystal structures of Mac1 in complex with docking hits.

**Dataset S5 (separate file).** Images showing crystal structures of Mac1 in complex with analogs of docking hits (LL1 series).

**Dataset S6 (separate file).** Images showing design of neutral probe set.

**SI References**

1.	M. Schuller, *et al.*, Fragment Binding to the Nsp3 Macrodomain of SARS-CoV-2 Identified Through Crystallographic Screening and Computational Docking. *bioRxiv* (2020) https:/doi.org/[10.1101/2020.11.24.393405](http://dx.doi.org/10.1101/2020.11.24.393405).

2.	R. Sharifi, *et al.*, Deficiency of terminal ADP-ribose protein glycohydrolase TARG1/C6orf130 in neurodegenerative disease. *EMBO J.* **32**, 1225–1237 (2013).

3.	G. Jankevicius, *et al.*, A family of macrodomain proteins reverses cellular mono-ADP-ribosylation. *Nat. Struct. Mol. Biol.* **20**, 508–514 (2013).

4.	D. Chen, *et al.*, Identification of macrodomain proteins as novel O-acetyl-ADP-ribose deacetylases. *J. Biol. Chem.* **286**, 13261–13271 (2011).

5.	X. Yang, *et al.*, Molecular basis for the MacroD1-mediated hydrolysis of ADP-ribosylation. *DNA Repair* **94**, 102899 (2020).

6.	M. Schuller, *et al.*, Discovery of a Selective Allosteric Inhibitor Targeting Macrodomain 2 of Polyadenosine-Diphosphate-Ribose Polymerase 14. *ACS Chem. Biol.* **12**, 2866–2874 (2017).

7.	A. H. Forst, *et al.*, Recognition of mono-ADP-ribosylated ARTD10 substrates by ARTD8 macrodomains. *Structure* **21**, 462–475 (2013).

8.	M. Schuller, *et al.*, Fragment binding to the Nsp3 macrodomain of SARS-CoV-2 identified through crystallographic screening and computational docking. *Sci Adv* **7** (2021).

9.	N. M. Pearce, *et al.*, A multi-crystal method for extracting obscured crystallographic states from conventionally uninterpretable electron density. *Nat. Commun.* **8**, 15123 (2017).
