## Supplementary material for "Iterative computational design and crystallographic screening identifies potent inhibitors targeting the Nsp3 Macrodomain of SARS-CoV-2": Dataset S3

**Dataset S3:** Crystal structures of Mac1 in complex with compounds from the Z8539 series. PanDDA event maps are shown for ligands (contoured at 2 σ). Protein-ligand hydrogen bonds are shown with dashed black lines. Hydrogen bonds between ligands and the Lys11 backbone nitrogen of a symmetry mate are highlighted with purple spheres/dashes.

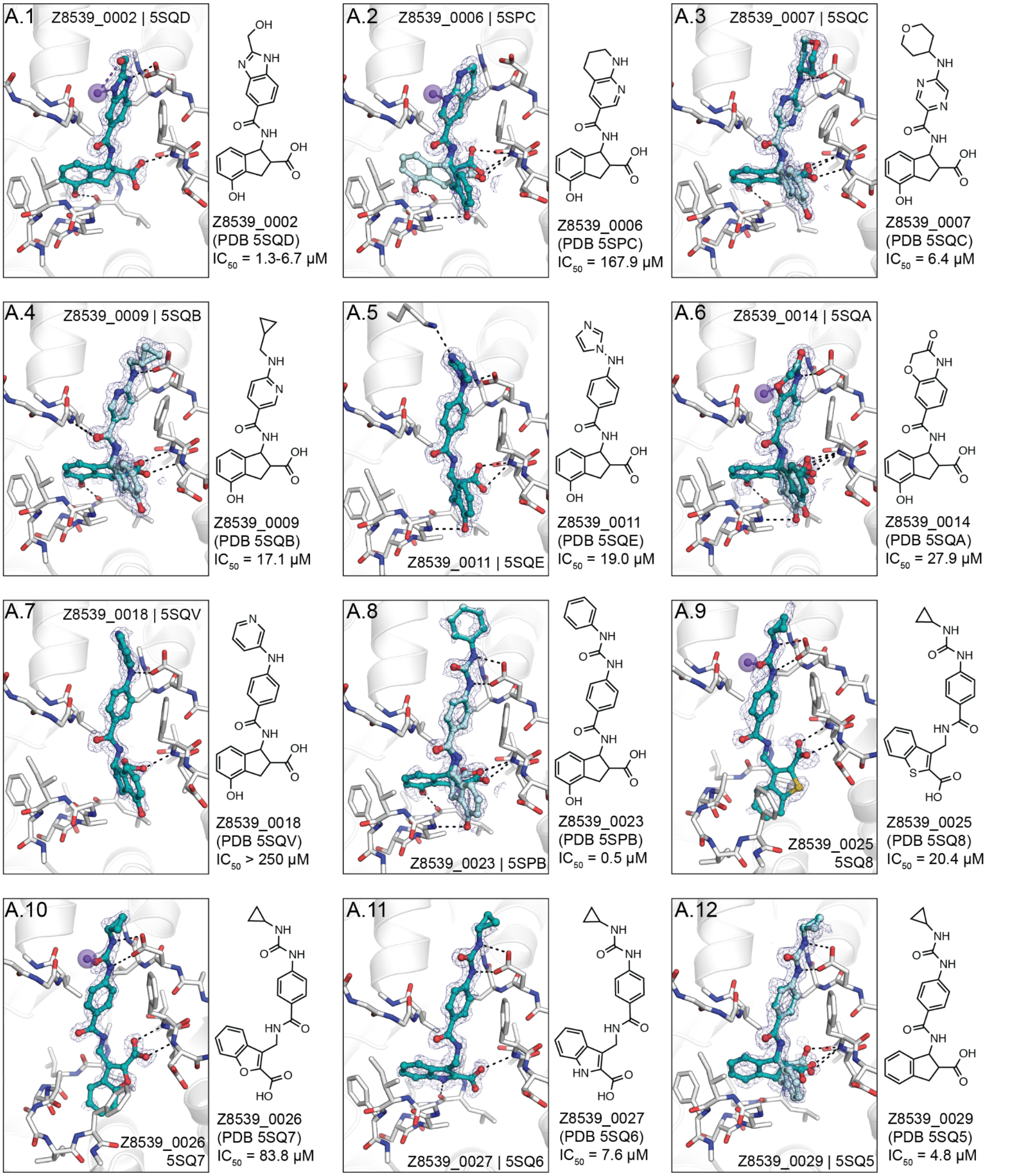

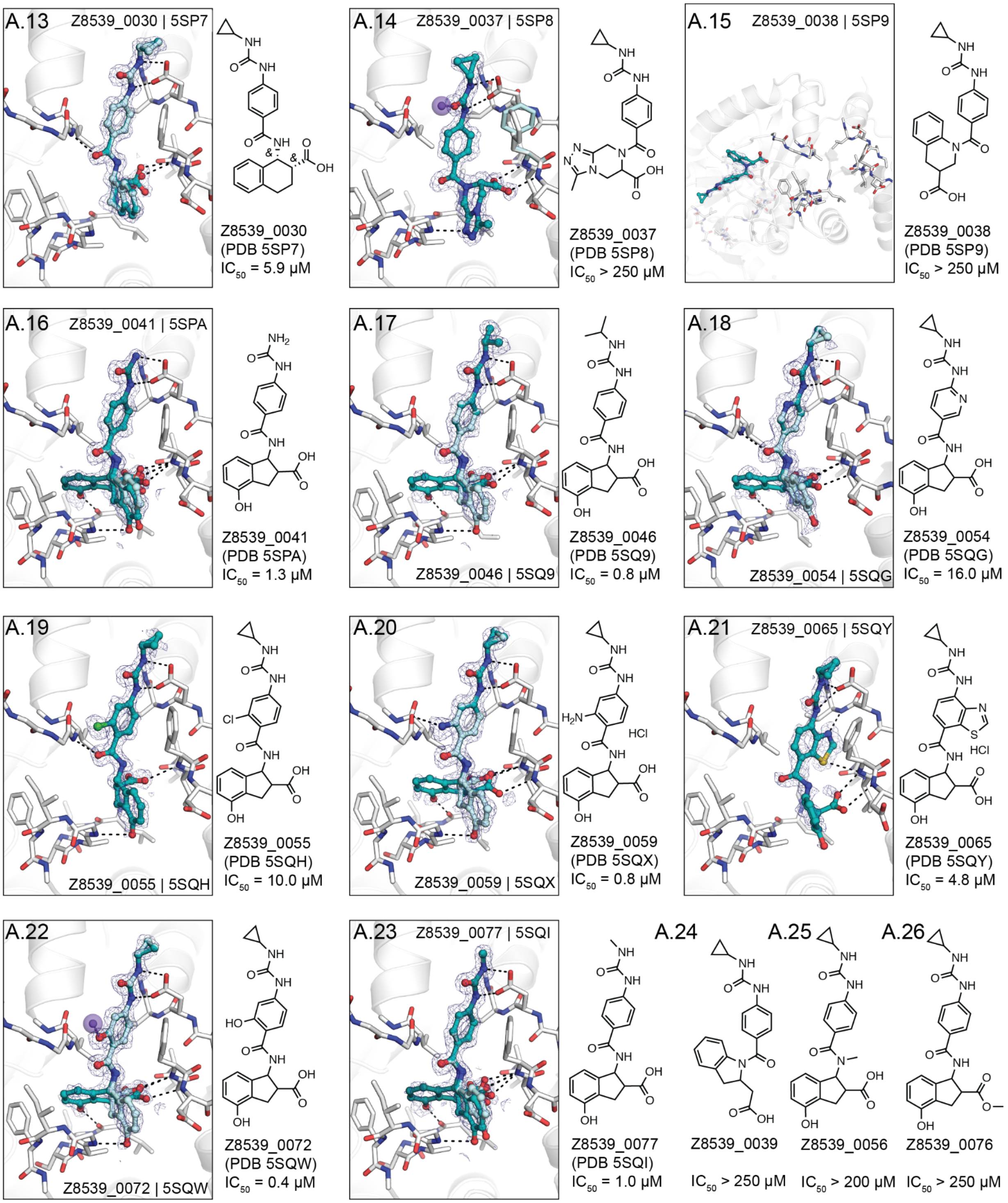
A.13
Z8539_0030 | 5SP7
Z8539_0030
(PDB 5SP7)
IC50 = 5.9 µM
A.14
Z8539_0037 | 5SP8
Z8539_0037
(PDB 5SP8)
IC50 > 250 µM
A.15
Z8539_0038 | 5SP9
Z8539_0038
(PDB 5SP9)
IC50 > 250 µM
A.16
Z8539_0041 | 5SPA
Z8539_0041
(PDB 5SPA)
IC50 = 1.3 µM
A.17
Z8539_0046 | 5SQ9
Z8539_0046
(PDB 5SQ9)
IC50 = 0.8 µM
A.18
Z8539_0054 | 5SQG
Z8539_0054
(PDB 5SQG)
IC50 = 16.0 µM
A.19
Z8539_0055 | 5SQH
Z8539_0055
(PDB 5SQH)
IC50 = 10.0 µM
A.20
Z8539_0059 | 5SQX
HCl
Z8539_0059
(PDB 5SQX)
IC50 = 0.8 µM
A.21
Z8539_0065 | 5SQY
HCl
Z8539_0065
(PDB 5SQY)
IC50 = 4.8 µM
A.22
Z8539_0072 | 5SQW
Z8539_0072
(PDB 5SQW)
IC50 = 0.4 µM
A.23
Z8539_0077 | 5SQI
Z8539_0077
(PDB 5SQI)
IC50 = 1.0 µM
A.24
Z8539_0039
IC50 > 250 µM
A.25
Z8539_0056
IC50 > 200 µM
A.26
Z8539_0076
IC50 > 250 µM
