## Supplementary material for "Iterative computational design and crystallographic screening identifies potent inhibitors targeting the Nsp3 Macrodomain of SARS-CoV-2": Dataset S4

**Dataset S4:** Crystal structures of Mac1 in complex with docking hits. PanDDA event maps are shown for ligands (contoured at 2 σ). Protein-ligand hydrogen bonds are shown with dashed black lines. Hydrogen bonds between ligands and the Lys11 backbone nitrogen of a symmetry mate are highlighted with purple spheres/dashes.

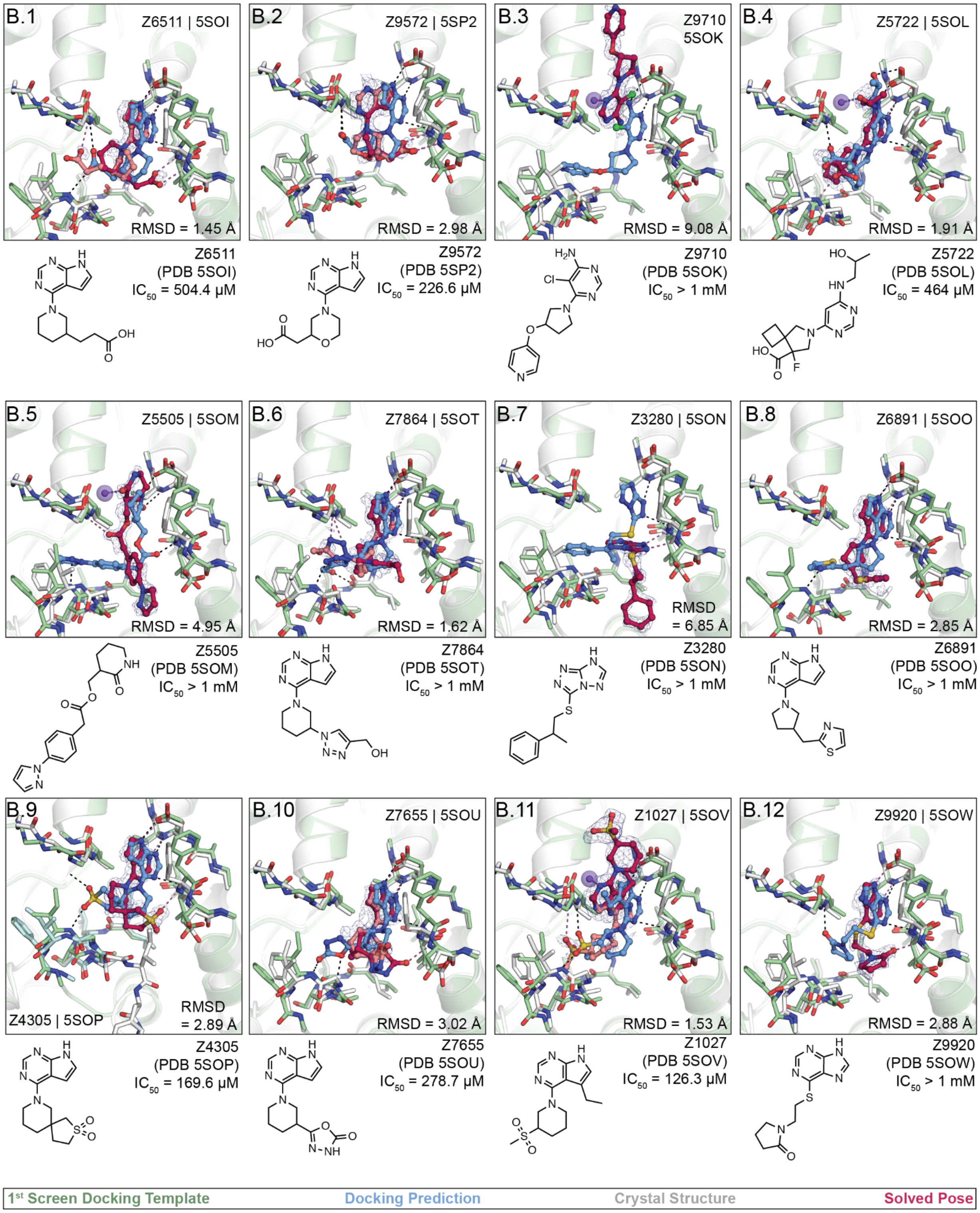

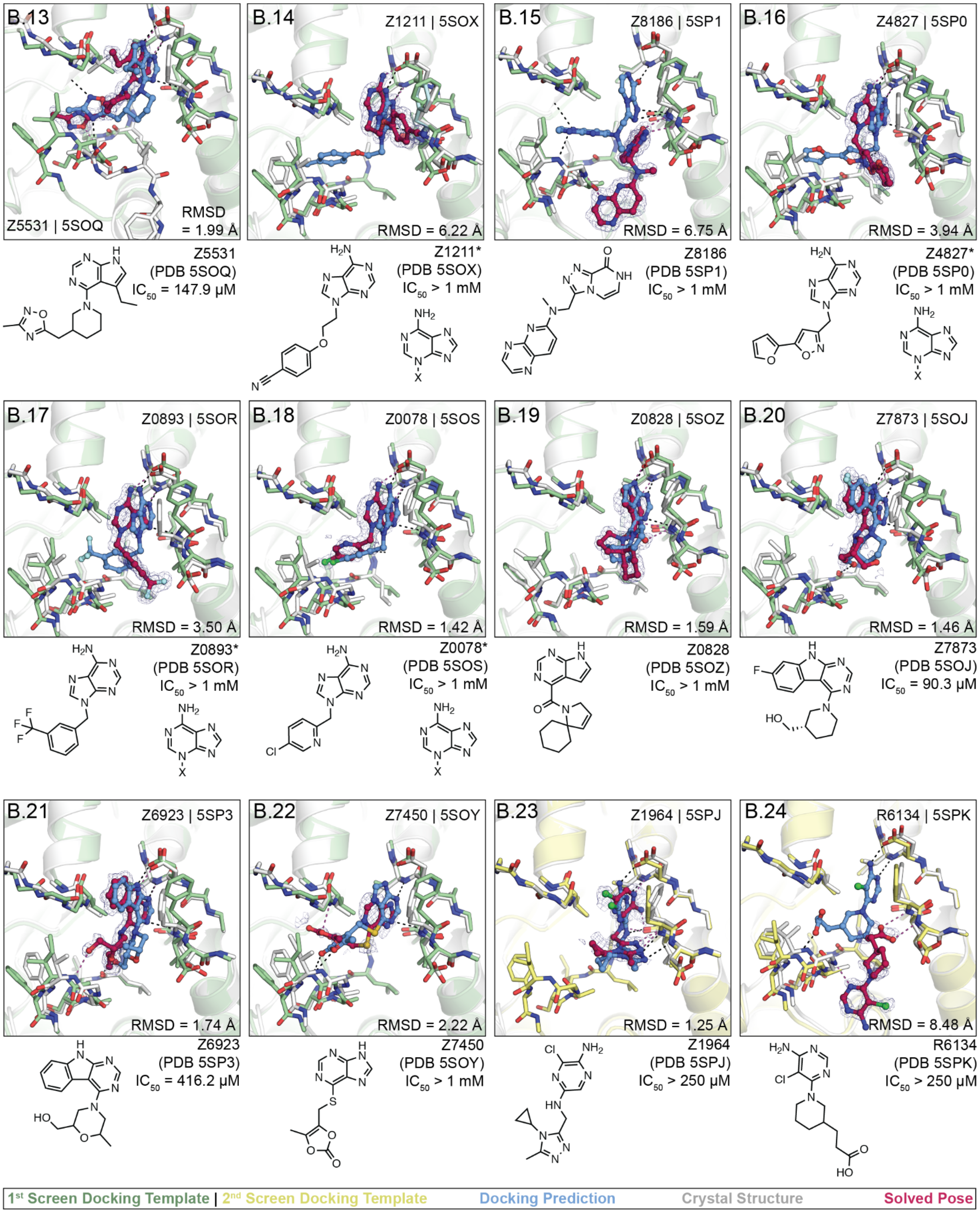

B.13
Z5531 | 5SOQ
RMSD = 1.99 Å
Z5531 (PDB 5SOQ) $IC_{50}$ = 147.9 μM
B.14
Z1211 | 5SOX
RMSD = 6.22 Å
Z1211* (PDB 5SOX) $IC_{50}$ > 1 mM
B.15
Z8186 | 5SP1
RMSD = 6.75 Å
Z8186 (PDB 5SP1) $IC_{50}$ > 1 mM
B.16
Z4827 | 5SP0
RMSD = 3.94 Å
Z4827* (PDB 5SP0) $IC_{50}$ > 1 mM
B.17
Z0893 | 5SOR
RMSD = 3.50 Å
Z0893* (PDB 5SOR) $IC_{50}$ > 1 mM
B.18
Z0078 | 5SOS
RMSD = 1.42 Å
Z0078* (PDB 5SOS) $IC_{50}$ > 1 mM
B.19
Z0828 | 5SOZ
RMSD = 1.59 Å
Z0828 (PDB 5SOZ) $IC_{50}$ > 1 mM
B.20
Z7873 | 5SOJ
RMSD = 1.46 Å
Z7873 (PDB 5SOJ) $IC_{50}$ = 90.3 μM
B.21
Z6923 | 5SP3
RMSD = 1.74 Å
Z6923 (PDB 5SP3) $IC_{50}$ = 416.2 μM
B.22
Z7450 | 5SOY
RMSD = 2.22 Å
Z7450 (PDB 5SOY) $IC_{50}$ > 1 mM
B.23
Z1964 | 5SPJ
RMSD = 1.25 Å
Z1964 (PDB 5SPJ) $IC_{50}$ > 250 μM
B.24
R6134 | 5SPK
RMSD = 8.48 Å
R6134 (PDB 5SPK) $IC_{50}$ > 250 μM
1st Screen Docking Template | 2nd Screen Docking Template
Docking Prediction
Crystal Structure
Solved Pose

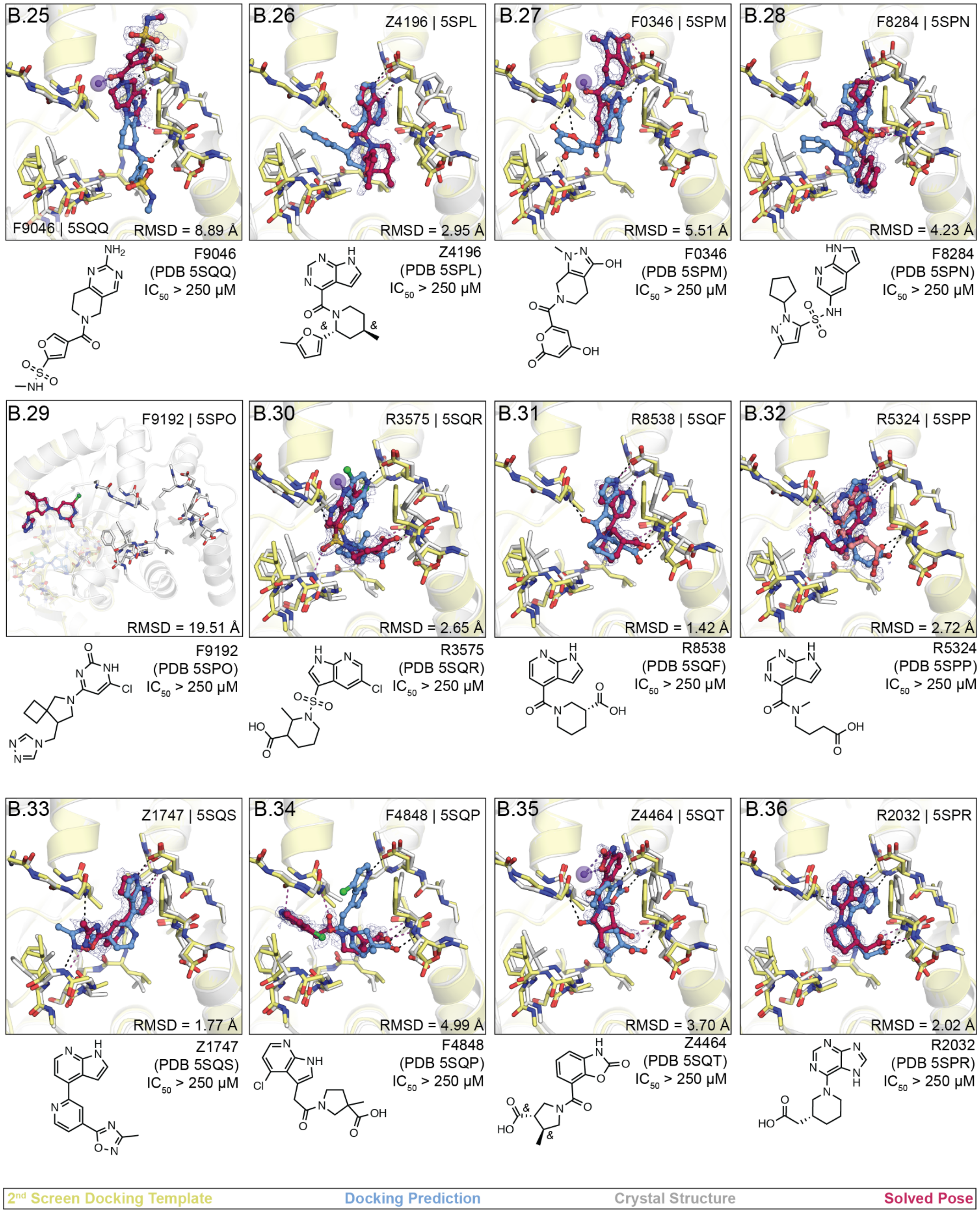
B.25
F9046 | 5SQQ
RMSD = 8.89 Å
F9046
(PDB 5SQQ)
IC50 > 250 µM
B.26
Z4196 | 5SPL
RMSD = 2.95 Å
Z4196
(PDB 5SPL)
IC50 > 250 µM
B.27
F0346 | 5SPM
RMSD = 5.51 Å
F0346
(PDB 5SPM)
IC50 > 250 µM
B.28
F8284 | 5SPN
RMSD = 4.23 Å
F8284
(PDB 5SPN)
IC50 > 250 µM
B.29
F9192 | 5SPO
RMSD = 19.51 Å
F9192
(PDB 5SPO)
IC50 > 250 µM
B.30
R3575 | 5SQR
RMSD = 2.65 Å
R3575
(PDB 5SQR)
IC50 > 250 µM
B.31
R8538 | 5SQF
RMSD = 1.42 Å
R8538
(PDB 5SQF)
IC50 > 250 µM
B.32
R5324 | 5SPP
RMSD = 2.72 Å
R5324
(PDB 5SPP)
IC50 > 250 µM
B.33
Z1747 | 5SQS
RMSD = 1.77 Å
Z1747
(PDB 5SQS)
IC50 > 250 µM
B.34
F4848 | 5SQP
RMSD = 4.99 Å
F4848
(PDB 5SQP)
IC50 > 250 µM
B.35
Z4464 | 5SQT
RMSD = 3.70 Å
Z4464
(PDB 5SQT)
IC50 > 250 µM
B.36
R2032 | 5SPR
RMSD = 2.02 Å
R2032
(PDB 5SPR)
IC50 > 250 µM
2nd Screen Docking Template
Docking Prediction
Crystal Structure
Solved Pose

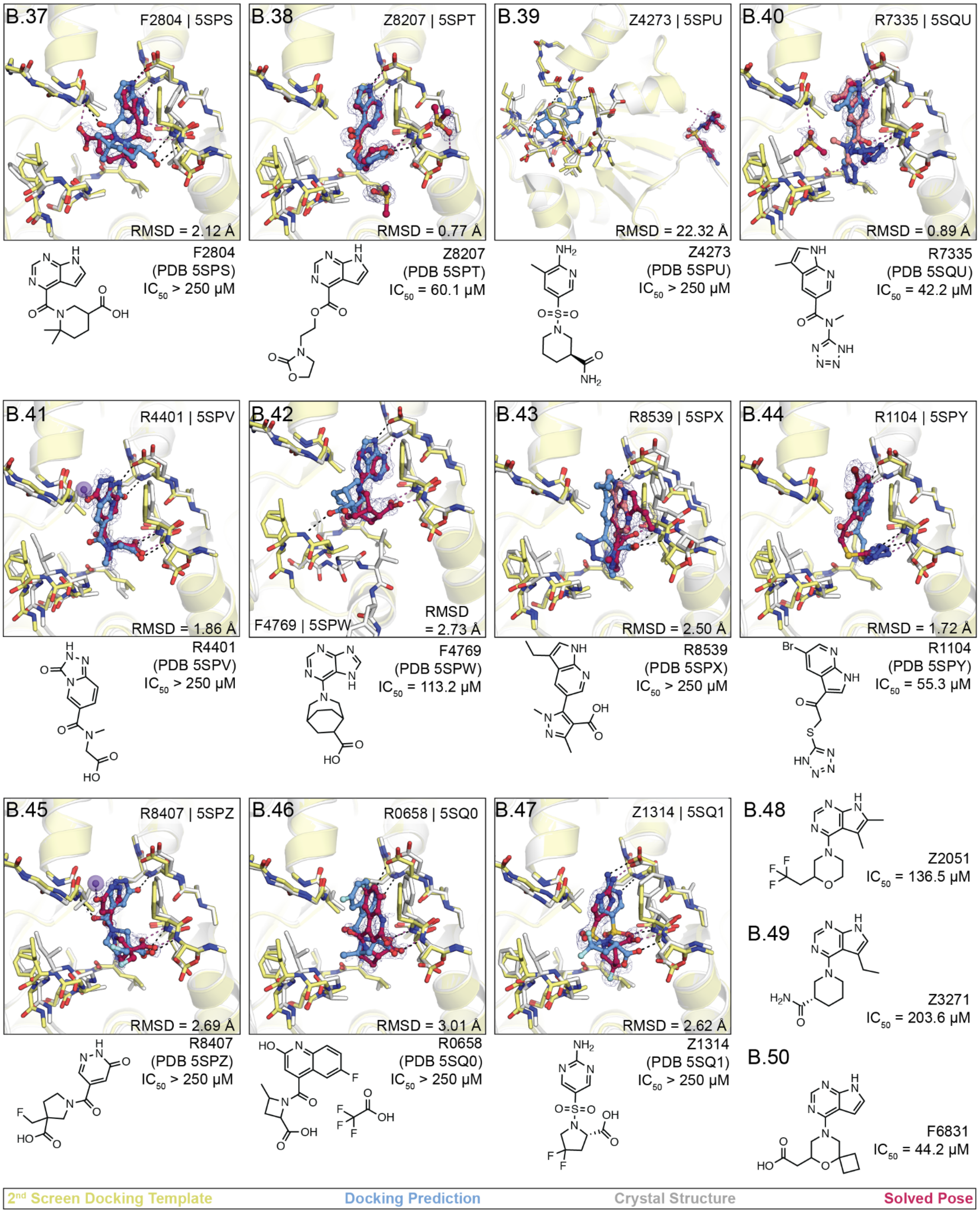
B.37
F2804 | 5SPS
RMSD = 2.12 Å
F2804
(PDB 5SPS)
IC50 > 250 μM
B.38
Z8207 | 5SPT
RMSD = 0.77 Å
Z8207
(PDB 5SPT)
IC50 = 60.1 μM
B.39
Z4273 | 5SPU
RMSD = 22.32 Å
Z4273
(PDB 5SPU)
IC50 > 250 μM
B.40
R7335 | 5SQU
RMSD = 0.89 Å
R7335
(PDB 5SQU)
IC50 = 42.2 μM
B.41
R4401 | 5SPV
RMSD = 1.86 Å
R4401
(PDB 5SPV)
IC50 > 250 μM
B.42
F4769 | 5SPW
RMSD
= 2.73 Å
F4769
(PDB 5SPW)
IC50 = 113.2 μM
B.43
R8539 | 5SPX
RMSD = 2.50 Å
R8539
(PDB 5SPX)
IC50 > 250 μM
B.44
R1104 | 5SPY
RMSD = 1.72 Å
R1104
(PDB 5SPY)
IC50 = 55.3 μM
B.45
R8407 | 5SPZ
RMSD = 2.69 Å
R8407
(PDB 5SPZ)
IC50 > 250 μM
B.46
R0658 | 5SQ0
RMSD = 3.01 Å
R0658
(PDB 5SQ0)
IC50 > 250 μM
B.47
Z1314 | 5SQ1
RMSD = 2.62 Å
Z1314
(PDB 5SQ1)
IC50 > 250 μM
B.48
Z2051
IC50 = 136.5 μM
B.49
Z3271
IC50 = 203.6 μM
B.50
F6831
IC50 = 44.2 μM
2nd Screen Docking Template
Docking Prediction
Crystal Structure
Solved Pose

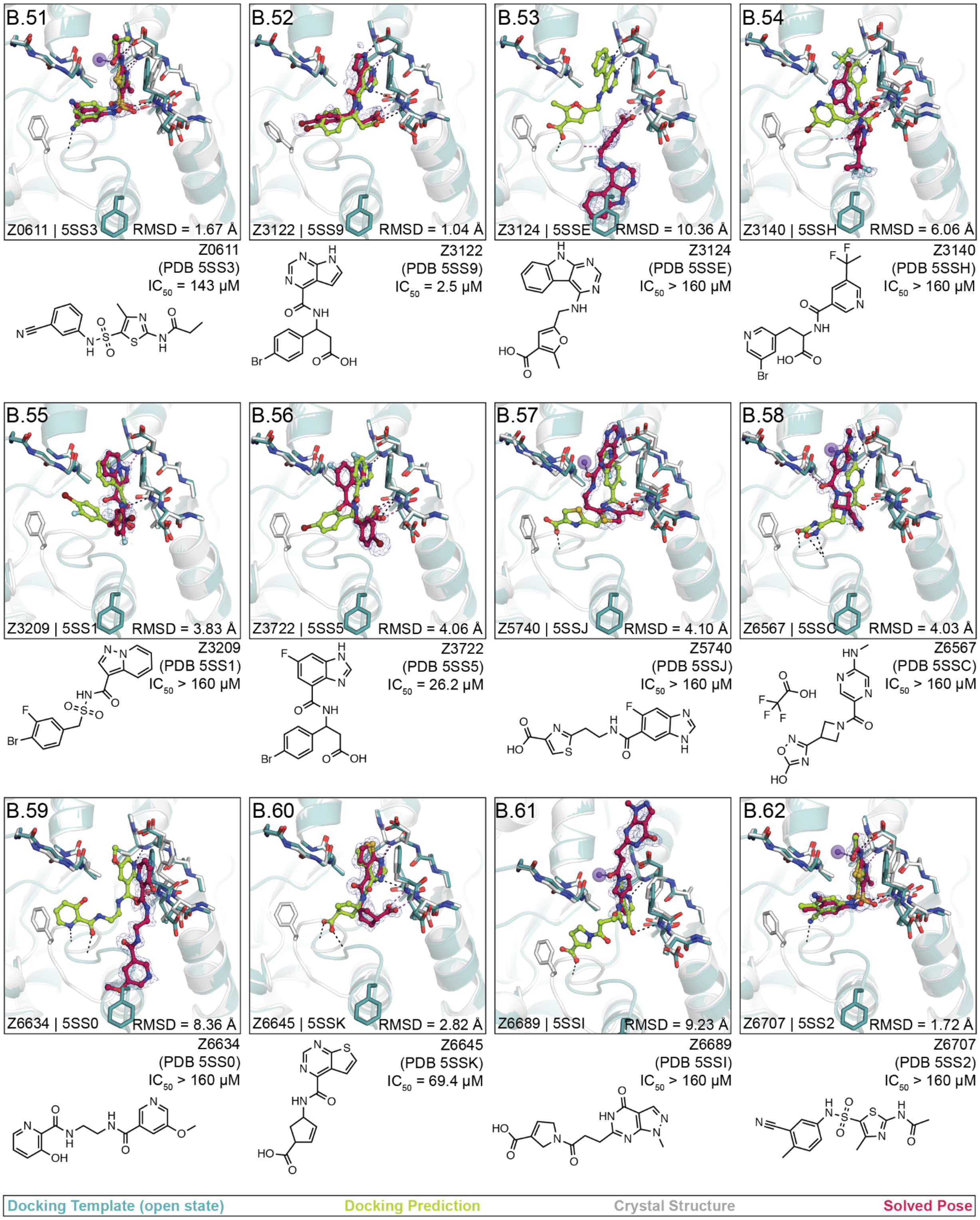

B.51
Z0611 | 5SS3
RMSD = 1.67 Å
Z0611
(PDB 5SS3)
IC50 = 143 μM
B.52
Z3122 | 5SS9
RMSD = 1.04 Å
Z3122
(PDB 5SS9)
IC50 = 2.5 μM
B.53
Z3124 | 5SSE
RMSD = 10.36 Å
Z3124
(PDB 5SSE)
IC50 > 160 μM
B.54
Z3140 | 5SSH
RMSD = 6.06 Å
Z3140
(PDB 5SSH)
IC50 > 160 μM
B.55
Z3209 | 5SS1
RMSD = 3.83 Å
Z3209
(PDB 5SS1)
IC50 > 160 μM
B.56
Z3722 | 5SS5
RMSD = 4.06 Å
Z3722
(PDB 5SS5)
IC50 = 26.2 μM
B.57
Z5740 | 5SSJ
RMSD = 4.10 Å
Z5740
(PDB 5SSJ)
IC50 > 160 μM
B.58
Z6567 | 5SSC
RMSD = 4.03 Å
Z6567
(PDB 5SSC)
IC50 > 160 μM
B.59
Z6634 | 5SS0
RMSD = 8.36 Å
Z6634
(PDB 5SS0)
IC50 > 160 μM
B.60
Z6645 | 5SSK
RMSD = 2.82 Å
Z6645
(PDB 5SSK)
IC50 = 69.4 μM
B.61
Z6689 | 5SSI
RMSD = 9.23 Å
Z6689
(PDB 5SSI)
IC50 > 160 μM
B.62
Z6707 | 5SS2
RMSD = 1.72 Å
Z6707
(PDB 5SS2)
IC50 > 160 μM
Docking Template (open state)
Docking Prediction
Crystal Structure
Solved Pose

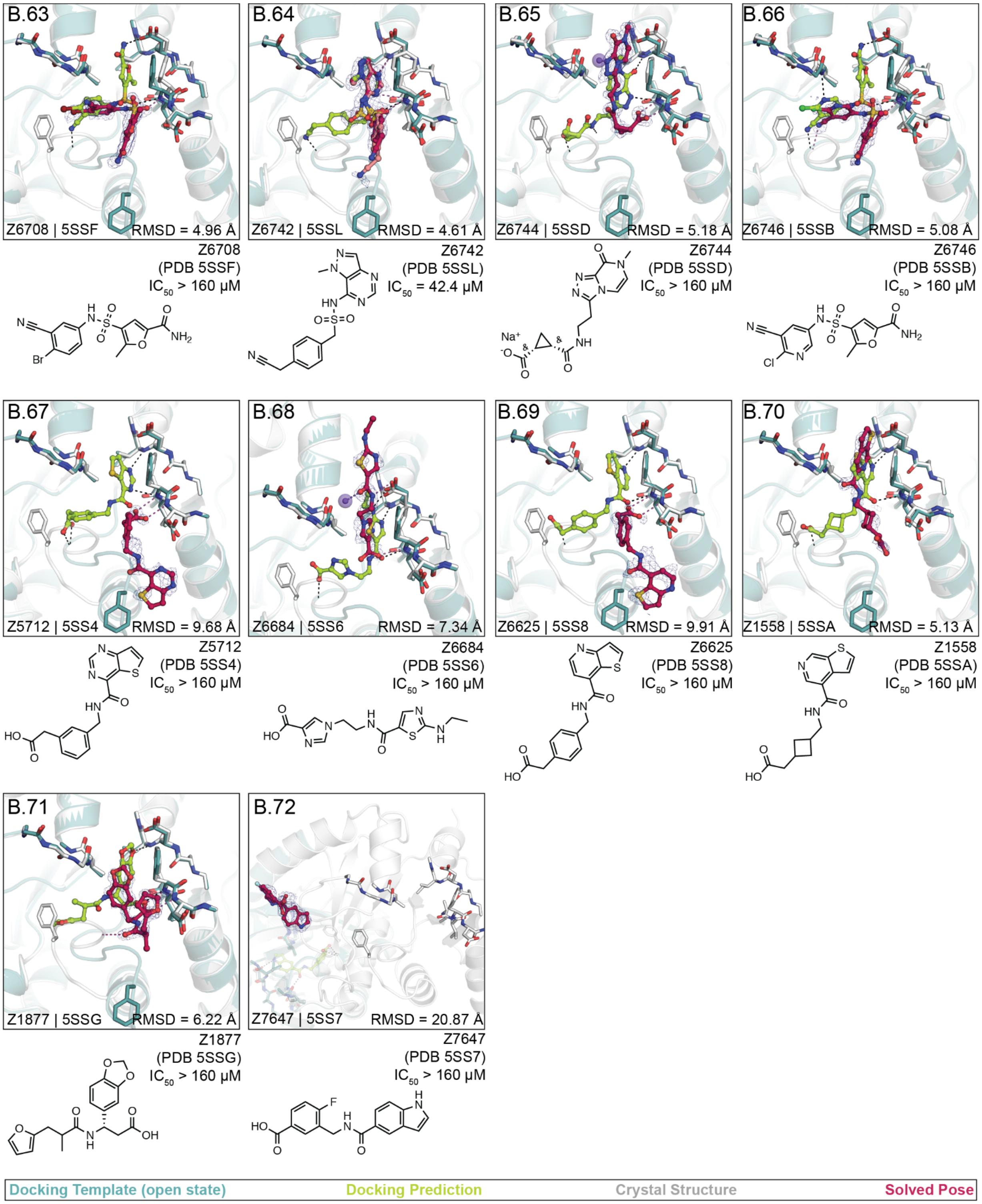
B.63
Z6708 | 5SSF
RMSD = 4.96 Å
Z6708
(PDB 5SSF)
IC50 > 160 µM
B.64
Z6742 | 5SSL
RMSD = 4.61 Å
Z6742
(PDB 5SSL)
IC50 = 42.4 µM
B.65
Z6744 | 5SSD
RMSD = 5.18 Å
Z6744
(PDB 5SSD)
IC50 > 160 µM
Na+
B.66
Z6746 | 5SSB
RMSD = 5.08 Å
Z6746
(PDB 5SSB)
IC50 > 160 µM
B.67
Z5712 | 5SS4
RMSD = 9.68 Å
Z5712
(PDB 5SS4)
IC50 > 160 µM
B.68
Z6684 | 5SS6
RMSD = 7.34 Å
Z6684
(PDB 5SS6)
IC50 > 160 µM
B.69
Z6625 | 5SS8
RMSD = 9.91 Å
Z6625
(PDB 5SS8)
IC50 > 160 µM
B.70
Z1558 | 5SSA
RMSD = 5.13 Å
Z1558
(PDB 5SSA)
IC50 > 160 µM
B.71
Z1877 | 5SSG
RMSD = 6.22 Å
Z1877
(PDB 5SSG)
IC50 > 160 µM
B.72
Z7647 | 5SS7
RMSD = 20.87 Å
Z7647
(PDB 5SS7)
IC50 > 160 µM
Docking Template (open state)
Docking Prediction
Crystal Structure
Solved Pose
