## Supplementary material for "Iterative computational design and crystallographic screening identifies potent inhibitors targeting the Nsp3 Macrodomain of SARS-CoV-2": Dataset S5

**Dataset S5:** Crystal structures of Mac1 in complex with analogs of docking hits (LL1 series). PanDDA event maps are shown for ligands (contoured at 2 σ). Protein-ligand hydrogen bonds are shown with dashed black lines. Hydrogen bonds between ligands and the Lys11 backbone nitrogen of a symmetry mate are highlighted with purple spheres/dashes.

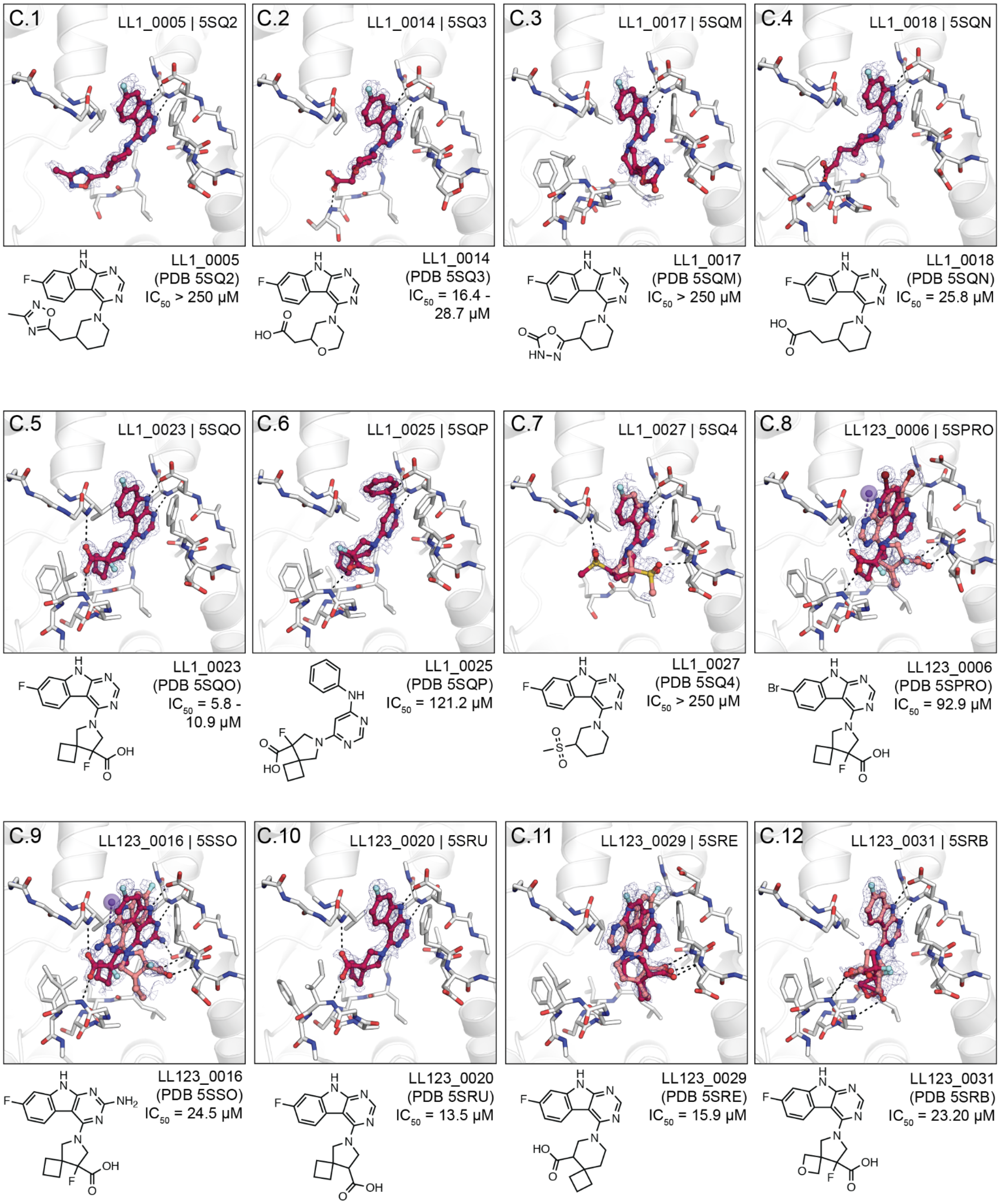

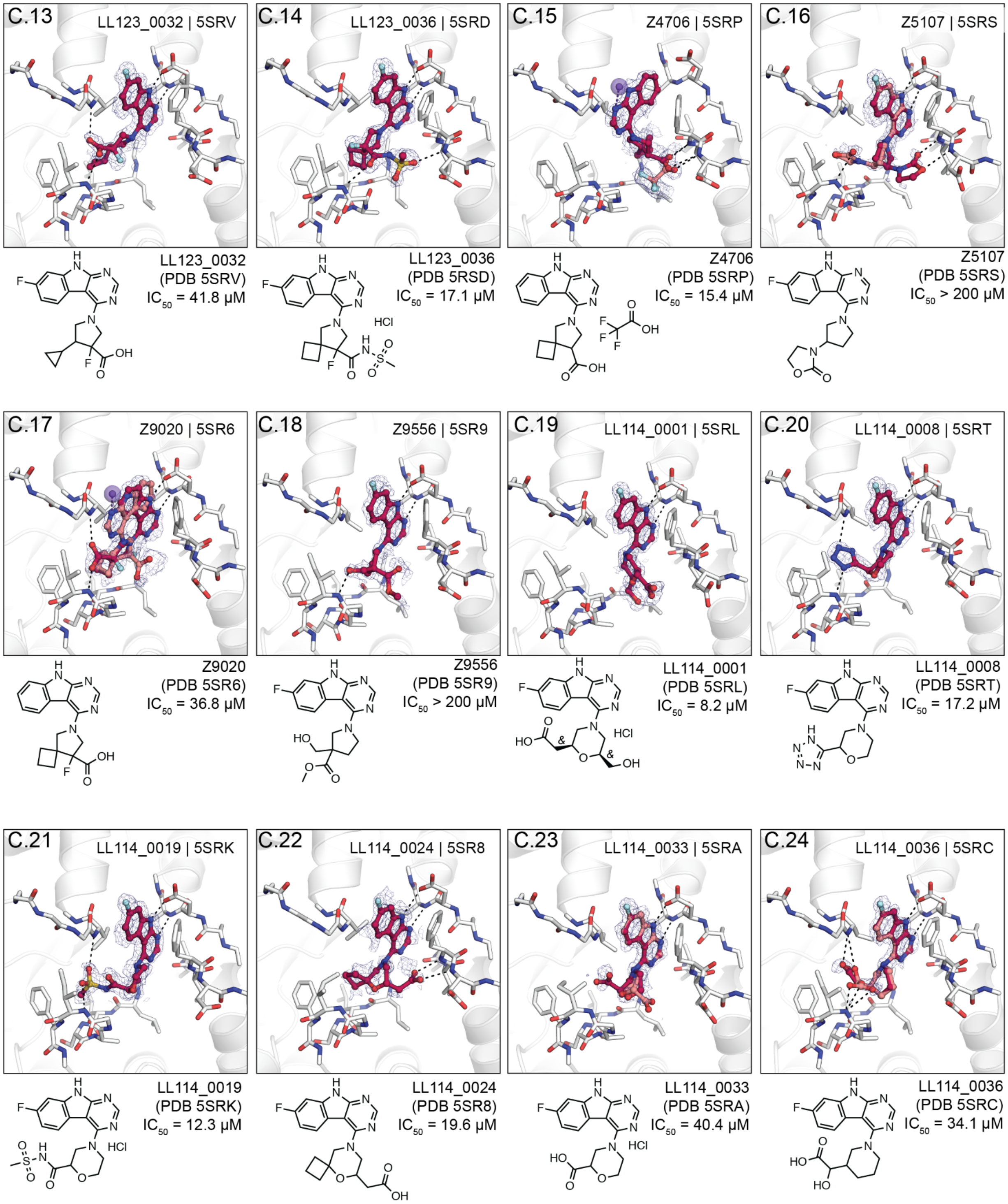

C.13
LL123_0032 | 5SRV
LL123_0032
(PDB 5SRV)
IC50 = 41.8 µM
C.14
LL123_0036 | 5SRD
LL123_0036
(PDB 5RSD)
IC50 = 17.1 µM
HCl
C.15
Z4706 | 5SRP
Z4706
(PDB 5SRP)
IC50 = 15.4 µM
C.16
Z5107 | 5SRS
Z5107
(PDB 5SRS)
IC50 > 200 µM
C.17
Z9020 | 5SR6
Z9020
(PDB 5SR6)
IC50 = 36.8 µM
C.18
Z9556 | 5SR9
Z9556
(PDB 5SR9)
IC50 > 200 µM
C.19
LL114_0001 | 5SRL
LL114_0001
(PDB 5SRL)
IC50 = 8.2 µM
HCl
C.20
LL114_0008 | 5SRT
LL114_0008
(PDB 5SRT)
IC50 = 17.2 µM
C.21
LL114_0019 | 5SRK
LL114_0019
(PDB 5SRK)
IC50 = 12.3 µM
HCl
C.22
LL114_0024 | 5SR8
LL114_0024
(PDB 5SR8)
IC50 = 19.6 µM
C.23
LL114_0033 | 5SRA
LL114_0033
(PDB 5SRA)
IC50 = 40.4 µM
HCl
C.24
LL114_0036 | 5SRC
LL114_0036
(PDB 5SRC)
IC50 = 34.1 µM

C.25

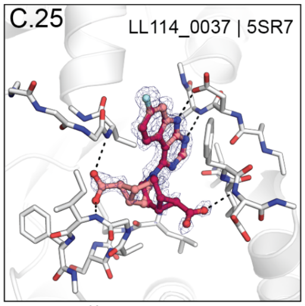


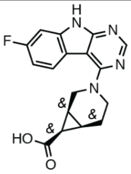

LL114_0037
(PDB 5SR7)
$IC_{50}$ = 37.6 μM

C.26

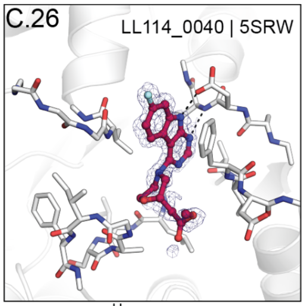


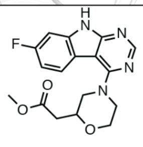

LL114_0040
(PDB 5SRW)
$IC_{50}$ > 200 μM

C.27

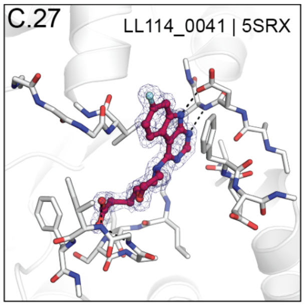


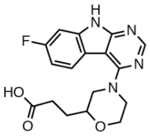

LL114_0041
(PDB 5SRX)
$IC_{50}$ = 69.2 μM

C.28

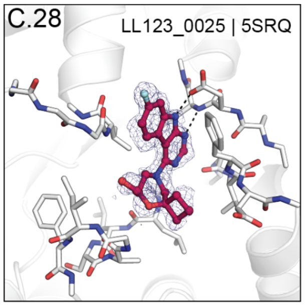


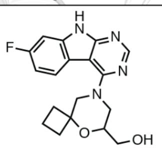

LL123_0025
(PDB 5SRQ)
$IC_{50}$ > 200 μM

C.29

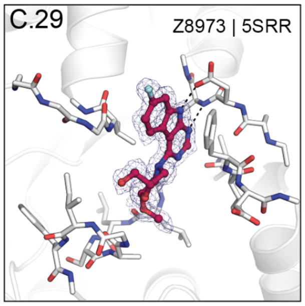


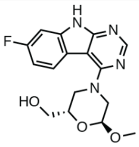

Z8973
(PDB 5SRR)
$IC_{50}$ > 200 μM

C.30

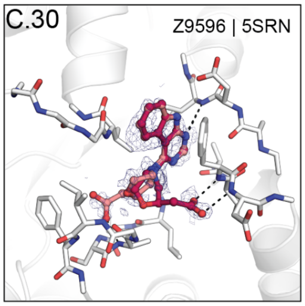


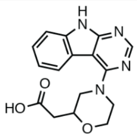

Z9596
(PDB 5SRN)
$IC_{50}$ = 37.1 μM

C.31

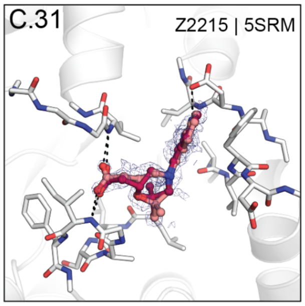


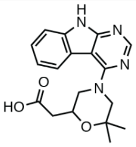

Z2215
(PDB 5SRM)
$IC_{50}$ = 116 μM

C.32

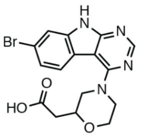

LL114_0014

$IC_{50}$ = 25.9 μM

C.33

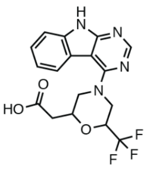

Z1229

$IC_{50}$ = 119.3 μM

C.34

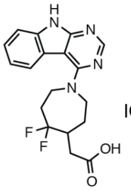

Z6423

$IC_{50}$ = 142.9 μM
