## Supplementary material for "Iterative computational design and crystallographic screening identifies potent inhibitors targeting the Nsp3 Macrodomain of SARS-CoV-2": Dataset S6

**Dataset S6:** Design of neutral probe set. Two carboxylic acids were included as close analogs. Crystal structures of Mac1 in complex with neutral probe set hits. PanDDA event maps are shown for ligands (contoured at 2 σ). Protein-ligand hydrogen bonds are shown with dashed black lines. Hydrogen bonds between ligands and the Lys11 backbone nitrogen of a symmetry mate are highlighted with purple spheres/dashes.

D.1

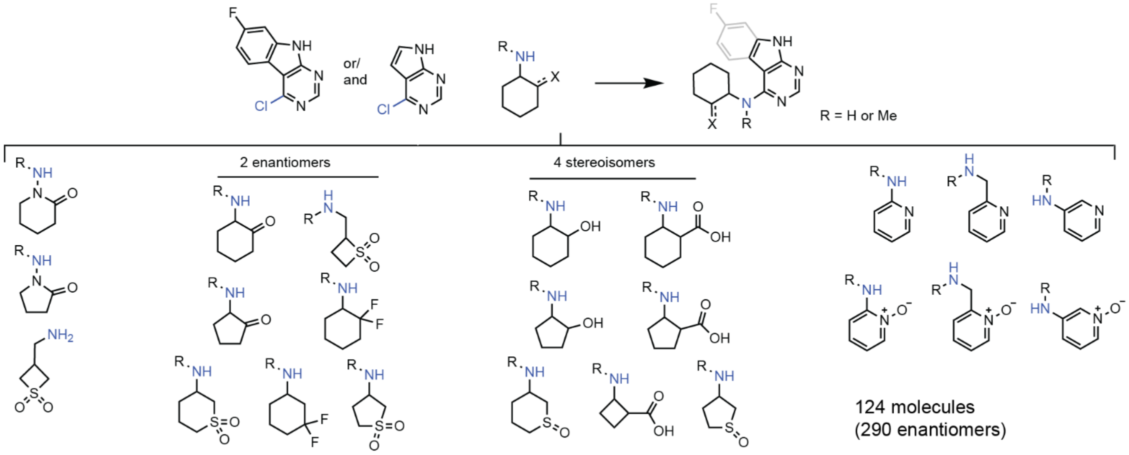


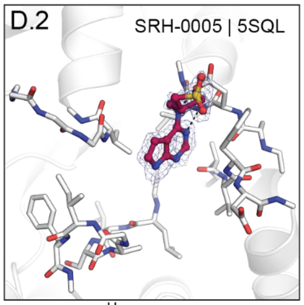


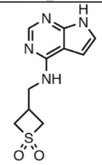

SRH-0005
(PDB 5SQL)
$IC_{50}$ > 200 µM

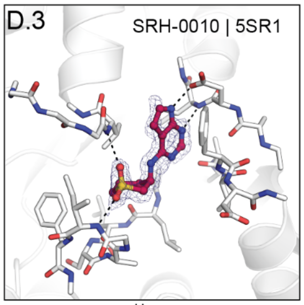


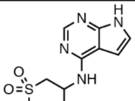

SRH-0010
(PDB 5SR1)
$IC_{50}$ > 200 µM

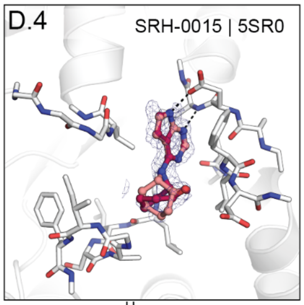


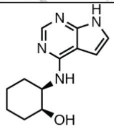

SRH-0015
(PDB 5SR0)
$IC_{50}$ = 131.6 µM

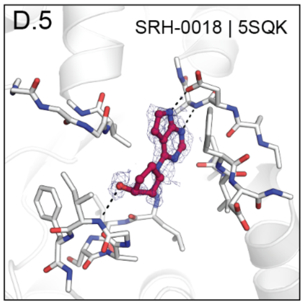


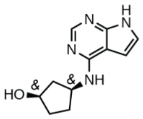

SRH-0018
(PDB 5SQK)
$IC_{50}$ > 200 µM

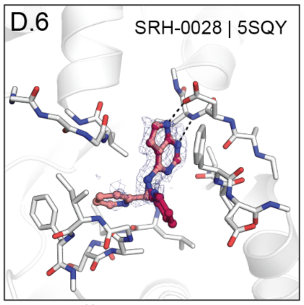


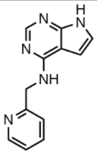

SRH-0028
(PDB 5SQY)
$IC_{50}$ > 200 µM

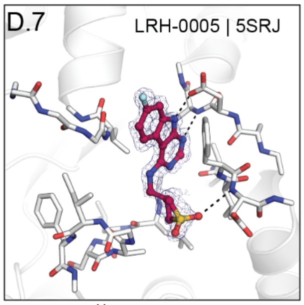


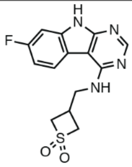

LRH-0005
(PDB 5SRJ)
$IC_{50}$ > 200 µM

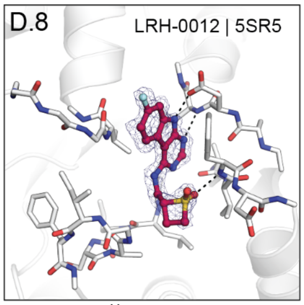


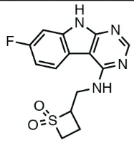

LRH-0012
(PDB 5SR5)
$IC_{50}$ > 200 µM

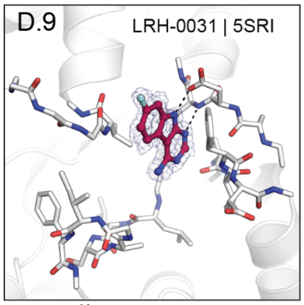


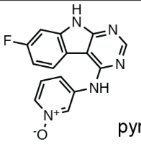

LRH-0031
(PDB 5SRI)
$IC_{50}$ > 200 µM

no desnity for
pyridin-1-ium-1-olate

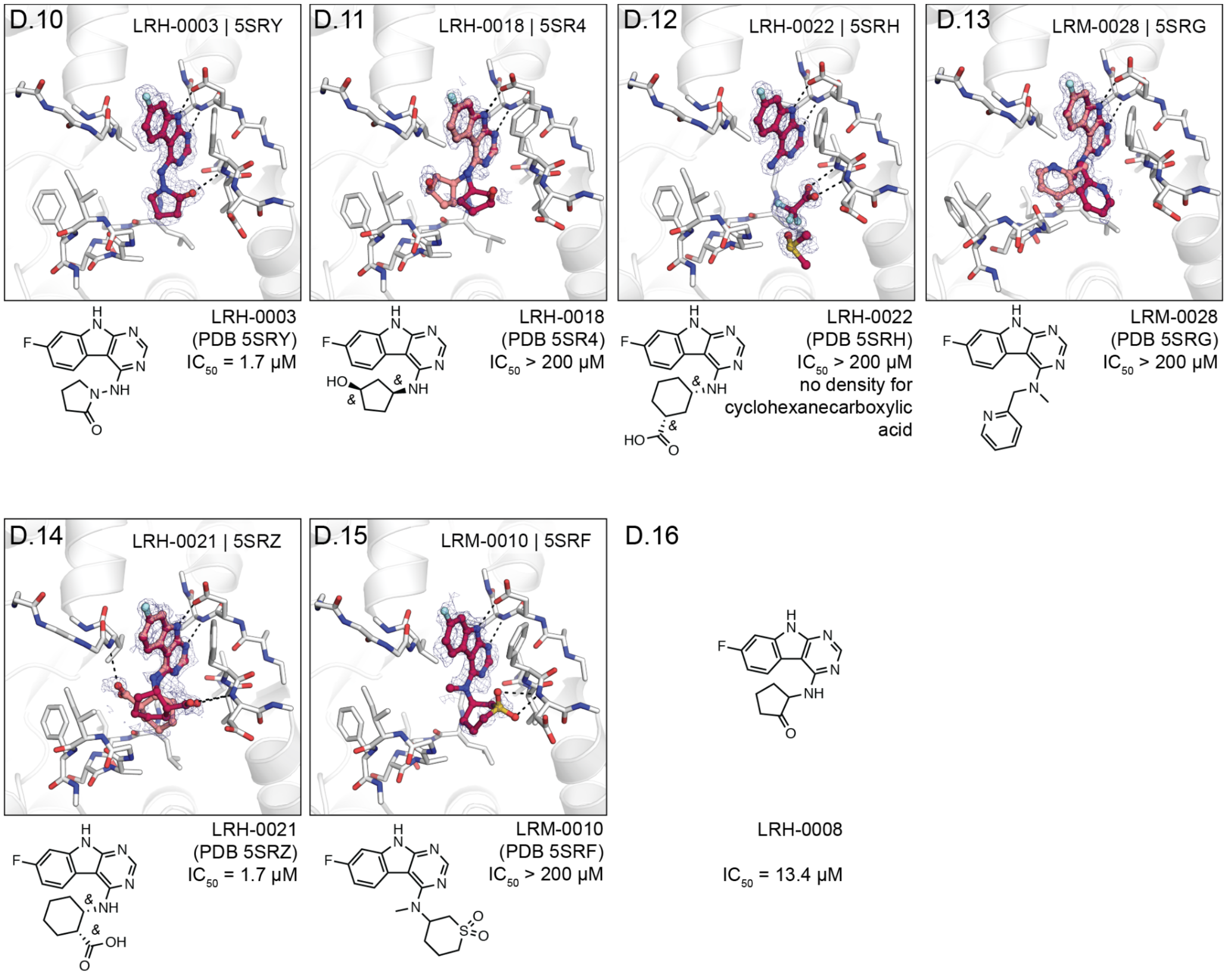
D.10
LRH-0003 | 5SRY
LRH-0003
(PDB 5SRY)
IC50 = 1.7 µM
D.11
LRH-0018 | 5SR4
LRH-0018
(PDB 5SR4)
IC50 > 200 µM
D.12
LRH-0022 | 5SRH
LRH-0022
(PDB 5SRH)
IC50 > 200 µM
no density for
cyclohexanecarboxylic
acid
D.13
LRM-0028 | 5SRG
LRM-0028
(PDB 5SRG)
IC50 > 200 µM
D.14
LRH-0021 | 5SRZ
LRH-0021
(PDB 5SRZ)
IC50 = 1.7 µM
D.15
LRM-0010 | 5SRF
LRM-0010
(PDB 5SRF)
IC50 > 200 µM
D.16
LRH-0008
IC50 = 13.4 µM
